# A hierarchical Bayesian model reveals increased precision weighting for afferent cardiac signals, and reduced anxiety, as a function of interoceptive training

**DOI:** 10.1101/2024.09.26.614928

**Authors:** Chatrin Suksasilp, Abigail McLanachan, Lisa Quadt, Blaise Boulton, James Mulcahy, Hugo D Critchley, Ryan Smith, Sarah N Garfinkel

## Abstract

Perceptual accuracy for interoceptive signals, such as heartbeats, varies in a trait-like manner across individuals and may influence the capacity for emotion regulation and vulnerability to affective symptoms, notably anxiety. Here, we demonstrate that an interoceptive training protocol improved perceptual accuracy in two tasks of heartbeat perception and reduced both state and trait anxiety in a subclinical sample, extending previous findings in autistic adults. Computational modelling indicated that accuracy improvement in the heartbeat discrimination task was associated with increases in the internal reliability estimate for interoceptive signals – their precision weighting – while a lower-level parameter representing noise in the interoceptive signal itself (which influences speed of learning) moderated this precision weighting improvement. Reductions in both state and trait anxiety in the training group were uniquely explained by computational parameter estimates, and not by conventional accuracy measures. These findings indicate that trait-like differences in interoceptive processing are modifiable and can be targeted to alleviate anxiety symptoms, and that interoceptive interventions may be best guided by a computational phenotyping approach.

## Introduction

Interoception refers to the afferent peripheral signalling, central neural processing, and mental representation of internal bodily changes. These processes inform homeostatic control (Pezzulo et al., 2015) and induce momentary changes in a variety of cognitive and emotional processes (Ashhad et al., 2022; Critchley & Garfinkel, 2017). There is a growing body of evidence for altered interoceptive processing across a range of clinical populations (Khalsa et al., 2018; Quadt et al., 2018), including anxiety disorders (Domschke et al., 2010; Ehlers & Breuer, 1992), eating disorders (Pollatos et al., 2008; Pollatos & Georgiou, 2016), and depression (Avery et al., 2014; Dunn et al., 2007).

When instructed to attend to bodily signals in behavioural tasks, healthy individuals vary in their ability to accurately perceive those signals (Katkin et al., 1982; Koch & Pollatos, 2014). Further, trait-like differences in both objective and self-reported interoceptive accuracy have been linked to the perceived intensity of affective stimuli (Schandry, 1981; Wiens et al., 2000), difficulties understanding one’s own emotions (alexithymia; Brewer et al., 2016; Trevisan et al., 2019), and the capacity to regulate one’s own emotional states (Edwards & Pinna, 2020; Zamariola et al., 2019).

Theoretical work has proposed that anxiety arises, in part, from dysfunction in processes of Bayesian perception that are thought to underpin interoception. Specifically, individuals may develop anxious symptoms when confronted with chronic interoceptive prediction errors, or discrepancies between expected and observed bodily signals (Khalsa & Feinstein, 2018; Paulus & Stein, 2006). Chronic interoceptive prediction errors are proposed to arise either from (a) chronically noisy or imprecise afferent interoceptive signals; or (b) from the brain inappropriately treating afferent signals as unreliable by maintaining low internal (sub-personal) estimates of their precision – referred to as precision weighting – and/or (c) the brain generating maladaptive prior expectations about interoceptive states (Paulus et al., 2019; Paulus & Stein, 2006). Correspondingly, ‘normalising’ the precision weighting afforded to interoceptive signals has been proposed as a potential means of improving affective symptomatology (Owens et al., 2018). Conventional measures of interoceptive accuracy, assessed using behavioural tasks, are also thought to derive from interoceptive precision weighting (Ainley et al., 2016), which therefore represents a mechanistic computational target for interoceptive interventions that aim to decrease anxiety.

The present study therefore aimed to test the utility of an intervention designed to improve cardiac interoceptive accuracy and ameliorate anxiety symptoms in a subclinical sample, following a previous report of successful results in a randomised controlled trial in autistic adults (Quadt et al., 2021). Participants in the training group completed tasks of cardiac perception with feedback over eight sessions across five weeks, and were compared against a passive control group. We hypothesised that the training intervention would reduce both trait and state anxiety, while improving cardiac interoceptive accuracy.

This study also aimed to clarify the mechanisms underpinning interoceptive training and test whether these mechanisms could explain individual differences in anxiety reduction and interoceptive changes. To do so, we fit Bayesian computational models to participant responses during a cardiac perception task, using a ‘computational phenotyping’ approach that can characterise individual differences, not just in terms of task responses, but in the belief updating mechanisms underlying those responses under ideal Bayesian observer assumptions (Schwartenbeck & Friston, 2016). This study extends previous computational models of interoception (Lavalley et al., 2023; Smith, Kuplicki, Feinstein, Forthman, Stewart, Paulus, Tulsa 1000 investigators, et al., 2020; Smith, Kuplicki, Teed, et al., 2020; Smith, Mayeli, et al., 2021) by implementing the bottom-up effects of interoceptive signals within a hierarchical Bayesian model, applying it to the heartbeat discrimination task in a novel implementation (i.e., the previous models were confined to a heartbeat tapping task), and testing a large number of competing hypotheses for the mechanisms of interoceptive learning.

We hypothesised that training-based improvements in cardiac interoceptive accuracy would correspond to increases in the precision weighting assigned to cardiac signals, and that these improvements would be moderated by a computational measure of baseline noise in the interoceptive signal that can influence speed of learning. Furthermore, we predicted that increases in the precision weighting assigned to cardiac signals due to training would be associated with anxiety reduction, in line with the hypothesis that ‘normalising’ interoceptive precision weighting should reduce anxiety.

## Methods

### Participants

Participants included staff and students recruited from the University of Sussex and through adverts placed around the local community in Brighton and Hove. A total of 54 participants took part in the study; 28 (20 F) were assigned to the interoceptive training group and 26 (22 F) to the control group. The mean age was 27.9 yrs in the training group (range 18 – 48 yrs) and 25.5 yrs in the control group (range 19 – 45 yrs). Ethical approval for the study was granted by the Research Ethics and Governance Committee (School of Psychology) at the University of Sussex. All participants gave informed consent after being provided with written details of the experiment.

### Interoceptive training

Participants in the training group completed eight training sessions within a five-week period, resulting in 1-3 training sessions per week. Each training session comprised two blocks of heartbeat perception tasks modified to incorporate feedback. In each block, participants completed six trials of the heartbeat counting task (Brener & Kluvitse, 1988; Whitehead et al., 1977), followed by twenty trials of the heartbeat discrimination task (Schandry, 1981).

In the heartbeat discrimination task, participants judged the synchronicity of sets of ten tones, relative to their own heartbeat. They were given the instruction: ‘*You will hear ten tones. Please tell me if the tones are in sync or out of sync with your own heartbeat*’. Within each trial, the tones were presented at 440 Hz for 100 ms and triggered by the participant’s own consecutive heartbeats. Synchronous tones were presented at the beginning of the rising edge of the pulse pressure wave and asynchronous tones were presented after a delay of 300 ms, adjusting for the average delay (∼250 ms) between the R-wave and the arrival of the pressure wave at the finger (Payne et al., 2006). At the end of each trial, participants reported whether the tones were synchronous or asynchronous with their own heartbeats and then received feedback about whether their response was correct or incorrect. Each block of the discrimination task included ten trials in the synchronous condition and ten trials in the asynchronous condition presented in random order.

In the heartbeat counting task, participants were instructed: ‘*Without manually checking, can you silently count each heartbeat you feel in your body from the time you hear “start” to when you hear “stop*”’. In each block of this task, participants completed six trials, across randomised time-windows of 25, 30, 35, 40, 45 and 50 s. The number of heartbeats counted was recorded after each trial, and participants were given accurate feedback about the true number of heartbeats that occurred.

In between training task blocks, participants were instructed to engage in a self-paced low-level physical activity for 1-2 minutes, to the point where their heartrate became noticeably elevated, but to stop before discomfort occurred. Suggested methods were star jumps or jogging on the spot, but other methods were accepted so long as participants reported feeling an elevated heart rate. The physical activity was always performed prior to the second block of tasks, to minimise the time taken for each training session, because performing physical activity prior to the first block would have necessitated a resting period between blocks to prevent cardiovascular arousal from the first ‘physically active’ block contaminating performance in the subsequent ‘resting’ block.

All tasks were programmed in Matlab GUIDE v2.5 running under MATLAB R2012a (The MathWorks, Inc., Natick, MA), while heartbeats were monitored using medical grade pulse oximeters (Nonin 8600 with a ‘soft’ sensor fitting to reduce exteroceptive feedback). Heartrate during each trial of the heartbeat discrimination task was recorded, and averaged within each training session, across all training sessions, and within each assessment session. Due to technical error, six participants in the training group were presented a slightly greater proportion of synchronous trials than asynchronous trials in the heartbeat discrimination task during training sessions, ranging between 53% to 63% synchronous trials, rather than the intended 50% (however, it should be noted that this difference is naturally accounted for when fitting computational models to this data).

#### Assessment sessions

Both the training and control groups completed baseline, mid-point, and final assessment sessions (**Figure 1**). All participants completed both the heartbeat counting and heartbeat discrimination tasks during each assessment to measure interoceptive accuracy. Importantly, participants were not given feedback on their responses during assessment tasks. In all three assessments, the heartbeat tracking task was always performed first so as not to prime participants with immediate temporal cues regarding their own heart rate. State anxiety and trait anxiety, along with self-reported interoception, were also measured at baseline and final assessment sessions.

**Figure 1.**
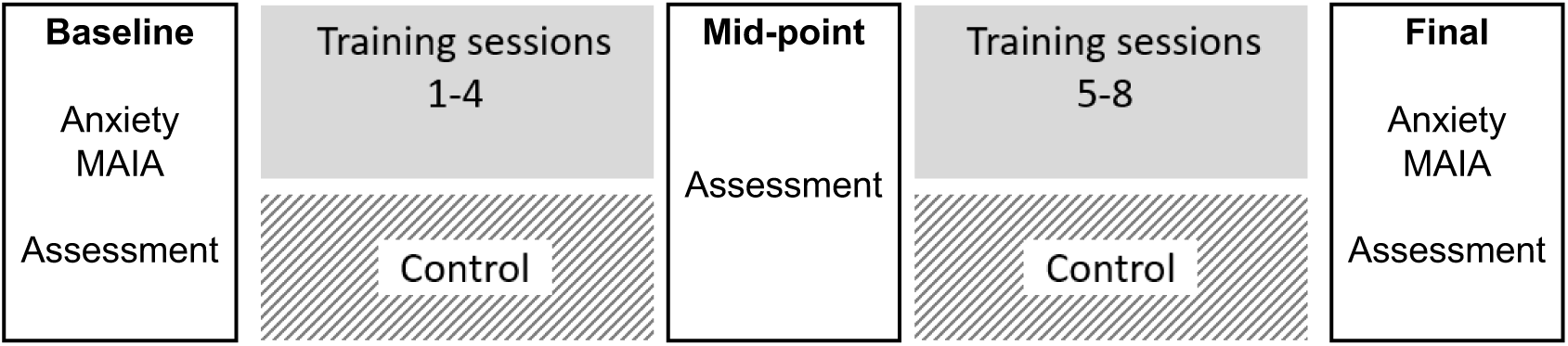
Illustration of procedure for interoceptive training and (passive) control groups. MAIA: Multidimensional Assessment of Interoceptive Awareness.

Due to technical error, participants in the control group completed 20 heartbeat discrimination trials during baseline, mid-point, and final sessions, while individuals in the training group completed 26 trials in each assessment.

### Computational Modelling

To test competing hypotheses concerning the mechanisms underpinning interoceptive learning, several computational models were fit to participant responses on the heartbeat discrimination task to estimate each participant’s prior beliefs, evidence accumulation rate, and interoceptive sensory precision(s), among other parameters in extended models (described below). **Figure 2** provides a graphical depiction and explanation of the model structure used for subsequent between-subject analyses, and its associated vectors and matrices.

**Figure 2.**
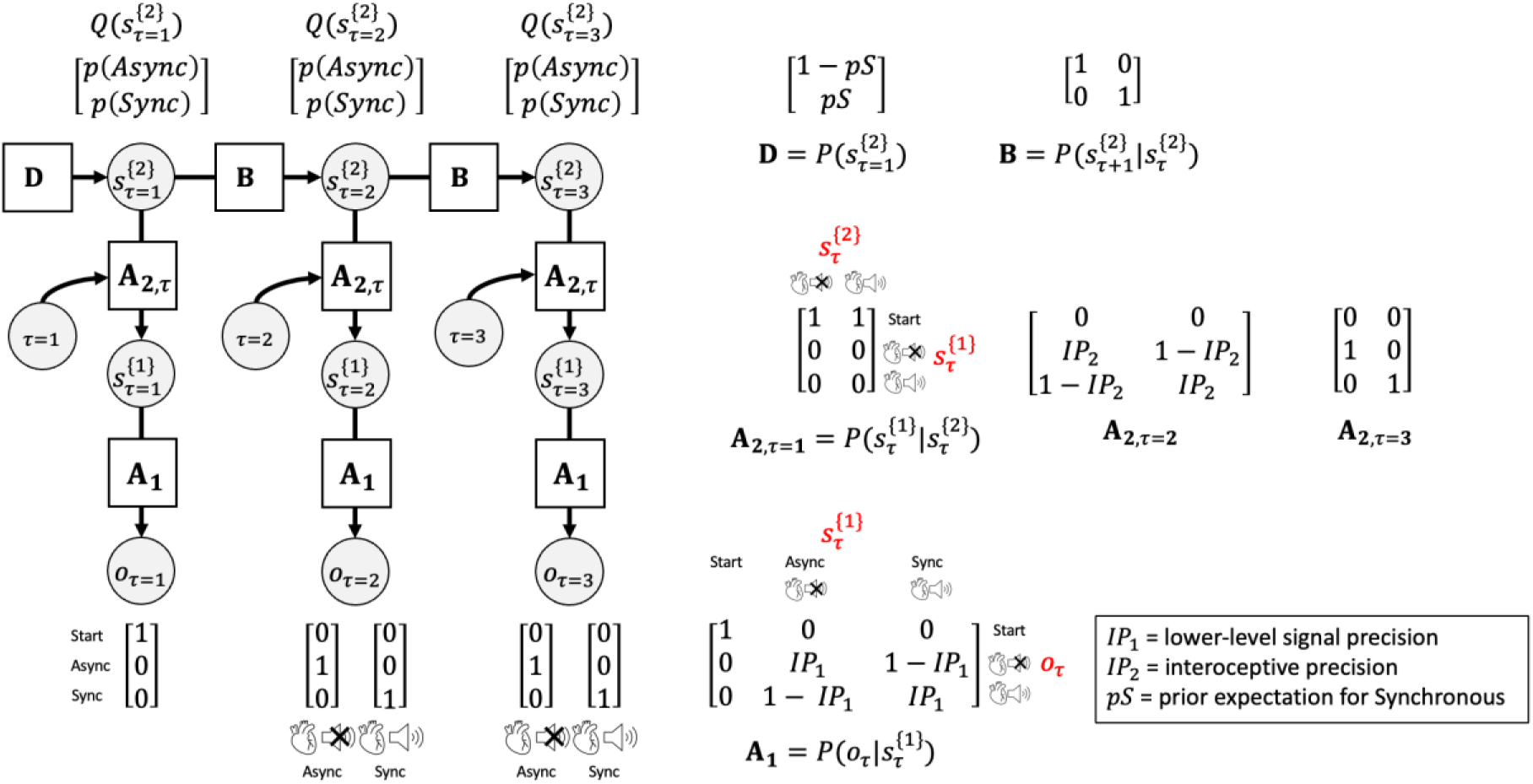
Bayesian approach used to model cardiac perception on the heartbeat discrimination task, assuming two hierarchical levels of inference. The generative model is here depicted graphically, such that arrows indicate dependencies between variables. Associated vectors/matrices are also shown. This hidden Markov model was adapted from the commonly used active inference formulation of partially observable Markov decision processes (Da Costa et al., 2020; Smith, Friston, et al., 2021). Each trial in the heartbeat discrimination task corresponded to a trial in the model, and was divided into three timepoints (*τ*): *τ* = 1 was a placeholder ‘start’ timepoint, while at *τ* = 2 participants listened to the auditory tones and gave responses ‘in sync’ or ‘out of sync’. At *τ* = 3, participants were informed whether their response was correct or incorrect. In a hierarchical model, observations at each timepoint (*o*_*τ*_) depend on lower-level hidden states 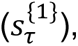 which in turn depend on higher-level hidden states 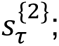 these relationships are specified in the **A_1_** and **A**_**2**,***τ***_ matrices, respectively. The initial higher-level hidden state depends on the probabilities specified in the vector **D**, while successive hidden states depend on the transition probabilities 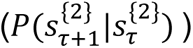 specified in the (identity) matrix **B**. In this model, observations corresponded to the (ground-truth) cardiac-auditory sensory signals received during a trial of the heartbeat discrimination task, which could be either synchronous or asynchronous. The higher-level hidden states corresponded to participants’ beliefs (i.e., posterior probability distributions) about whether the cardiac-auditory sensory signals were synchronous or asynchronous, and participants’ responses were assumed to be sampled from these beliefs. On each trial, the participant begins at time *τ* = 1 and receives a placeholder ‘start’ observation (*o*_*τ*=1_) regardless of the hidden state (asynchronous or synchronous trial condition), and then updates their probabilistic beliefs about hidden states (*Q*(*s*_*τ*_)) based on observations received at timepoint 2 (*o*_*τ*=2_) and timepoint 3 (*o*_*τ*=3_). Belief updating was assumed to rely on Bayesian inference, as implemented in the “Heartbeat-tone perception” equation (see **Table 1**). Learning (evidence accumulation and possible forgetting) was hypothesised to occur in the **A**_**2**,***τ***_ matrix and was controlled by the learning equation (also see **Table 1**). [Note that non-hierarchical models were also considered, in which matrix **A**_**1**_ and the lower-level hidden state 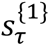 were not present. In these non-hierarchical models, observations *o*_*t*_ depend directly on hidden states 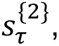 with this relationship specified in matrix **A**_**2**,***τ***_.]

**Table 1.**
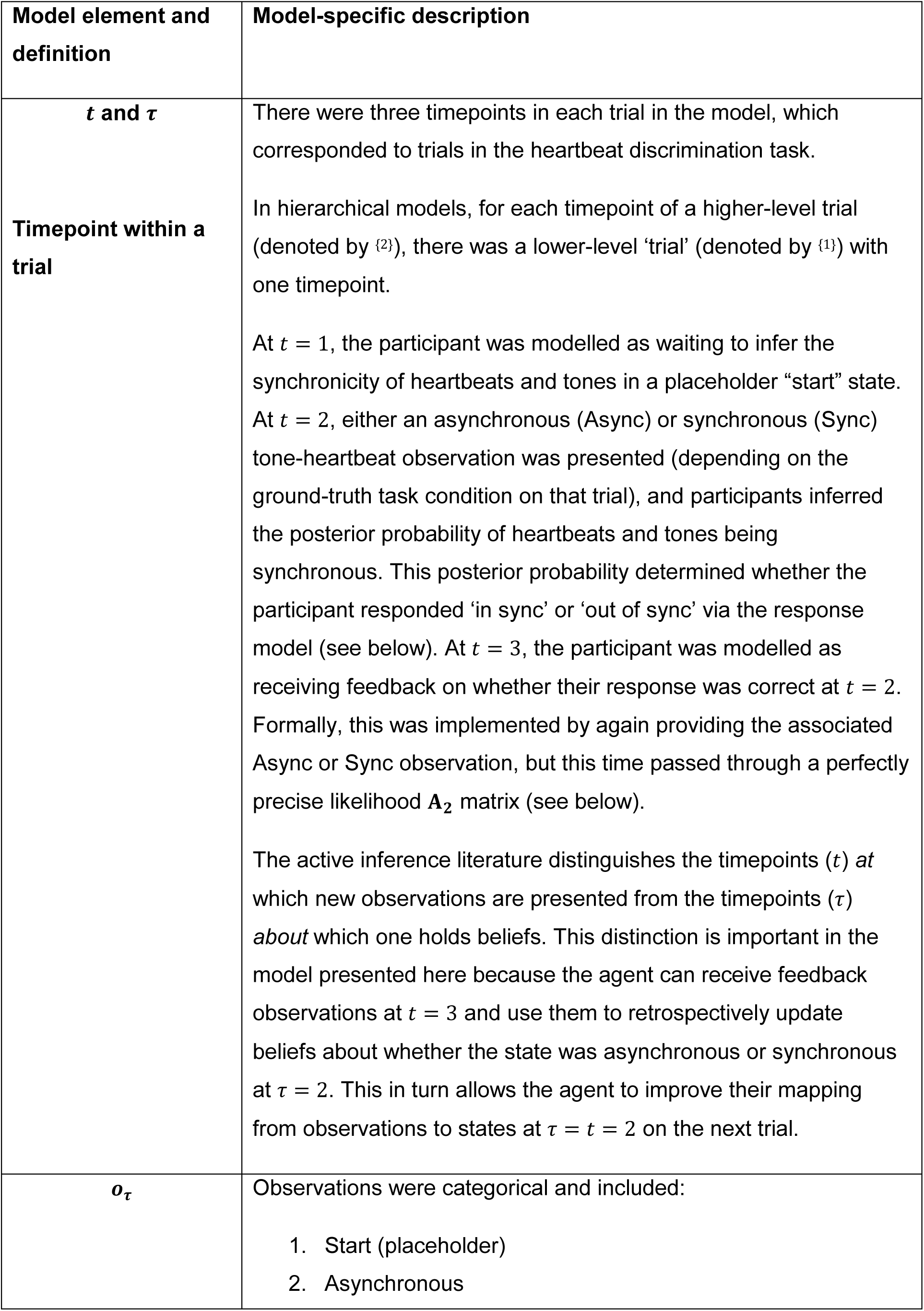
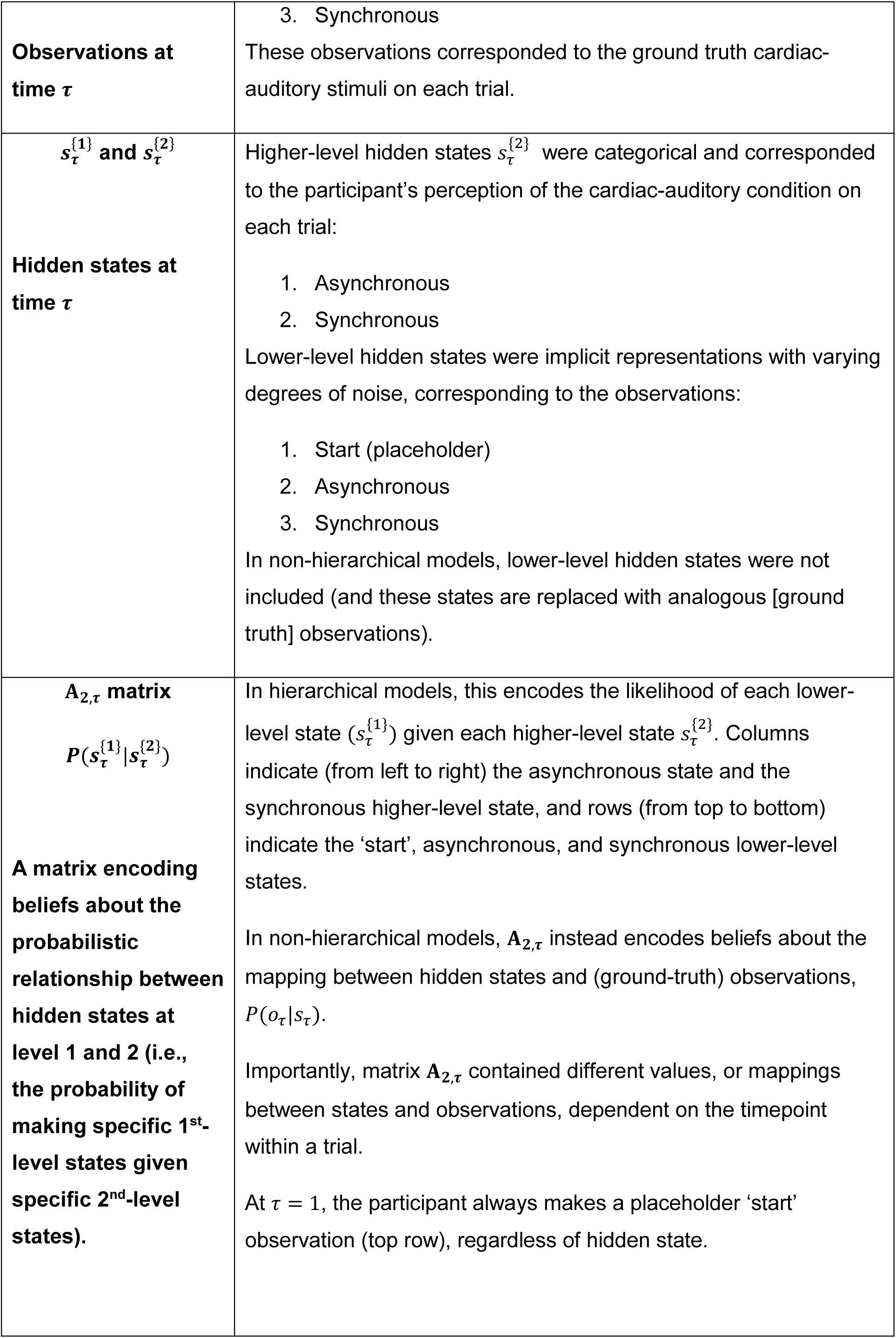
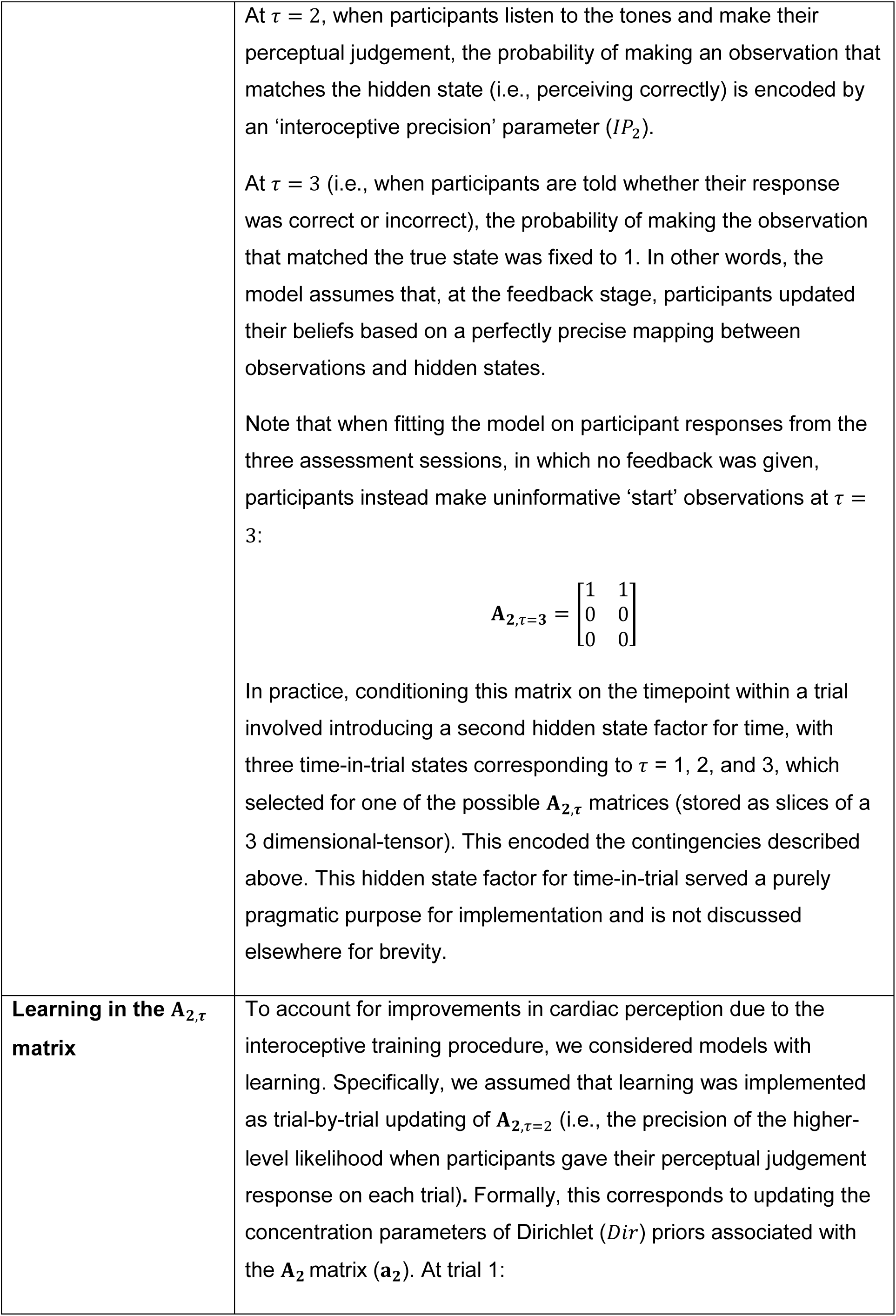
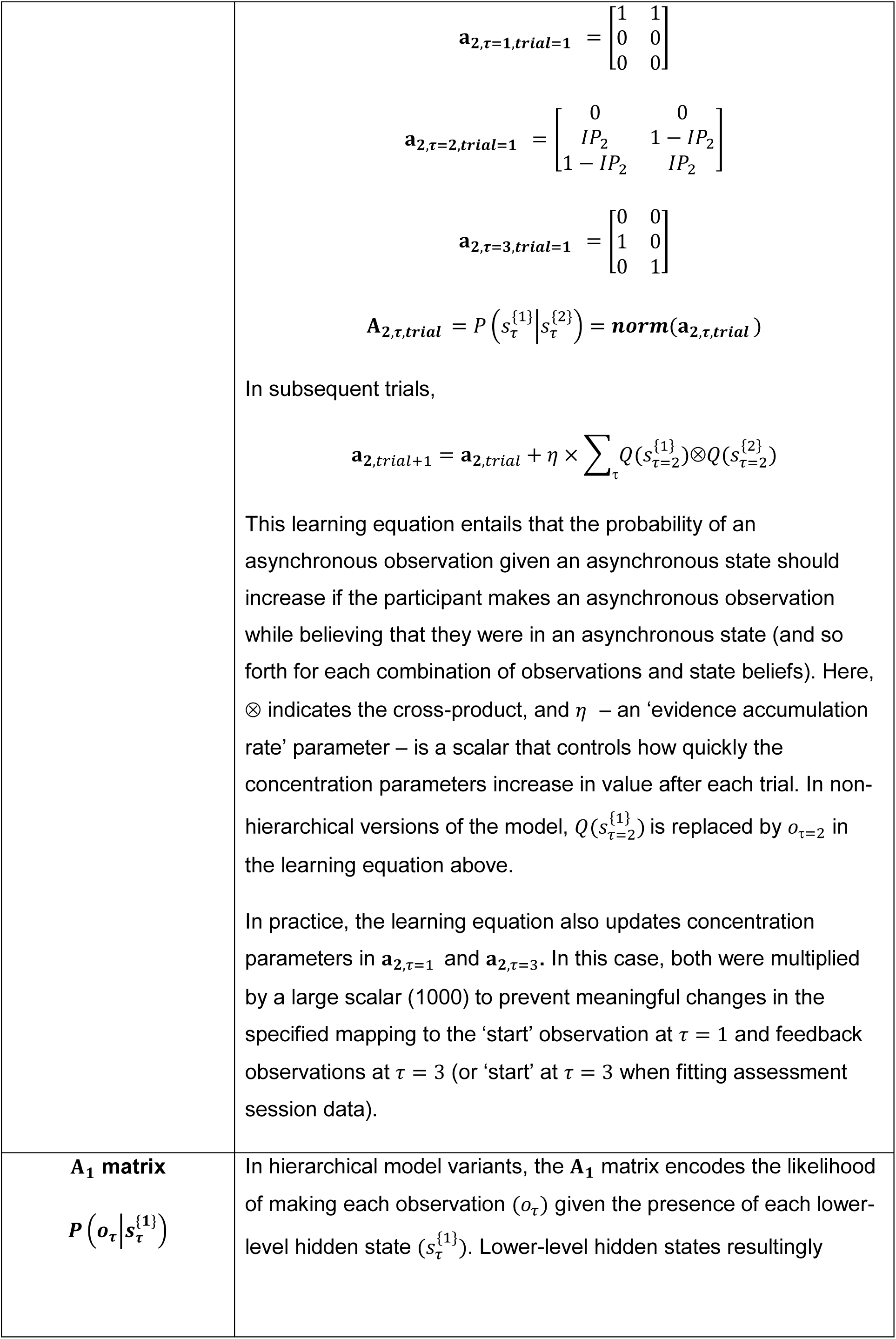
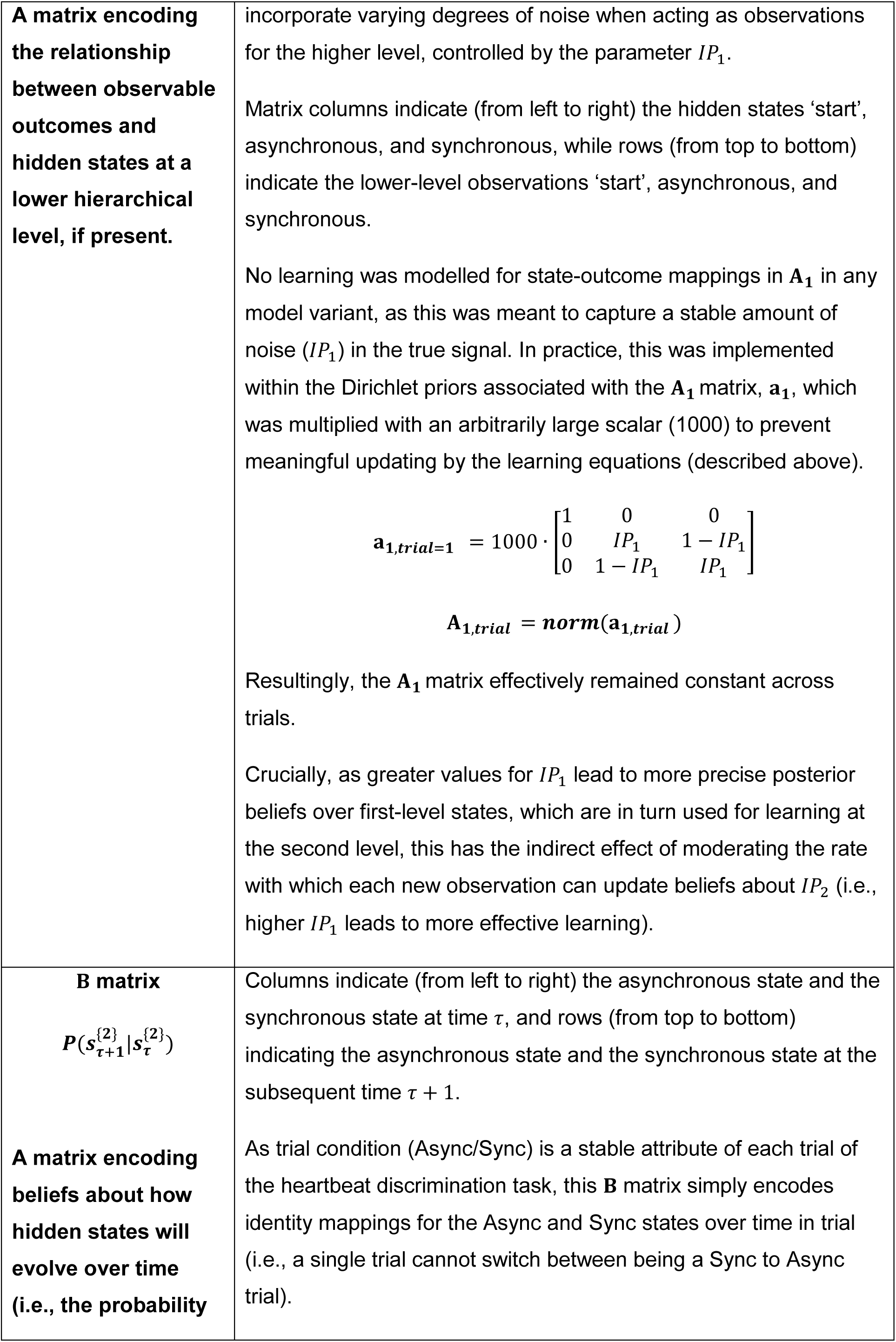
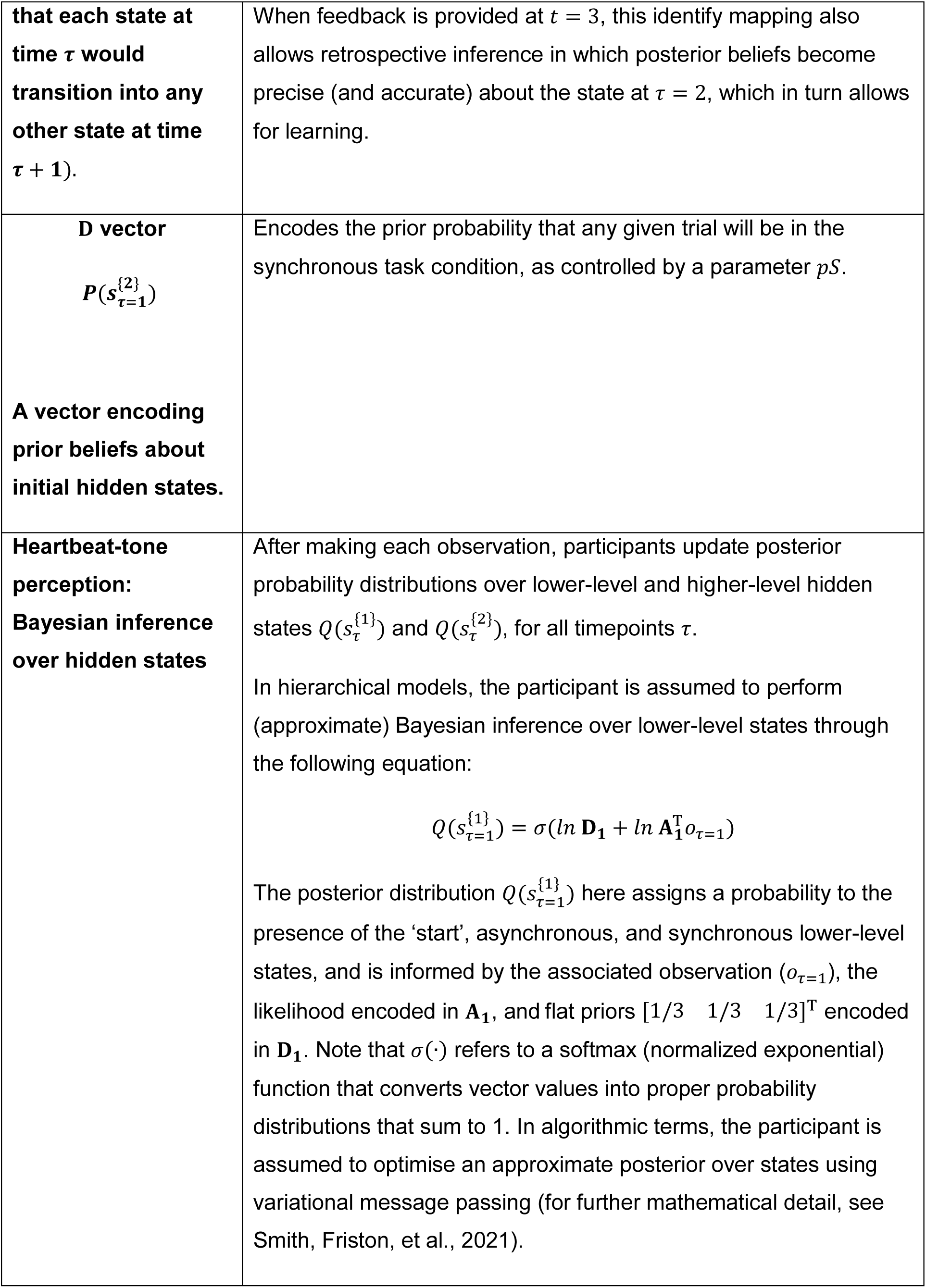
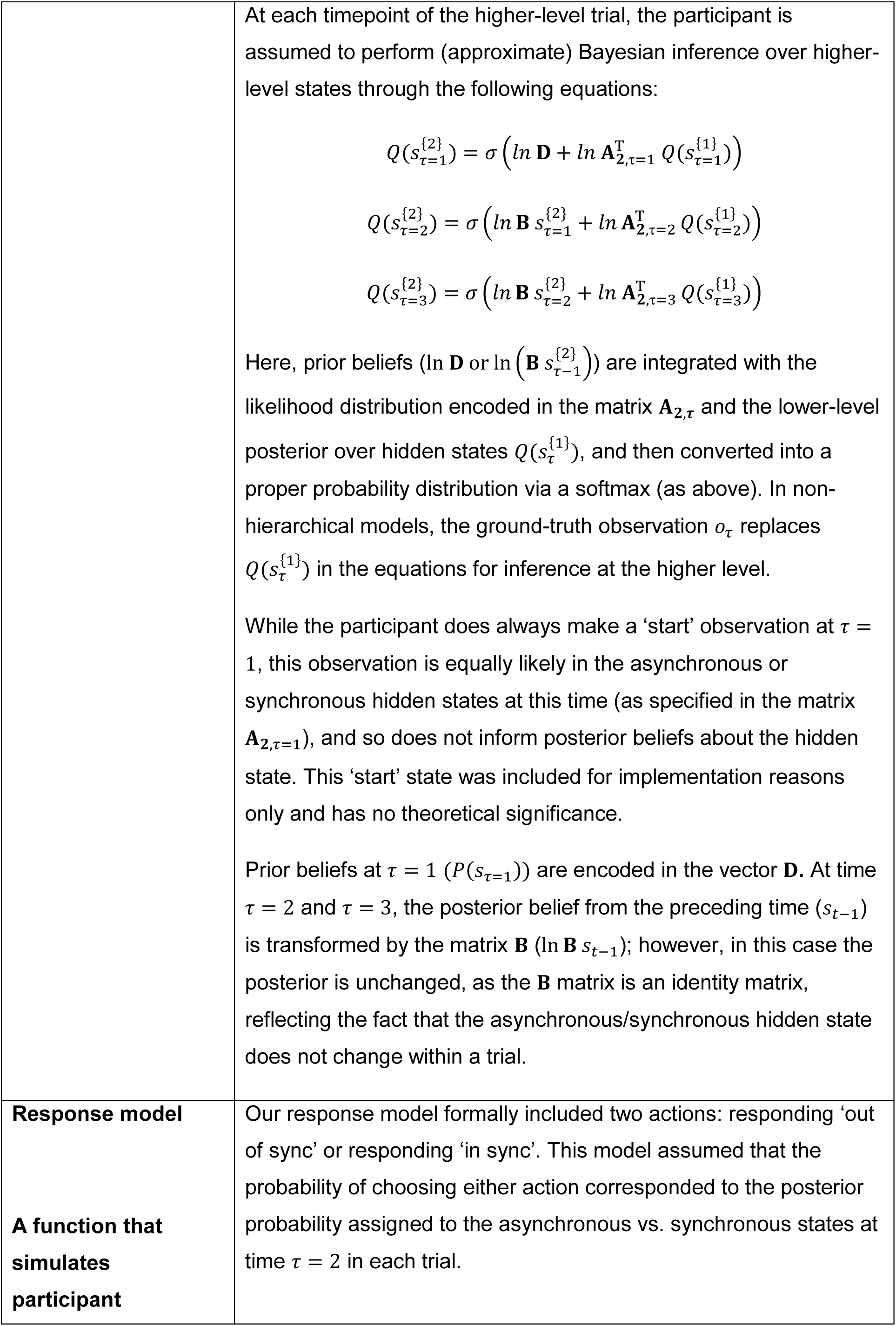
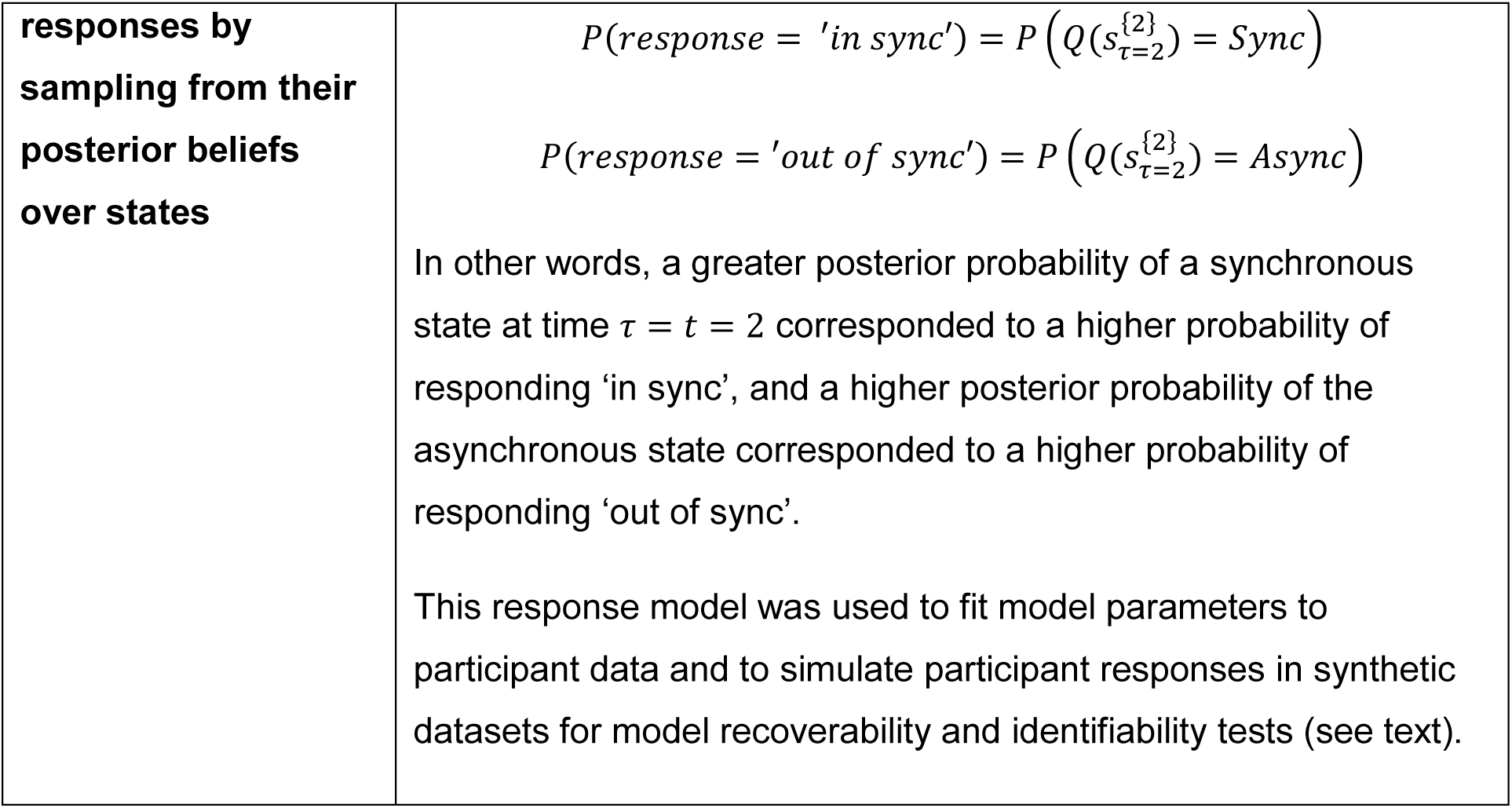
Description of computational model elements and processes.

Here we extended a hidden Markov model that has previously been used to implement Bayesian inference processes underlying interoceptive task behaviour in the cardiac and gastric domains (Lavalley et al., 2023; Smith, Kuplicki, Feinstein, Forthman, Stewart, Paulus, Investigators, et al., 2020; Smith, Kuplicki, Teed, et al., 2020; Smith, Mayeli, et al., 2021). Smith, Friston et al. (2021) and Da Costa et al. (2020) offer an overview of the structure and mathematics of the broader class of decision models (active inference models) from which the present model was adapted. **Table 1** gives full definitions of the various elements in the present model and explains the equations that governed (Bayes-optimal) perceptual inference and learning. MATLAB code used to implement this model and produce the computational modelling results is available on GitHub (https://github.com/ChatrinS/interoceptive-training-Bayesian-modelling).

#### Parameter estimation

For each model fitted to heartbeat discrimination task responses, we employed a Bayesian optimization algorithm to estimate the set of parameter values for each individual that best explained their task behaviour, as described by Schwartenbeck and Friston (2016). Parameter estimation was specifically carried out using variational Laplace (Friston et al., 2007), implemented using the spm_nlsi_Newton.m routine (freely available within the SPM12 software package; Wellcome Trust Centre for Neuroimaging, London, UK, http://www.fil.ion.ucl.ac.uk/spm). This estimation approach maximizes the log-likelihood of participant behaviour under a model while incorporating a complexity cost to deter overfitting (based on parameter covariance and divergence from prior values). The prior variance for estimation was set to .5 for each parameter, while prior means were set as follows: *IP*_1_ = .75, *IP*_2_ = .75, *pS* = .50, *η* = .50, *ω* = .75, *ω*_*Block*_ = .50, *ζ* = .50, *η*_D_ = .50, *IP*_1 *diff*_ = .25 (see **Table 2** for explanation of each parameter). Most prior means were set to the midpoint within the range of plausible values to minimize estimate bias. For example, prior means for *IP*_1_ and *IP*_2_ were set at the midpoint between completely imprecise (.50) and completely precise (1.00) mappings between outcomes and states, while the prior mean for *pS* assumes flat or balanced prior beliefs. An exception to this principle was inverse forgetting rate (*ω*), where initial simulations suggested that a midpoint prior mean (.50) produced implausibly fast forgetting over the course of trials, and so .75 was chosen to bias towards slower forgetting (i.e., higher values correspond to less forgetting). Another exception was *IP*_1 *diff*_, for which the prior mean was chosen to fit the bound *IP*_1_ + *IP*_1 *diff*_ ≤ 1.

**Table 2.**
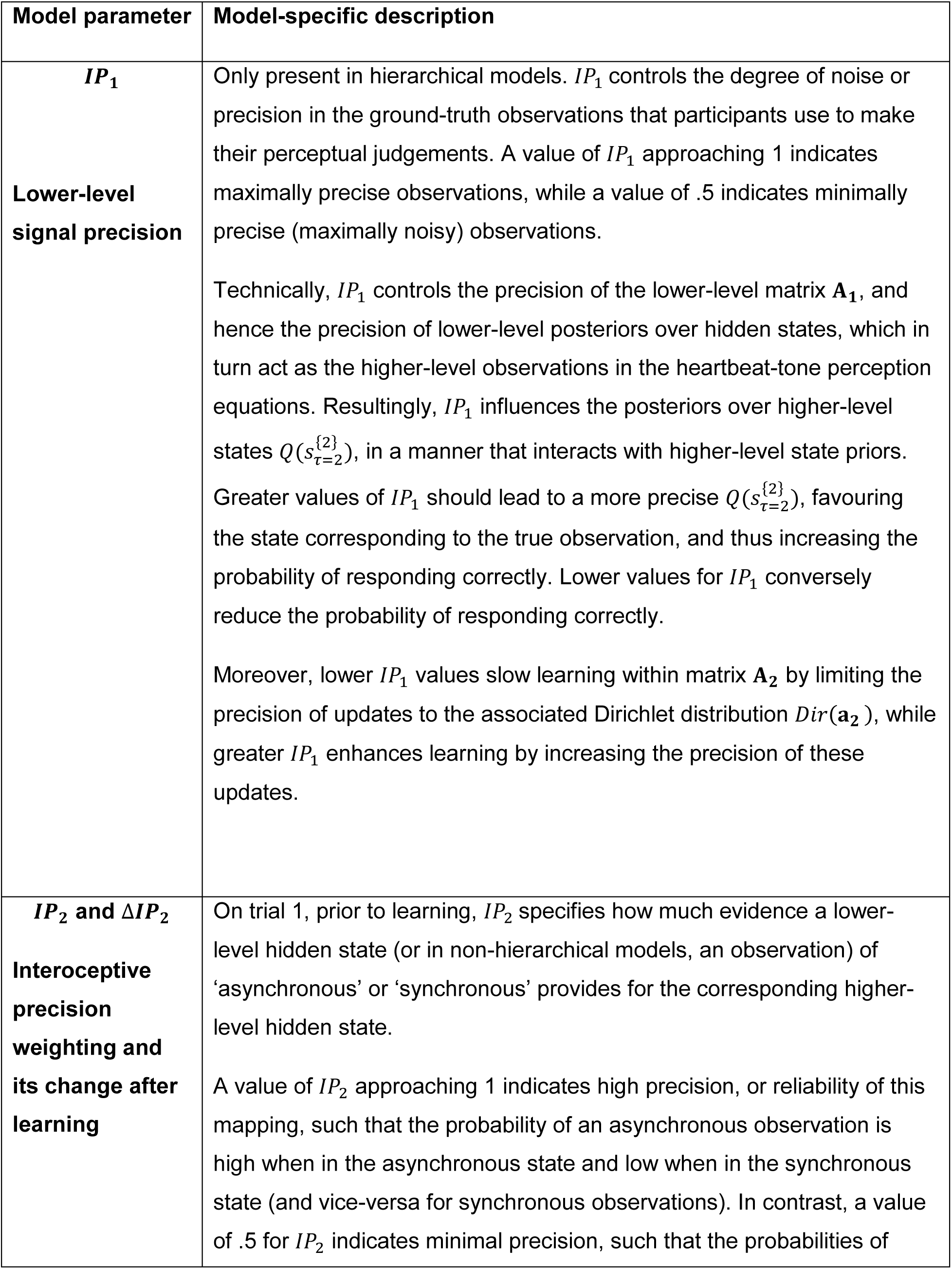
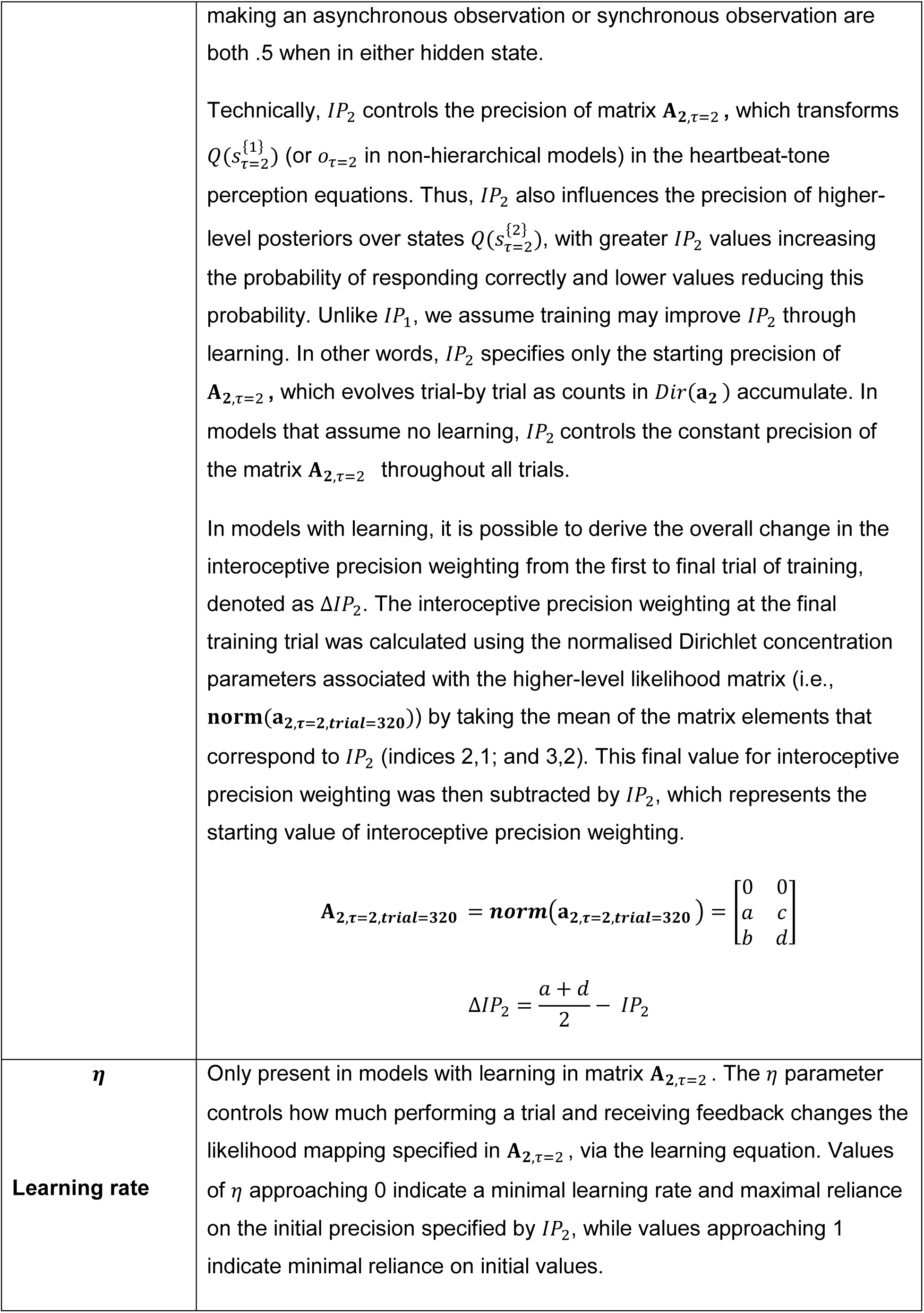
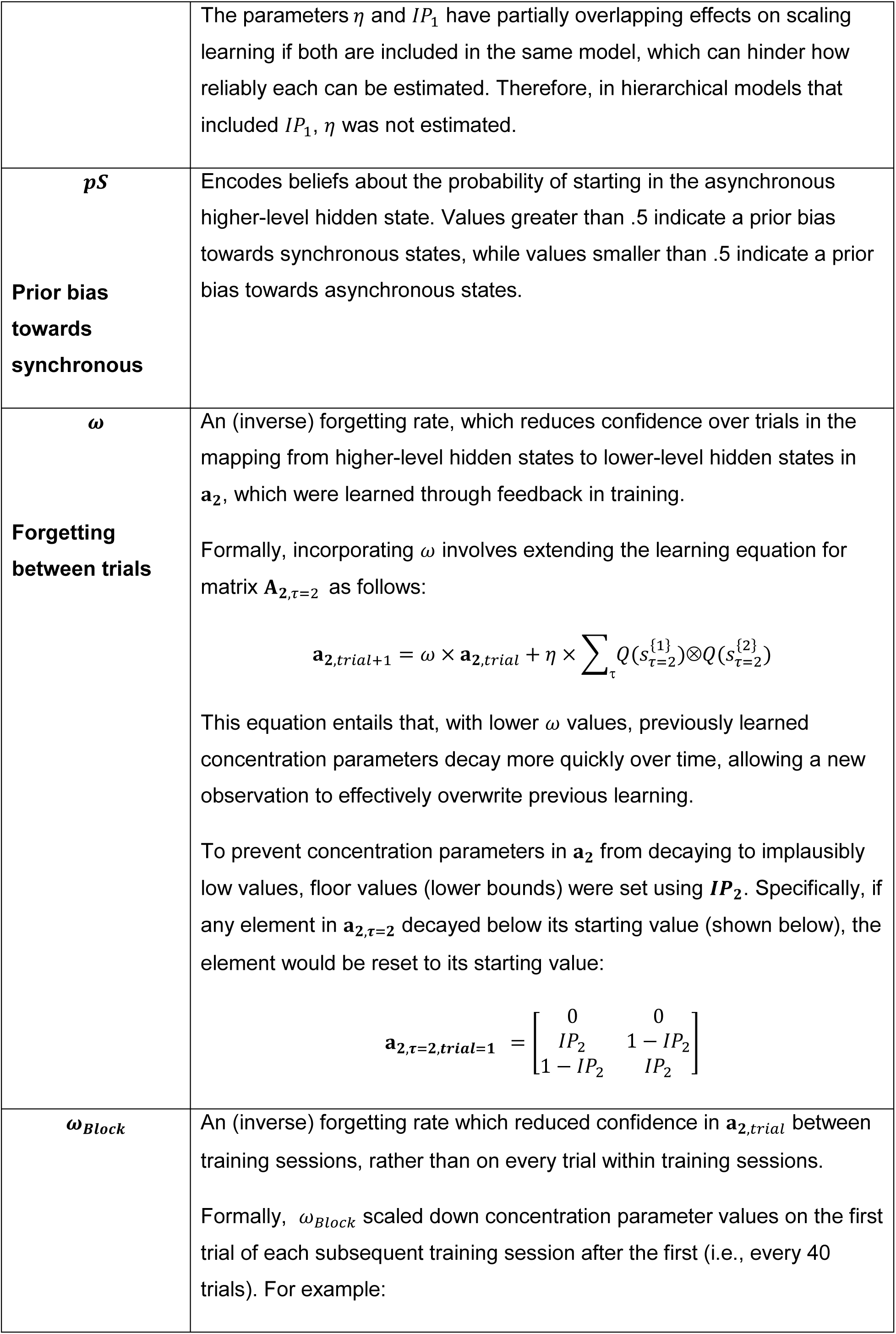
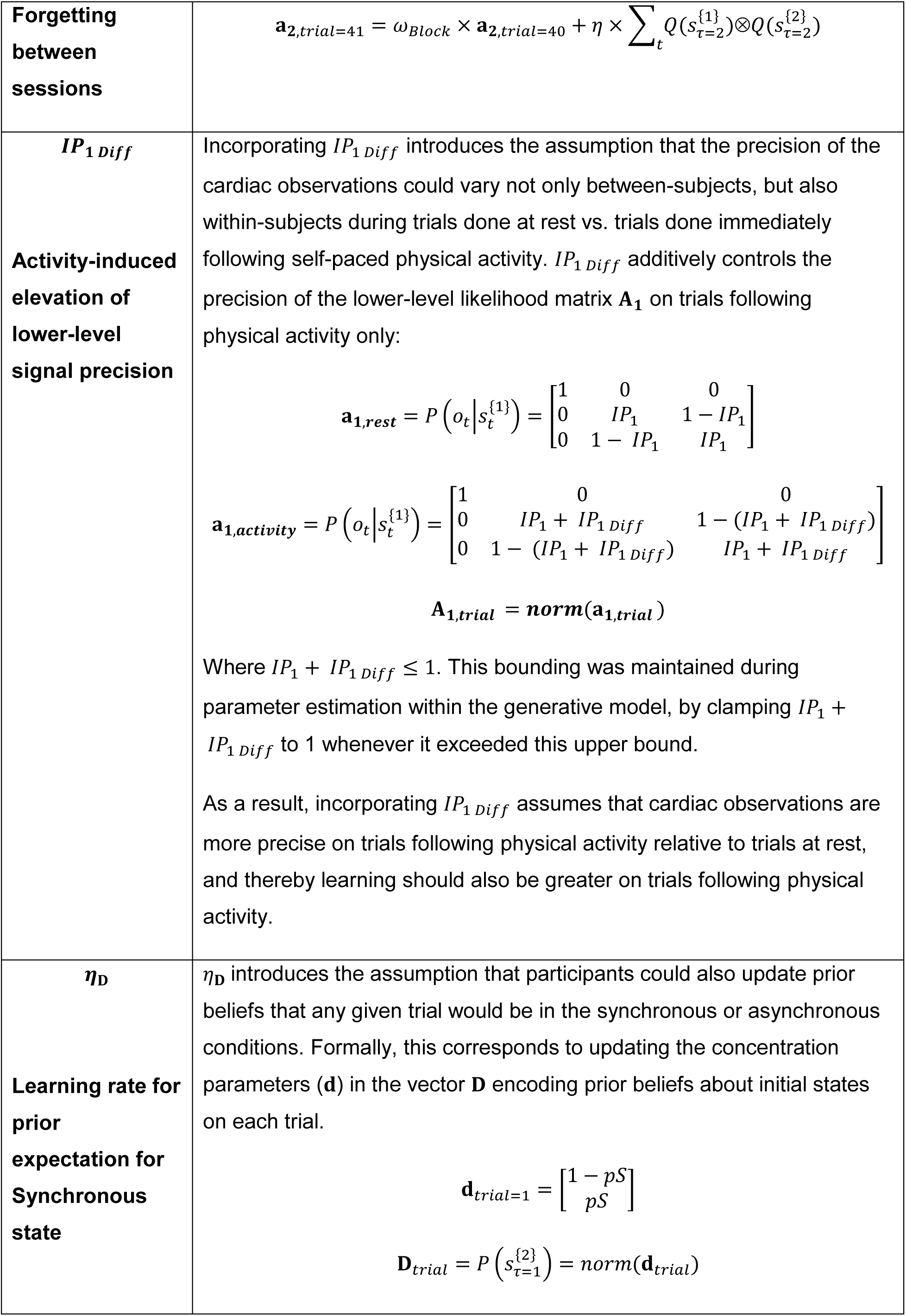
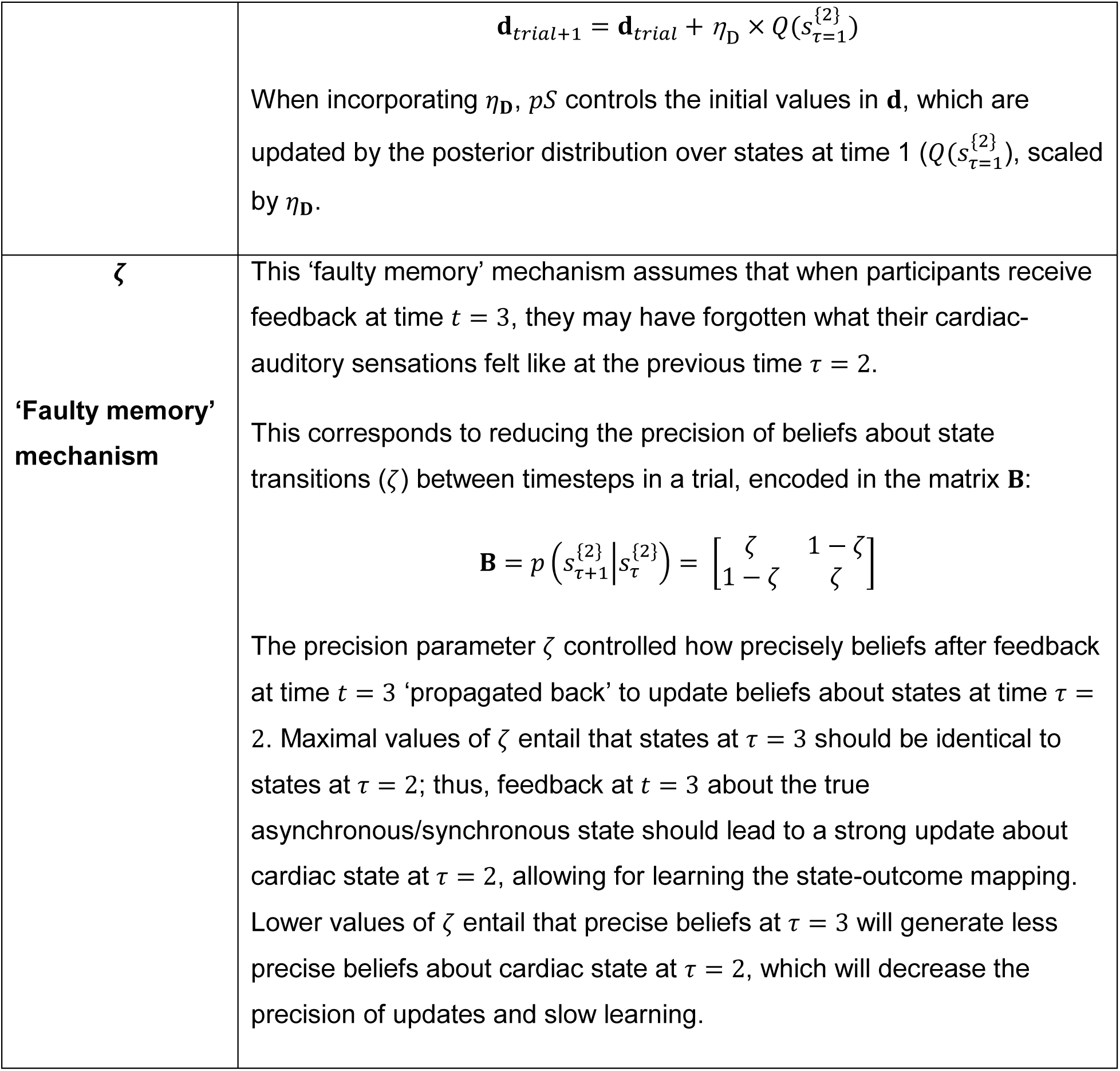
Description of computational model parameters estimated for each participant.

#### Model comparison, parameter recoverability, and model identifiability

**Table 2** explains the different computational parameters considered, which represent potential mechanisms of learning, forgetting, and Bayesian belief updating. **Table 3** displays which parameters were included in each model, each representing competing hypotheses about the mechanisms involved in interoceptive training. Model 1 was the simplest, which assumed that there was no learning, no hierarchical structure, and no prior bias, and that a static interoceptive precision weighting (*IP*_2_) could solely explain participants’ responses. Model 2 added the effect of prior bias (*pS*) on participant responses. Models 3 and 4 incorporated learning controlled by *η*, without or with prior bias, respectively. Models 5 through 13 incorporated the bottom-up effects of afferent cardiac signal precision (represented by *IP*_1_) and interactions between levels in a hierarchical structure. In these models, *IP*_1_ also effectively served as a (static) rate of evidence accumulation for learning *IP*_2_ over the course of trials, as higher values lead to more precise posteriors over first-level states, which in turn amplifies change in second-level precision estimates after each observation. Model 5 assumed no learning or prior bias within this hierarchical structure, while model 6 additionally included a prior bias. Models 7 and 8 assumed both learning and a hierarchical structure, without or with prior bias, respectively. Models 9 – 13 extended model 8 by considering additional learning and forgetting mechanisms: model 9 introduced an (inverse) forgetting between each trial (*ω*), while model 10 further posited forgetting between each training session (*ω*_*Block*_). Model 11 hypothesised heightened lower-level precision during physical activity blocks (*IP*_1 *Diff*_), while model 12 introduced learning for the prior bias (*η*_***D***_), and model 13 introduced a ‘faulty memory’ mechanism (*ζ*).

**Table 3.** Computational models compared.

| Parameter | $IP_1$ | $IP_2$ | $pS$ | $\eta$ | Additional parameters |
| --- | --- | --- | --- | --- | --- |
| **Model 1<sup>a</sup>** | N | Y | N | N | - |
| **Model 2<sup>a</sup>** | N | Y | Y | N | - |
| **Model 3<sup>a</sup>** | N | Y | N | Y | - |
| **Model 4<sup>a</sup>** | N | Y | Y | Y | - |
| **Model 5** | Y | Y | N | N | - |
| **Model 6** | Y | Y | Y | N | - |
| **Model 7** | Y | Y | N | N | - |
| **Model 8** | Y | Y | Y | N | - |
| **Model 9<sup>b</sup>** | Y | Y | Y | N | $\omega$ |
| **Model 10<sup>b</sup>** | Y | Y | Y | N | $\omega_{Block}$ |
| **Model 11** | Y | Y | Y | N | $IP_1 Diff$ |
| **Model 12** | Y | Y | Y | N | $\eta_D$ |
| **Model 13<sup>a</sup>** | Y | Y | Y | N | $\zeta$ |

**Legend.** Y indicates the parameter was estimated for that model, while N indicates the parameter was not. Models 1 – 4 were non-hierarchical, while models 5 – 13 were hierarchical. Models 1, 2, 5, and 6 assumed no learning occurs in matrix $A_{2,\tau=2}$ , while learning was present in all other models. When $IP_1$ or $\eta$ were not estimated, they were removed from the model (as mentioned in the main text, $\eta$ was not considered in hierarchical models due to overlapping effects with $IP_1$ ). When $pS$ was not estimated, it was instead fixed to .50 (i.e., no bias in prior beliefs).
<sup>a</sup> Excluded from final model comparison as model was not identifiable.
<sup>b</sup> Excluded from identifiability analysis and model comparison due to unrecoverable parameters.

Bayesian model comparison evaluated the relative evidence for each model provided by the behavioural data, to determine the best-fitting or ‘winning’ model – for details on this procedure, see Rigoux et al., (2014). This model comparison procedure produces the protected exceedance probability (*pxp*) and *α* metrics for each model compared. A model’s *pxp* quantifies the posterior probability that the model is more likely (given the data) than all other models being compared, while *α* quantifies the expected number of participants whose data was generated by the model.

Heartbeat discrimination task responses in the training group, concatenated across all training sessions (i.e., up to 320 trials with feedback), was used to determine the winning model (i.e., as only this group underwent training sessions for which learning would be expected and could be modelled).

Prior to performing Bayesian model comparison, the space of possible models listed above was first checked to confirm parameter recoverability and model identifiability. Only models that were recoverable and identifiable were then compared. Recoverability and identifiability were accomplished by first generating 13 synthetic datasets (one per model). This was done by simulating behavioural data for each participant under each model’s optimised parameter estimates. For parameter recovery, we then fit each of the 13 models on the synthetic dataset generated by itself, to produce parameter estimates, and then tested the correlation between estimated parameter values and the parameter values used to generate the simulated data. Models with parameters that proved unrecoverable (i.e., no significant correlation between the generative and estimated values) were eliminated from further consideration. Subsequently, we tested the identifiability of the remaining models by fitting them on all remaining synthetic datasets and passing the resulting model fits into Bayesian model comparison. A model was deemed identifiable if it was selected in Bayesian model comparison on the synthetic dataset that it generated, indicating that if this model was indeed the ground truth, it could be successfully ‘identified’ in model comparison. Non-identifiable models were excluded from consideration, and the remaining models were finally passed into Bayesian model comparison to determine the best-fitting model to the empirical data.

Synthetic datasets were also used to assess models in terms of their ability to reproduce the empirical data. To do so, we calculated for each participant the ‘model response accuracy’, or the proportion of trials in which simulated responses (in the model’s synthetic dataset) matched the participant’s actual responses during the task. The model response accuracy was averaged across training group participants and reported. We also calculated the ‘mean action probability’, or the probability of emitting the participant’s actual response (determined by the higher-level posterior over hidden states after stimulus presentation, 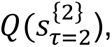 averaged across trials. The mean action probability was averaged across training group participants (as a mean of means) and reported.

Parameter estimates for the best-fitting model (i.e., out of those that were recoverable and identifiable) were then used for between-subjects analyses, as latent variables that best explained behavioural responses across training sessions for each participant. For each control group participant, who only had available data at assessment sessions, responses from up to 60 trials (without feedback) were concatenated and modelled together to produce complementary parameter estimates. Additionally, for both groups, responses from the three assessment sessions were modelled as three separate blocks, producing ‘snapshot’ parameter estimates for each participant at each timepoint.

### Interoceptive accuracy

Model-free measures of task performance were also calculated using responses during assessment sessions to index their change over time, following the ‘conventional’ approach for analysing task behaviour. Interoceptive accuracy on the heartbeat discrimination task was calculated using signal detection analysis (Stanislaw & Todorov, 1999): the number of hits (correct in-sync trials), misses (incorrect in-sync trials), correct rejections (correct out-of-sync trials) and false alarms (incorrect out-of-sync trials) were counted for each assessment session. The sensitivity index *d’* was calculated as *z*(*Hit rate*) − *z*(*False alarm rate*) and used as the measure of interoceptive accuracy for this task. Here, *z* denotes the left-tail *p*-value from the normal distribution. The Criterion (*C*), which specifies response bias, was also derived as 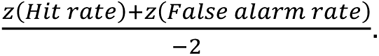

For the heartbeat counting task, accuracy in each trial was calculated using the number of heartbeats that occurred (*nbeats*_*real*_) and the number of heartbeats the participant reported in each trial (*nbeats*_*reported*_), as 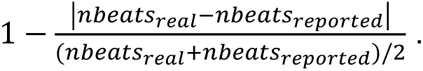 Resulting accuracy scores were averaged over the 6 trials, and used as the overall measure of interoceptive accuracy for this task (Garfinkel et al., 2015; Hart et al., 2013).

### Anxiety

State anxiety and trait anxiety were assessed using the Spielberger State-Trait Anxiety Inventory (Spielberger et al., 1983). This questionnaire is divided into two 20-question sections: the first section includes questions intended to capture current state anxiety, such as “*I feel strained*”, using a response scale which runs from “*Not at all*”, to “*Very much so*”. The second section targets a dispositional tendency for trait anxiety, and includes questions such as “*I worry too much over something that doesn’t really matter*”, with a response scale from “*Almost never*” to “*Almost always*”.

### Self-reported interoception

The Multidimensional Assessment of Interoceptive Awareness (MAIA) was used to measure subjectively perceived facets of interoception (Mehling et al., 2012). The MAIA is a 32-item self-report scale, divided into eight different subscales: 1) Noticing, assessing the awareness of uncomfortable, comfortable, or neutral body sensations, 2) Not-Distracting, assessing the tendency *not* to use distraction to cope with discomfort, 3) Not-Worrying, assessing the tendency *not* to experience emotional distress with physical discomfort, 4) Attention Regulation, assessing the reported ability to sustain and control attention to body sensations, 5) Emotional Awareness, assessing the reported ability to attribute specific physical sensations to physiological manifestations of emotions, 6) Self-Regulation scale, assessing the reported ability to regulate distress by attention to body sensations, 7) Body Listening scale, assessing the tendency to actively listen to the body for insight, and 8) Trusting scale, assessing the experience of one’s body as safe and trustworthy.

### Statistical Analyses

A series of linear mixed effects models (LMEs) tested the effects of group (training vs. control), time (baseline, midpoint, final), and the group by time interaction on conventional interoceptive task measures (tracking task accuracy, discrimination task *d’,* and *C*) and parameter estimates of the winning computational model derived from the assessment sessions (*IP*_1_, *IP*_2_, and *pS*). Similar LMEs tested the effects of group and time (baseline vs. final) and their interaction on self-reported anxiety (STAI-T and STAI-S) and self-reported interoception (scores on each MAIA sub-scale). Participant age and sex were controlled for as fixed effects. Sum-coding was used for group (control = -1, training = 1) and sex (male = -1, female = 1), while treatment coding was used for time (baseline = 0, midpoint = 1; and baseline = 0, final = 1). Age was mean-centered. Resultingly, coefficients for time are interpretable as main effects. All LMEs initially included the maximal random effects structure that was testable given the number of observations, but these were subsequently simplified to produce converging and non-singular model fits whenever required. In most cases, the above LMEs retained only random intercepts for each participant. Random effects structures and tabular model outputs are reported for all LMEs in **Supplementary Results.**

Another series of LMEs tested whether changes in trait and state anxiety from baseline to final assessment were associated with computational parameter estimates, in the training group only. Change scores for both trait and state anxiety were calculated for each participant, and used as the outcome variables for these LMEs. Estimates for the three computational parameters, *IP*_1_, *IP*_2_, and *pS* were included as fixed effects factors. Similar LMEs tested whether changes in trait and state anxiety could be explained by changes in conventional interoceptive task measures (tracking task accuracy, discrimination task *d’,* and *C*) from baseline to final assessment. Age and sex were again controlled for as fixed effects. Sex was sum-coded (male = -1, female = 1), and age was mean-centered, such that coefficients for other fixed effects factors could be interpreted as main effects. To determine whether baseline computational and interoceptive measures could predict future change in trait and state anxiety, these LMEs predicting change scores for trait and state anxiety were repeated using parameter estimates and model-free interoceptive task measures derived from the baseline assessment only. Again, these LMEs initially included the maximal random effects structure, which were subsequently simplified to produce converging and non-singular model fits. In most cases, LMEs retained a random intercept for Sex, and when no viable random effects structures could be found, ordinary multiple regression was used instead (see **Supplementary Results 6.1 – 6.3)**.

For all LMEs, the normality of residuals was assessed using Q-Q plots and Shapiro-Wilk tests (also reported in **Supplementary Results**). Whenever required, non-normally distributed variables were log-transformed using the R package ’optLog’ (https://github.com/kforthman/optLog), which identifies optimal log-transformations to minimize variable skew – log-transformations are noted in the results whenever they have been applied. Where multicollinearity was suspected in LMEs finding significant effects (i.e., if variance inflation factors [VIFs] > 5), predictors were removed from models until VIFs were all below 5, and ridge regression was performed to confirm that coefficient estimates remained in the same direction using the R library ‘ridge’ (Moritz et al., 2012). All relevant VIFs are reported in **Supplementary Results**. However, we also note that coefficient estimates and uncertainty estimates produced by LMEs have been shown to demonstrate robustness to violations of distributional assumptions (Schielzeth et al., 2020).

LMEs were implemented using the *lme4* and *lmerTest* packages within RStudio, and reported using the *sjPlot* package (Bates et al., 2015; Kuznetsova et al., 2017; Lüdecke, 2024; RStudio Team, 2022). Significance values and degrees of freedom for fixed effects were derived using Kenward-Roger approximations, following Luke (2017). Significant effects were interrogated with post-hoc pairwise comparisons using the *emmeans* package (Lenth et al., 2023), which calculated estimated marginal means (*EM*), standard errors (*SE*), and associated effect sizes (Cohen’s *d* using estimated marginal means). To account for the sex imbalance in the sample, estimated marginal means were calculated using proportional weights.

## Results

### Computational modelling

#### Model comparison, parameter recoverability, and model identifiability

Bayesian model comparison indicated that behavioural data provided the most evidence for model 8 (*IP*_1_, *IP*_2_, *pS*; protected exceedance probability [*pxp*] = .37; model response accuracy = .62, range .49 - .88; mean action probability = .62, range .51 - .87). This model comparison included six models that survived both parameter recovery and identifiability analyses **(Table 3)**. Results for parameter recovery and identifiability are shown in **Supplementary Results 1.1 – 1.2.**

We note that the evidence for model 8 increased (*pxp* = .83) when model 12 was additionally excluded from comparison. Model 12 incorporated an additional learning rate for prior biases favouring sync or async percepts, assuming this learning process differed between individuals). This is likely because although model 8 produced the best fit to the overall dataset (*α* = 15.52), model 12 produced a better fit to a small number of participants (*α* = 4.62). When model 12 was excluded, these participants are subsequently largely explained by model 8 (*α* = 18.88). Given this pattern, and the fact that model 8 is more parsimonious in containing fewer parameters, we move forward with subsequent analyses based on this model.

Parameter recovery for model 8 indicated good reliability for all three parameters (**Table 4**). Identifiability analysis for model 8 also supported its robustness: in a synthetic dataset generated by model 8, Bayesian model comparison again found the most evidence for model 8 (*pxp* = .90). This model posited that the interoceptive precision weighting is learned over time, with its starting value controlled by the parameter *IP*_2_.Here, it was possible to derive the overall change in the interoceptive precision weighting over the course of training (denoted by Δ*IP*_2_) for each participant using their optimised parameter values and the model equations that govern how interoceptive precision weighting evolves across trials (described in **Table 2**), allowing a model-based metric of individual training effectiveness.

**Table 4.** Tests of parameter recovery for model 8.

| Parameter | Range of generative values (i.e., based on participant estimates) | Correlation between generative values and estimated values |
| --- | --- | --- |
| $IP_1$ | .61 – .91 | $r(26) = .97, p < .001$ |
| $IP_2$ | .33 – .76 | $r(26) = .41, p = .03$ |
| $pS$ | .46 – .71 | $r(26) = .93, p < .001$ |

**Legend.** Correlation analyses tested the association between generative and estimated parameter values in a synthetic dataset simulated using model 8 and parameter values estimated for study participants.

#### Computational parameter estimates

Parameter values that best explained heartbeat discrimination task responses in the training group (across 320 training trials with feedback) and the control group (across 60 assessment trials without feedback) were estimated for *IP*_1_, *IP*_2_, and *pS*, and the derived change in interoceptive precision weighting (Δ*IP*_2_) was subsequently calculated. **Figure 3 (top)** illustrates descriptive correlations between estimated model parameters and indices of task responses, while **Figure 3 (bottom)** shows the distribution of parameter estimates in the training group and their correlations with each other, including Δ*IP*_2_. **Figure 4** illustrates the same measures for the parameter estimates in the control group, derived from concatenated assessment trials.

**Figure 3.**
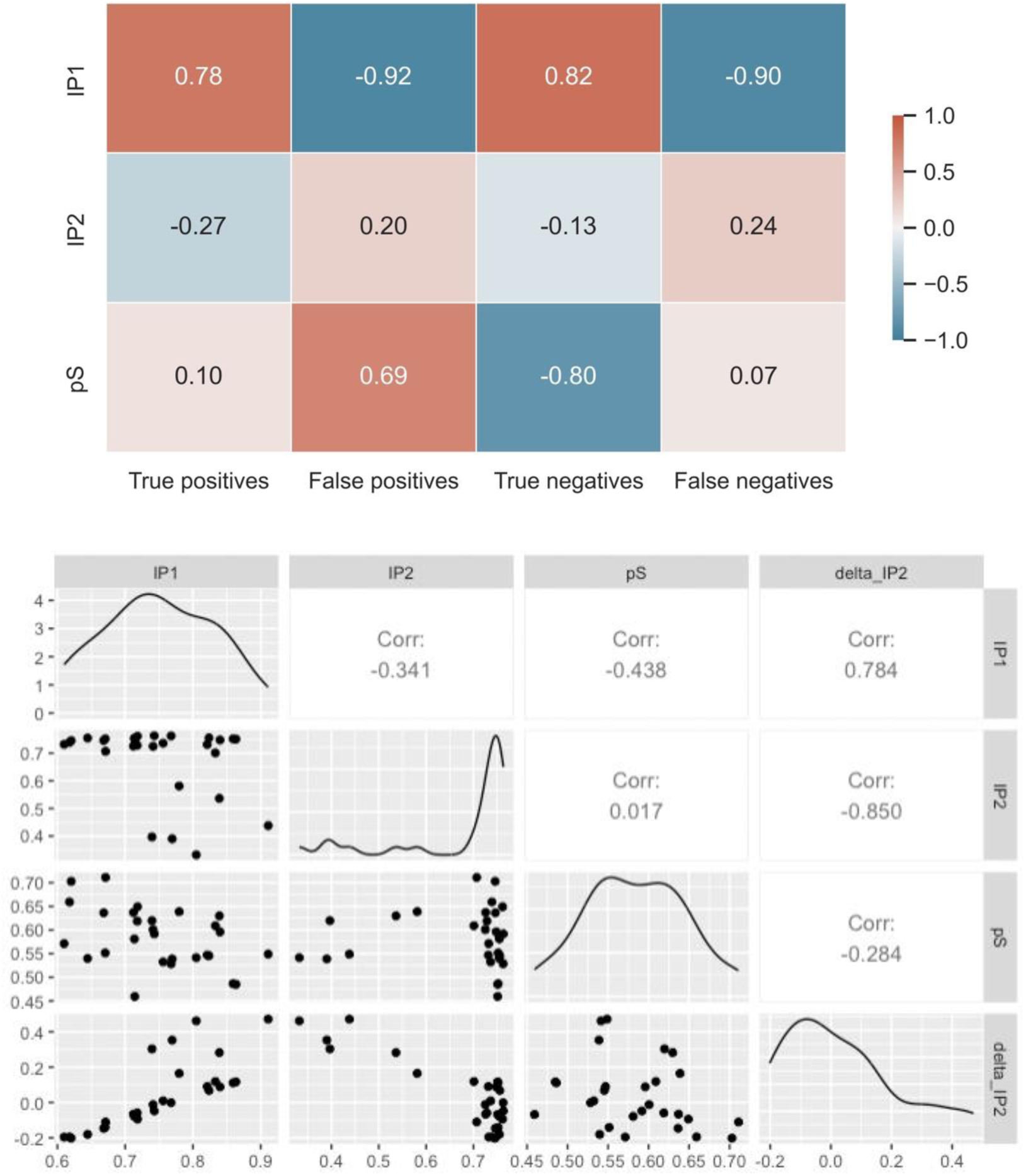
(Top) Pearson correlations between parameter estimates (*IP*_1_, *IP*_2_, *pS*) in the training group and confusion matrix indices derived from heartbeat discrimination task responses throughout the eight training sessions, showing largely expected relationships. Greater *IP*_1_ estimates were positively associated with true positive and true negative responses, and negatively associated with false positive and false negative responses. Parameter estimates for *IP*_2_ unsurprisingly showed weak associations with other measures, since this parameter determines only the starting conditions of the model prior to learning. Note that the few participants shown with starting values for *IP*_2_ in the .3 - .6 range would be expected to start with largely random (or even somewhat anti-correlated) responses relative to ground truth, but could still improve over time with learning. Greater *pS* estimates (i.e., more bias towards responding ‘in sync’) were associated with more false positives and fewer true negatives (as expected), but interestingly were only weakly associated with true positives and false negatives, suggesting that poorer performance on the task was partly driven by over-endorsement of ‘in sync’ responses. (Bottom) Pairwise Pearson correlations within the parameter estimates and the derived change in interoceptive precision weighting (Δ*IP*_2_), along with associated scatterplots and histograms for each variable. Significance indicators are not shown, as these were descriptive analyses to inform parameter face validity, rather than hypothesis tests.

**Figure 4.**
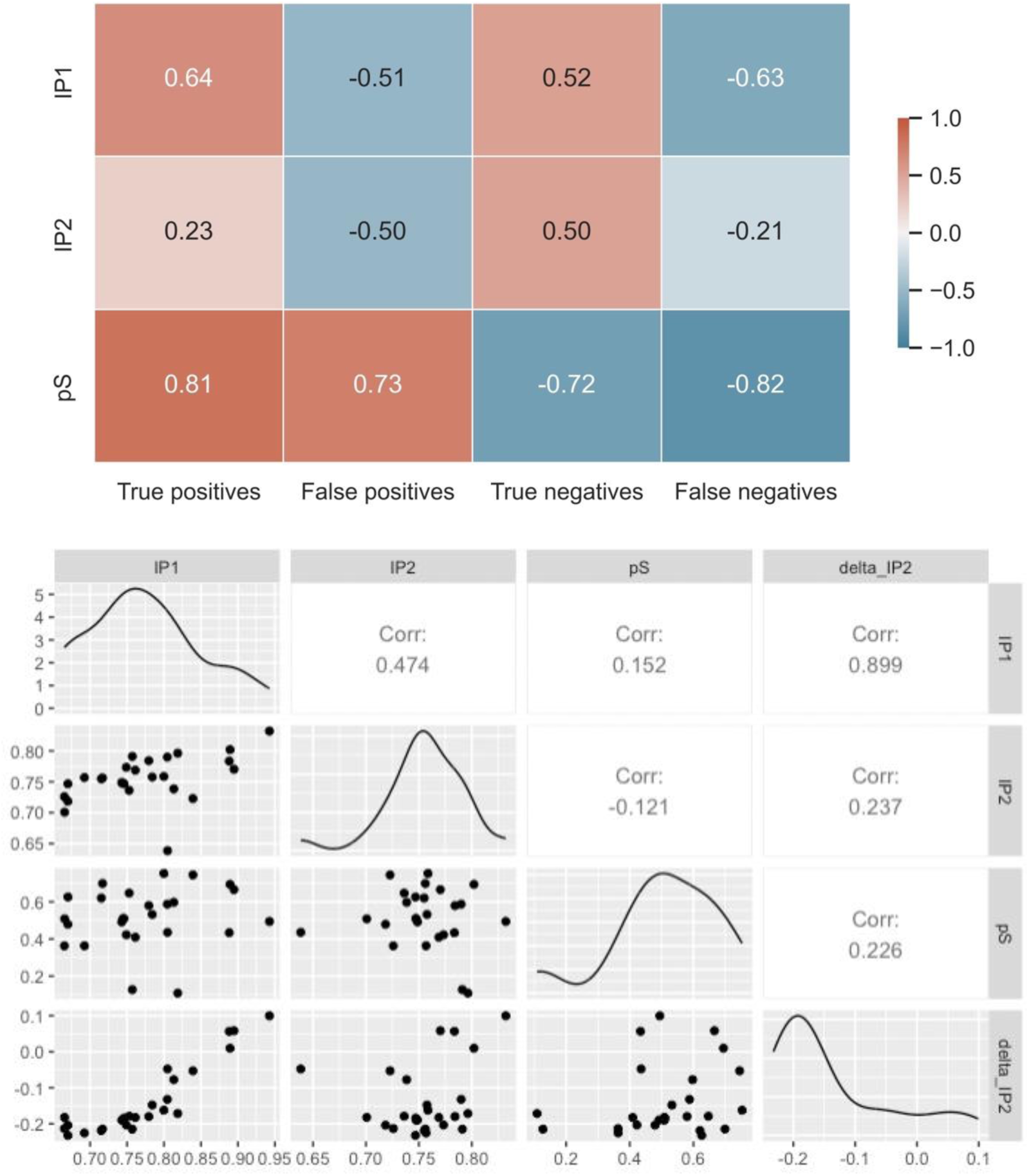
(Top) Pearson correlations (Corr) between parameter estimates in the control group and confusion matrix indices derived from heartbeat discrimination task responses across three assessment sessions. Note that both *IP*_2_ and *pS* showed qualitatively different patterns of association with behaviour in the control group compared to the training group, likely reflecting the difference in experimental procedure for the data used to estimate parameter values (i.e., no feedback on each trial and fewer trials overall in the control group). (Bottom) Pearson correlations within parameter estimates and the derived change in interoceptive precision weighting in the control group, along with associated scatterplots and histograms for each parameter. Significances of these descriptive patterns were not tested, as these did not reflect hypothesis tests.

Parameter estimates in the training group were normally distributed for *IP*_1_ (*W* = .97, *p* = .60) and *pS* (*W* = .98, *p* = .75), but were non-normally distributed for *IP*_2_ (*W* = .64, *p* < .001) and Δ*IP*_2_ (*W* = .91, *p* = .02). In the control group, *IP*_1_ (*W* = .95, *p* = .28), *IP*_2_ (*W* = .94, *p* = .13), and *pS* (*W* = .93, *p* = .07) were normally distributed, while Δ*IP*_2_ was not (*W* = .80, *p* < .001). When pooling both groups together, *IP*_1_ was normally distributed (*W* = .98, *p* = .69), but *IP*_2_(*W* = .63, *p* < .001), Δ*IP*_2_ (*W* = .85, *p* < .001), and *pS* (*W* = .89, *p* < .001) were non-normally distributed. As such, computational variables that were non-normally distributed were log-transformed for further statistical analysis (noted whenever applicable).

#### Relationship with heartrate and conventional interoceptive task measures

Across both groups, the mean heartrate taken across all trials of the heartrate discrimination task was significantly associated with *IP*_1_ parameter estimates (*r*(52) = -.33, *p* = .01), such that a lower heartrate across all trials was associated with higher *IP*_1_. Multiple regression analyses indicated that this effect was significant independent of contributions from other parameter estimates, age, and sex (*β* = -0.003, *SE* = 0.001, *t*(20.00) = -2.87, *p* = .01; **Figure 5**, **Supplementary Results 2.1**). The effect remained significant when considering the training group only (**Supplementary Results 2.2**). This could be seen to support the assumption made in the computational model that *IP*_1_ represents a cardiovascular trait corresponding to the physiological afferent signal precision, which would be expected to moderate learning (i.e., an objectively noisier signal would be more difficult to learn from).

**Figure 5.**
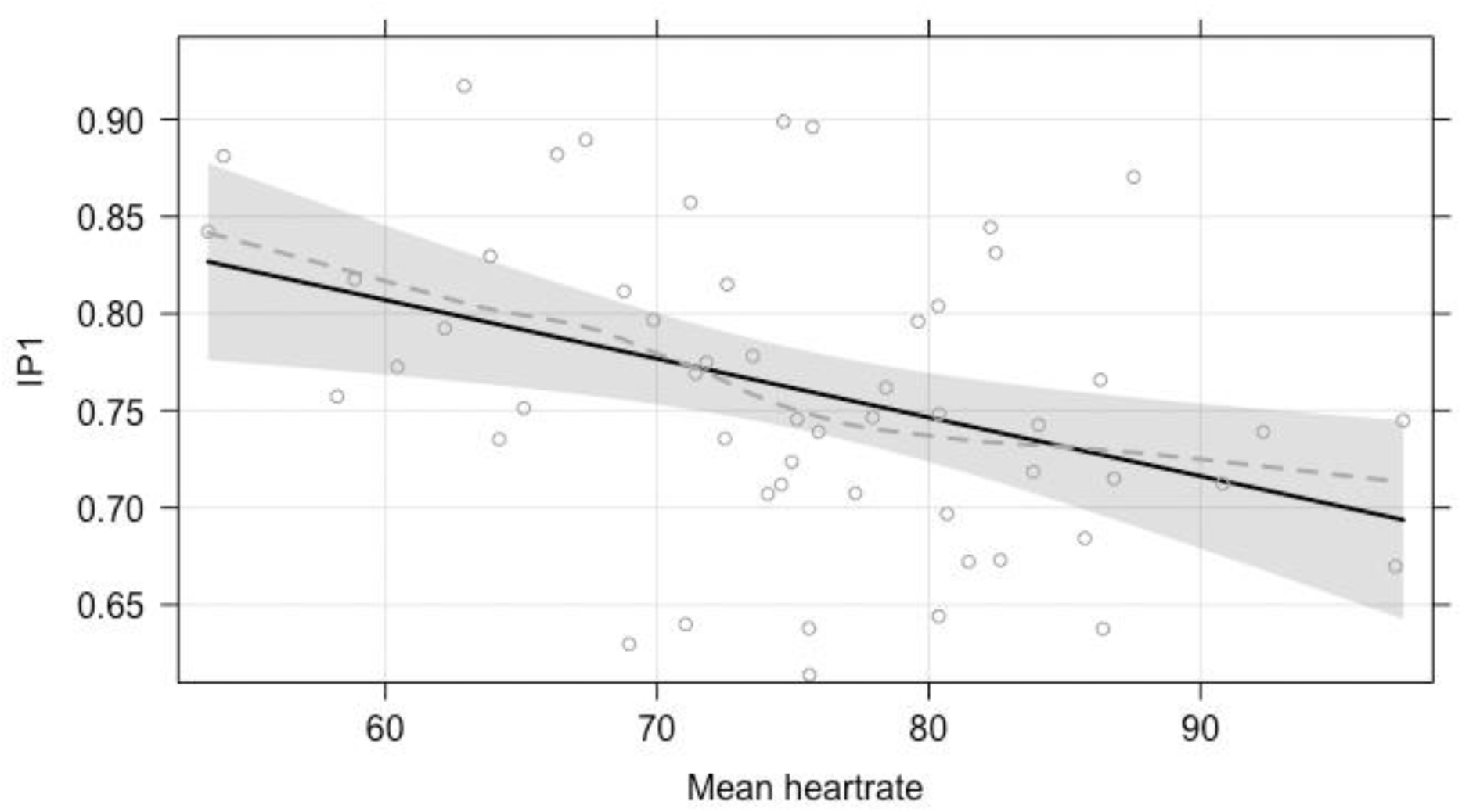
Effect plot with partial residuals for significant association between mean heartrate across heartbeat discrimination trials and estimates of the afferent signal precision (*IP*_1_) across both groups. The solid line visualises the partial slope for the predictor (i.e., when all other fixed effects are held constant), as estimated using multiple regression. Grey circles represent partial residuals (i.e., the dependent variable adjusted for all other fixed effects), while the dashed grey line visualises the loess smooth of partial residuals. Effect plots were generated using the *Effects* R package (Fox & Weisberg, 2018).

Conventional measures of perceptual accuracy in the heartbeat discrimination and heartbeat counting tasks improved over time in the training group, but not the control group (**Figure 6**). The training group showed a significant increase in tracking task accuracy from the baseline (estimated marginal mean [*EM*] = 0.53, standard error [*SE*] = 0.04) to midpoint assessments (*EM* = 0.78, *SE* = 0.04; *t*(102) = 6.99, *p* < .001, Cohen’s *d* = 1.87), with no subsequent improvement between the mid-point and final assessments (*EM* = 0.83, *SE* = 0.04; *t*(102) = 1.30, *p =* .20, Cohen’s *d* = 0.35). The control group showed no significant improvement from the baseline (*EM* = 0.59, *SE* = 0.04) to midpoint assessments (*EM* = 0.59, *SE* = 0.04; *t*(103) = -0.07, *p* = .94, Cohen’s *d* = 0.02) or final assessments (*EM* = 0.65, *SE* = 0.04; *t*(103) = 1.66, *p* = .10, Cohen’s *d* = 0.47; linear mixed model reported in **Supplementary Results 3.1)**.

**Figure 6.**
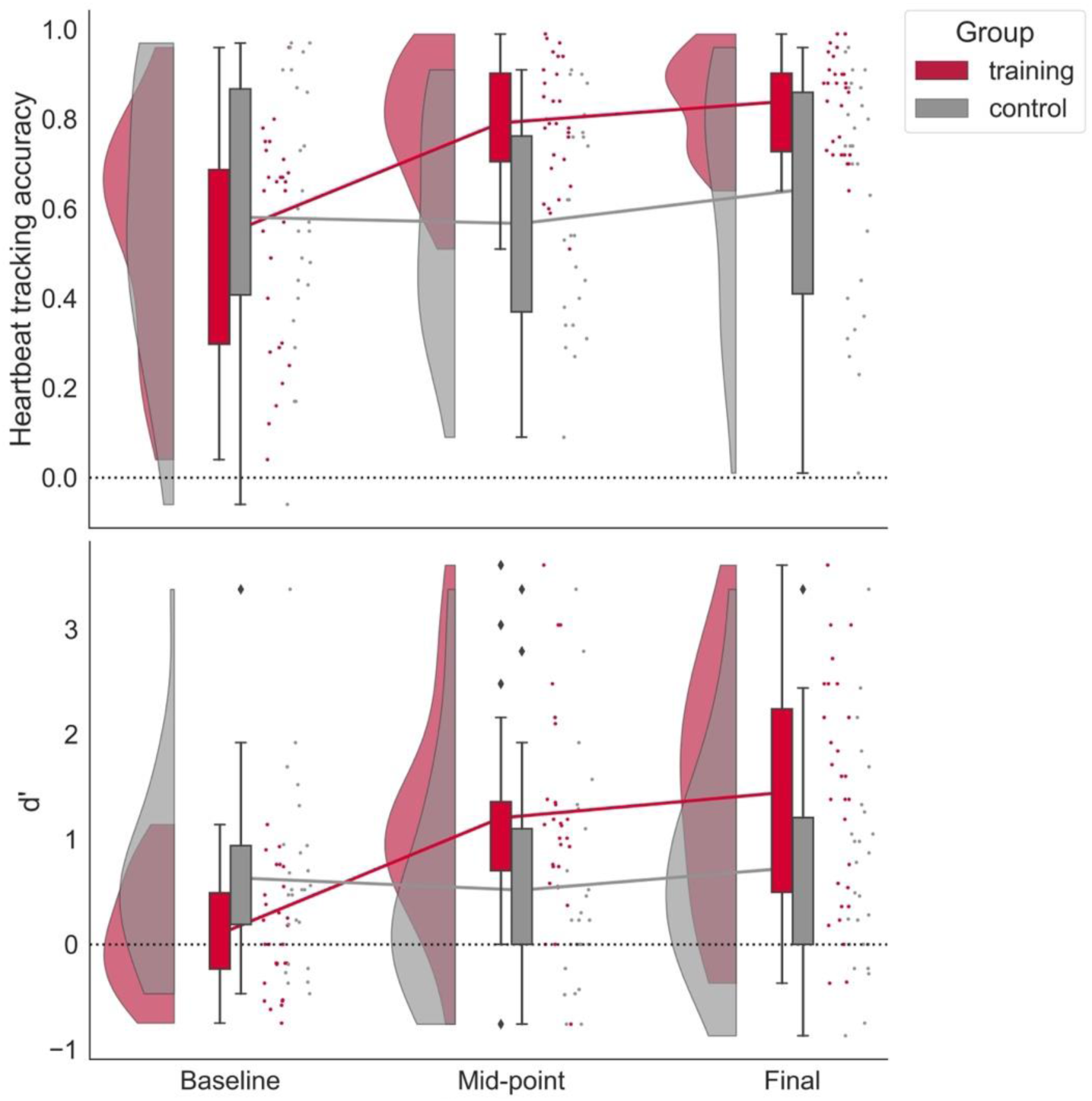
Raincloud plots showing heartbeat tracking accuracy (top) and heartbeat discrimination sensitivity (*d’*; bottom) scores for the training and control groups across baseline, mid-point and final assessment sessions. Lines across plots connect the group means at each timepoint. Outliers are indicated by diamonds above and below box and whisker plots. All raincloud plots were produced using the PtitPrince Python package (Allen et al., 2021).

The training group showed a significant increase in discrimination task *d’* between the baseline (*EM* = 0.12, *SE* = 0.18) and mid-point assessments (*EM* = 1.22, *SE* = 0.18; *t*(101) = 6.38, *p* < .001, Cohen’s *d* = 1.70), with no subsequent improvement between the mid-point and final assessments (*EM* = 1.46, *SE* = 0.18; *t*(101) = 1.39, *p =* .17, Cohen’s *d* = .37). The control group again showed no significant improvement from the baseline (*EM* = 0.58, *SE* = 0.19) to midpoint assessments (*EM* = 0.54, *SE* = 0.19; *t*(103) = -0.25, *p* = .81, Cohen’s *d* = 0.07) or final assessments (*EM* = 0.71, *SE* = 0.19; *t*(102) = 0.97, *p* = .33, Cohen’s *d* = 0.27; linear mixed model reported in **Supplementary Results 3.2)**.

Importantly, gains in heartbeat discrimination accuracy – indexed by change in signal detection *d’* from the baseline to final timepoints – were positively correlated with the increase in interoceptive precision weighting (Δ*IP*_2_), both within the training group (Spearman’s *r_s_*(26) = .67, *p* < .001) and across both groups combined (*r_s_*(52) = .65, *p* < .001). *IP*_1_ estimates were significantly positively correlated with increase in signal detection *d’* from baseline to final timepoints (Pearson’s *r*(26) = .64, *p* < .001) and with Δ*IP*_2_ in the training group (*r_s_*(26) = .87, *p* < .001). Similar associations were found when combining both groups for signal detection *d’* (*r*(51) = .31, *p* = .023) and Δ*IP*_2_: (*r_s_*(52) = .68, *p* < .001).

Furthermore, a mediation analysis including both groups indicated that the relationship between *IP*_1_ (as the independent variable) and the change in signal detection *d’* (as the dependent variable) was fully mediated by log-transformed Δ*IP*_2_ (Average Causal Mediation Effect = 6.12, *p* < .001; Average Direct Effect = -1.55, *p* = .41; Total Effect = 4.63, *p* = .007; proportion mediated = 1.34, *p* = .007). These results were consistent with the hypotheses that learning to detect cardiac signals was underpinned by increasing the precision weighting afforded to cardiac signals and that the (lower-level) precision of the afferent signal itself moderated the rate of learning.

#### Computational parameter estimates from separate assessment sessions

Computational parameters were also estimated on heartbeat discrimination task responses in each assessment session separately, producing ‘snapshot’ computational measures at each timepoint for both groups.

*IP*_1_ significantly increased in the training group from the baseline (*EM* = 0.72, *SE* = 0.01) to mid-point (*EM* = 0.81, *SE* = 0.01) assessments (*t*(103) = 6.10, *p* < .001, Cohen’s *d* = 1.63), while the subsequent increase from mid-point to final assessment (*EM* = 0.83, *SE* = 0.01) was non-significant (t(103) = 1.67, *p* =.10, Cohen’s *d* = 0.45; **Figure 7, top;** linear mixed model reported in **Supplementary Results 4.1**). In contrast, the control group showed no significant increases in *IP*_1_ from the baseline (*EM* = 0.74, *SE* = 0.01) to midpoint assessments (*EM* = 0.75, *SE* = 0.01; *t*(104) = 0.61, *p* = .54, Cohen’s *d* = 0.17) or final assessments (*EM* = 0.77, *SE* = 0.01 *t*(104) = 1.17, *p* = .24, Cohen’s *d* = 0.33).

**Figure 7.**
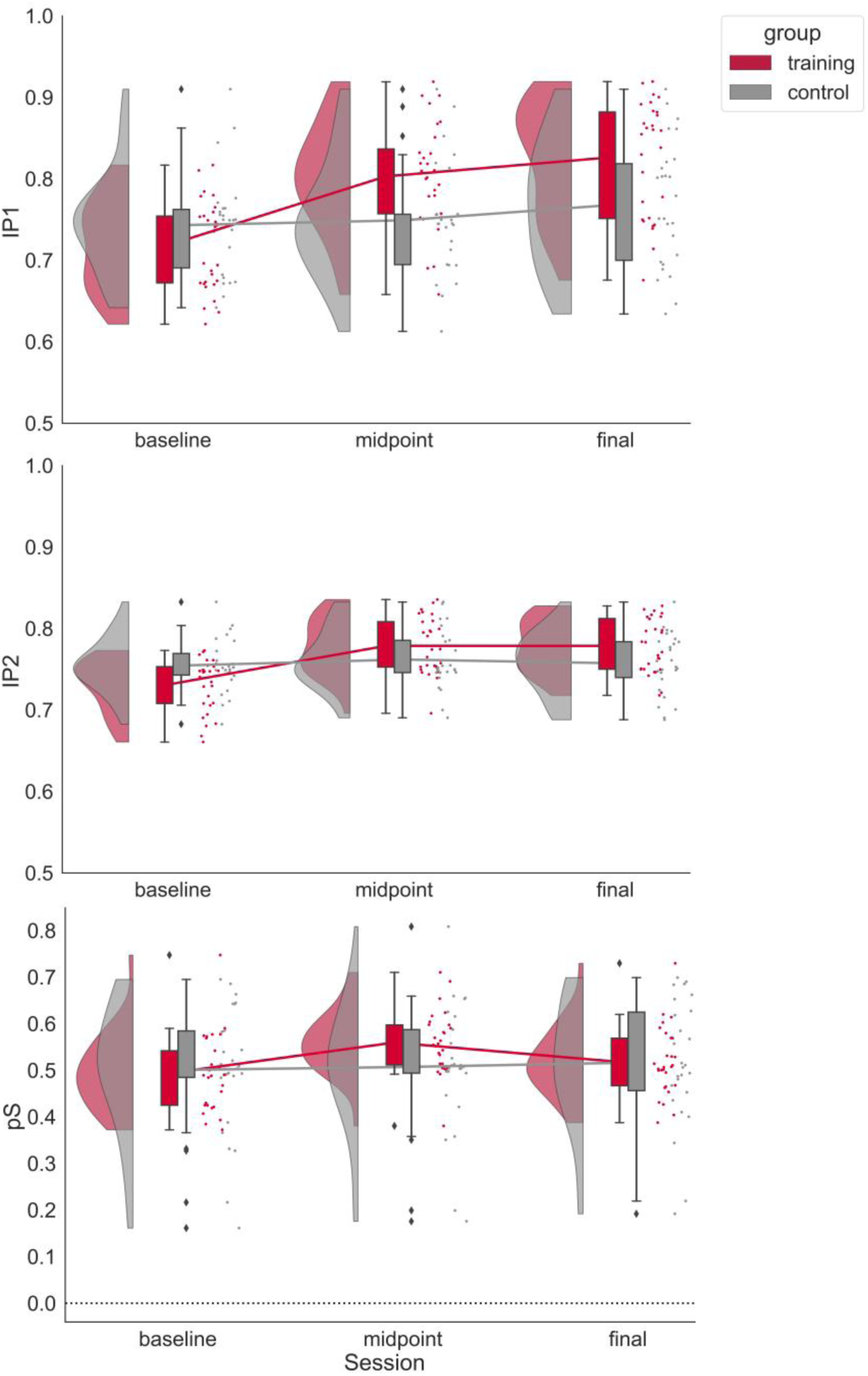
Raincloud plots showing *IP*_1_ (top), *IP*_2_(middle) and *pS* (bottom) estimates for the training and control groups across baseline, mid-point, and final sessions.

The training group also showed significant increases in *IP*_2_ from the baseline (*EM* = 0.73, *SE* = 0.006) to mid-point (*EM* = 0.78, *SE* = 0.006) assessments (*t*(103) = 6.32, *p* < .0001, Cohen’s *d* = 1.69), but not between the mid-point to final assessments (*EM* = 0.78, *SE* = 0.006; *t*(103) = -0.01, *p* = .99, Cohen’s *d* = 0.00; **Figure 7, middle**; **Supplementary Results 4.2**). The control group again did not show any significant differences in *IP*_2_ estimates from the baseline (*EM* = 0.75, *SE* = 0.01) to midpoint assessments (*EM* = 0.76, *SE* = 0.01; *t*(104) = 1.02, *p* = .31, Cohen’s *d* = 0.29) or final assessments (*EM* = 0.76, *SE* = 0.01 *t*(104) = - 0.69, *p* = .49, Cohen’s *d* = 0.19).

The training group showed a significant increase in *pS* from the baseline (*EM* = 0.50, *SE* = 0.02) to midpoint assessments (*EM* = 0.56, *SE* = 0.02; *t*(103) = 3.10, *p* = .003, Cohen’s *d* = 0.83), but a subsequent decrease between the mid-point and final assessments (*EM* = 0.52, *SE* = 0.02; *t*(103) = -2.03, *p =* .045, Cohen’s *d* = 0.54; **Figure 7, bottom**; **Supplementary Results 4.3**). Recall that a value of *pS* > .5 indicates biased prior beliefs towards synchronous cardiac-auditory stimuli (and biased towards asynchronous stimuli below .5); thus, based on the Ems, these results indicate the temporary induction of a positive bias that subsequently returned to unbiased values by the end of training. The control group showed no significant change in *pS* from the baseline (*EM* = 0.50, *SE* = 0.02) to midpoint assessments (*EM* = 0.51, *SE* = 0.02; *t*(104) = 0.45, *p* = .66, Cohen’s *d* = 0.13) or final assessments (*EM* = 0.51, *SE* = 0.2 *t*(104) = 0.29, *p* = .77, Cohen’s *d* = 0.08)

### Anxiety

#### Baseline anxiety and change over time

Participants showed a moderate range of subclinical anxiety scores on the STAI at baseline (max: 80 points, range 23-70 for trait anxiety; 21-71 for state anxiety). Note that baseline trait anxiety was normally distributed (*W* = .98, *p* = .41), while baseline state anxiety was not (*W* = .92, *p* < .01).

State anxiety showed a non-significant decrease in the training group [baseline: *EM* = 40.5, *SE* = 2.06; final: *EM* = 37.1, *SE* = 2.06; *t*(52) = -1.55, *p* = .13, Cohen’s *d* = 0.41], and a non-significant increase in the control group (baseline: *EM* = 36.6, *SE* = 2.14; final: *EM* = 40.0, *SE* = 2.14; *t*(52)= 1.50, *p* = .14, Cohen’s *d* = 0.42) [**Figure 8, top**]. These changes resulted in a significant group by time interaction (*β* = -3.37, *SE* = 1.56, *t*(52) = -2.15, *p* = .036). The two groups did not differ significantly in baseline state anxiety (*β* = 3.83, *SE* = 3.00, *t*(83.2) = 1.28, *p* = .21). Age was significantly associated with greater state anxiety (*β* = 0.36, *SE* = 0.16, *t*(50) = 2.22, *p* = .03), while participant sex was not (**Supplementary Results 5.1**).

**Figure 8.**
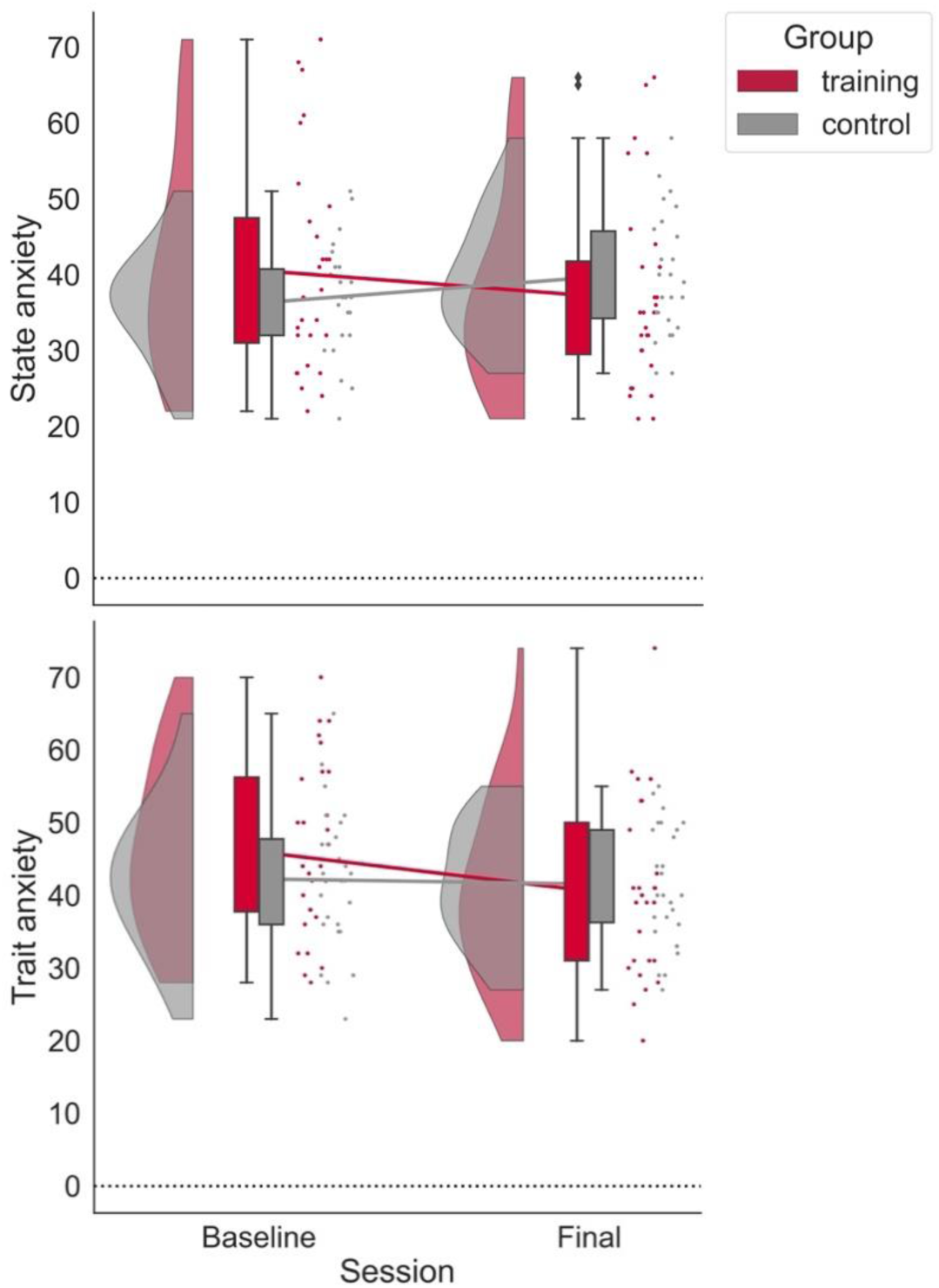
(Top) State anxiety showed a non-significant decrease in the training group and a non-significant increase in the control group, resulting in a significant interaction effect. (Bottom) Trait anxiety significantly decreased in the training group, but not in the control group. Note that these anxiety assessments were not gathered at the mid-point visit.

Trait anxiety decreased in the training group from baseline (*EM* = 46.2, *SE* = 2.01) to final assessment (*EM* = 40.8, *SE* = 2.01; *t*(52) = -4.12, *p* < .001, Cohen’s *d* = 1.10). There was no significant change in trait anxiety within the control group (baseline: *EM* = 42.2, *SE* = 2.09; final: *EM* = 41.5, *SE* = 2.09; *t*(52) = -0.51, *p* = .61, Cohen’s *d* = 0.14; **Figure 8, bottom**), resulting in a significant group by time interaction (*β* = -2.35, *SE* = 0.94, *t*(52) = -2.49, *p* = .02). The two groups did not differ significantly in baseline trait anxiety (*β* = 3.83, *SE* = 2.92, *t*(61.1) = 1.29, *p* = .20). Neither participant age nor sex were significantly associated with trait anxiety (**Supplementary Results 5.2**).

#### Relationships between change in anxiety and computational parameters

In the training group, we tested whether reduction in state and/or trait anxiety following interoceptive training could be explained by computational parameter estimates using LMEs. LMEs initially included each model parameter (*IP*_1_, log-transformed *IP*_2_, and *pS*) and the derived change in interoceptive precision weighting (log-transformed Δ*IP*_2_), while controlling for baseline state anxiety, age, and sex as fixed effects (**Supplementary Results 6.1.1 – 6.1.2**). However, multicollinearity was suspected in these LMEs, and so the log-transformed *IP*_2_ was later excluded from the predictors, such that VIFs we all below 5.

This analysis revealed that weaker prior biases toward perceiving synchrony (*pS*) were associated with greater decreases (or smaller increases) in state anxiety (*β* = 123.18, *SE* = 42.09, *t*(21.00) = 2.93, *p* = .008; **Figure 9, top; Supplementary Results 6.1.3**), while greater baseline state anxiety was associated with greater decreases in state anxiety (as expected due to regression to the mean; *β* = -0.70, *SE* = 0.15, *t*(21.00) = -4.59, *p* < .001).

**Figure 9.**
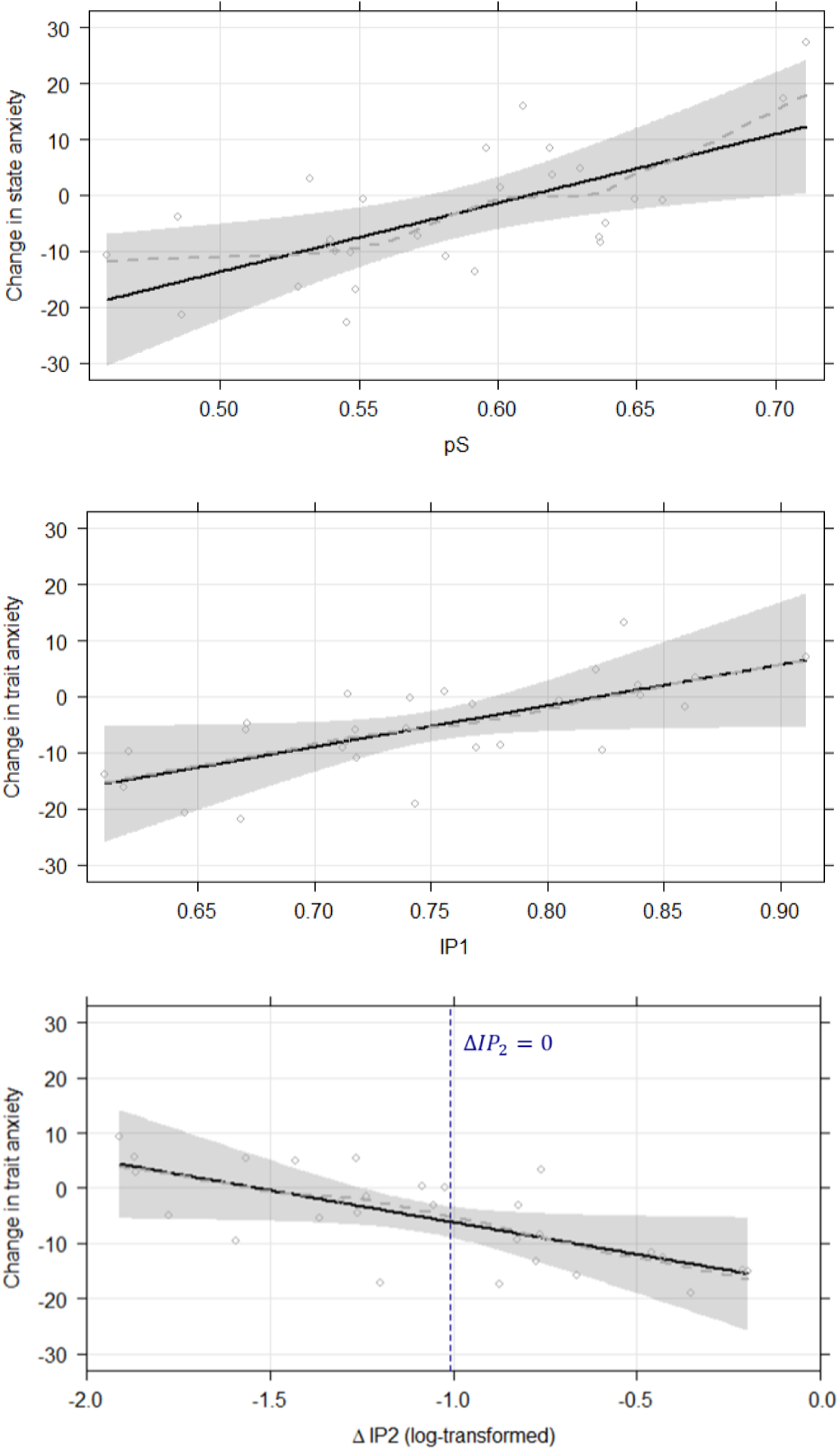
Effect plots with partial residuals for significant associations between computational parameter estimates and change in anxiety in the interoceptive training group, as estimated by linear mixed effects models. (Top) Lower values of *pS*, representing less biased perceptual priors towards ‘in sync’ percepts, predicted greater state anxiety reduction. (Middle) Lower values of *IP*_1_, representing noisier interoceptive signals (and less effective learning) predicted greater trait anxiety reductions. (Bottom) Greater increases in interoceptive precision weighting during training (Δ*IP*_2_) predicted greater trait anxiety reduction. The vertical dashed line indicates Δ*IP*_2_ = 0 (−1.05 on the log-transformed axis), with increased interoceptive precision weighting on the right and decreased interoceptive precision weighting on the left.

The LME predicting change in trait anxiety in the training group indicated that greater *IP*_1_values were significantly associated with greater increases in trait anxiety (*β* = 73.41, *SE* = 34.47, *t*(21.00) = 2.13, *p* = .045; **Figure 9, middle**). Conversely, greater Δ*IP*_2_ values (log-transformed) predicted greater reductions in trait anxiety (i.e., or smaller increases; *β* = - 11.59, *SE* = 5.41, *t*(21.00) = -2.14, *p* = .044; **Figure 9, bottom; Supplementary Results 6.1.4**). As expected, greater baseline trait anxiety was also associated with greater decreases in trait anxiety (*β* = -0.26, *SE* = 0.11, *t*(21.00) = -2.26, *p* = .035).

Ridge regression results confirmed coefficient estimates in the same direction for significant effects in both LMEs, but only the effect for *pS* was significant in ridge regression (*p* = .046; **Supplementary Results 6.1.5 – 6.1.6**). This is likely due to the reduced statistical power afforded by this more conservative approach. However, these findings suggest results should be interpreted with some caution.

When pooling both the training and control groups, an LME including all computational predictors indicated that greater reductions (or smaller increases) in trait anxiety were again associated with greater Δ*IP*_2_ values (*β* = -3.72, *SE* = 1.82, *t*(46.00) = -2.04, *p* = .047; **Supplementary Results 6.2.2**). A similar LME found no significant associations with state anxiety change in the two groups when pooled (**Supplementary Results 6.2.1**).

In contrast, the control group alone showed no significant effects in multiple regression models to explain trait or state anxiety reduction using computational parameters (**Supplementary Results 6.3**). Multiple regression models were used here instead of LMEs, as no random effects structures could be found that produced a converging and non-singular fit.

Conventional (model-free) measures of interoceptive task performance were associated with neither state nor trait anxiety change, when considering the training group only, the control group only, or both groups pooled. That is, LMEs (or multiple regressions when no viable mixed effects structure was found) controlling for age, sex, and baseline anxiety levels found no significant associations with the change in heartbeat counting accuracy, heartbeat discrimination *d’*, or *C* (**Supplementary Results 7.1 – 7.3**).

Parameter estimates that were derived separately from each assessment session showed no significant associations with anxiety reduction, whether using the change from baseline to final assessments or the baseline parameter values only (all *p*s > .24).

### Self-reported interoception

An LME testing possible effects of group, time, and their interaction (while controlling for age and sex) on MAIA total score found a significant group by time interaction (*β* = 1.28, *SE* = 0.43, *t*(52) = 3.01, *p* = .004; **Supplementary Results 8.1**). Follow-up contrasts indicated that the training group showed significant gains in MAIA total score from baseline (*EM* = 21.4, *SE* = 0.92) to final timepoints (*EM* = 23.8, *SE* = 0.92; *t*(52) = 4.08, *p* < .001), while the control group did not (baseline *EM* = 23.1, *SE* = 0.95; final *EM* = 22.9, *SE* = 0.95; *t*(52) = -0.25, *p* = .80)

To better understand this relationship with MAIA total scores, analogous post-hoc (uncorrected) LMEs were subsequently conducted on each MAIA subscale for interpretive purposes. Here we found significant group by time interactions for Noticing (*β* = 0.26, *SE* = 0.11, *t*(52) = 2.25, *p* = .028; **Supplementary Results 8.2.1**), Emotional Awareness (*β* = 0.24, *SE* = 0.10, *t*(52) = 2.44, *p* = .018; **Supplementary Results 8.2.5**), and Self-Regulation scores (*β* = 0.29, *SE* = 0.12, *t*(52) = 2.47, *p* = .017; **Supplementary Results 8.2.6**). Only random intercepts for each participant were retained in the random-effects structure to produce converging and non-singular model fits.

Follow-up contrasts indicated that the training group showed significant gains in Noticing from baseline (*EM* = 3.17, *SE* = 0.18) to final timepoints (*EM* = 3.63, *SE* = 0.18; *t*(52) = 2.94, *p* = .005), while the control group did not (baseline *EM* = 3.04, *SE* = 0.19; final *EM* = 2.99, *SE* = 0.19; *t*(52) = -0.29, *p* = .77). Similarly, the training group showed significant gains in Emotional Awareness (baseline *EM* = 3.15, *SE* = 0.19; final *EM* = 3.50, *SE* = 0.19; *t*(52) = 2.53, *p* = .01), while the control group did not (baseline *EM* = 3.22, *SE* = 0.20; final *EM* = 3.09, *SE* = 0.20; *t*(52) = -0.95, *p* = .35). Finally, the training group showed significant increases in Self-Regulation (baseline *EM* = 2.60, *SE* = 0.21; final *EM* = 3.06, *SE* = 0.21; *t*(52) = 2.77, *p* = .008), while the control group did not (baseline *EM* = 2.85, *SE* = 0.22; final *EM* = 2.72, *SE* = 0.22; *t*(52) = -0.77, *p* = .45).

#### Relationships with computational parameter estimates

Exploratory follow-up correlations revealed that model parameters also showed some correspondence with MAIA scores. In the training group, the change in MAIA total score was positively associated with *pS* (*r*(26) = .45, *p* = .017). Post-hoc correlations with subscales indicated that this relationship to total scores was best explained by greater *pS* in those with decreased Noticing (*r*(26) = -.41, *p* = .030) and decreased Trusting (*r*(26) = -.40, *p* = .033) at the final assessment. In other words, participants who overly endorsed that the tones were in sync with heartbeats also subsequently noticed and trusted bodily signals less after interoceptive training (**Figure 10**). These effects should be interpreted as hypothesis-generating, given that these analyses were exploratory.

**Figure 10.**
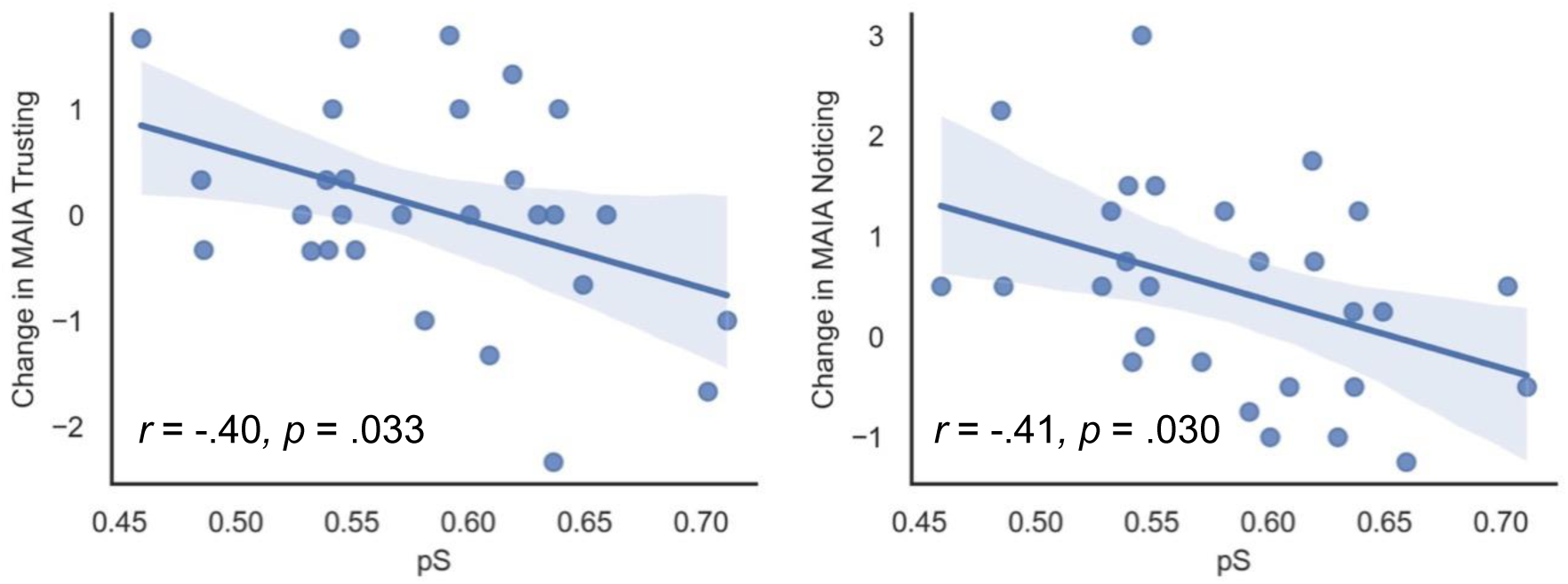
Scatterplots showing the association in the training group between estimates of *pS* and change in self-reported MAIA Trusting (left) and Noticing (right) subscales from baseline to final timepoints. MAIA = Multidimensional Assessment of Interoceptive Awareness.

#### Relationships with anxiety reduction

In the training group, similar exploratory correlations indicated that changes in trait anxiety were positively correlated with changes in MAIA total score (*r*(26) = .47, *p* = .010). Post-hoc correlations with subscales indicated that this relationship with total scores was best explained by increases in Attention Regulation in those with reductions in trait anxiety (*r*(26) = -.42, *p* = .026). In both groups pooled, changes in trait anxiety were again positively correlated with change in MAIA total score (*r*(52) = .31, *p* = .025), but post-hoc correlations with subscales indicated no significant associations. Again, these effects should be interpreted as exploratory and hypothesis-generating.

## Discussion

This study implemented a structured form of cardiac interoceptive training, which significantly enhanced interoceptive accuracy and reduced both trait and state anxiety. A novel computational modelling approach tested several competing hypotheses about the mechanisms that underpin interoceptive learning, and identified computational phenotypes that may explain individual variation in anxiety reduction, over and above conventional interoceptive task measures.

The computational model that was most supported by the data posited that cardiac perception involves combining afferent interoceptive signals (weighted by one’s internal estimate of their reliability or precision) with prior biases to produce (Bayesian) posterior beliefs that determine one’s response on each trial of the heartbeat discrimination task. Interoceptive learning under this formulation therefore involves increasing (beliefs about) the precision weighting that should be assigned to afferent signals. Furthermore, noise in afferent cardiac signals themselves was assumed to vary across individuals, with resultant bottom-up effects on rate of interoceptive learning. Estimating the values of model parameters *IP*_1_ (afferent signal precision), *IP*_2_ (starting value of interoceptive precision weighting), and *pS* (prior bias) that best reproduced each participant’s responses on the heartbeat discrimination task allowed us to quantify individual differences in the above mechanisms and investigate their relationships with other measures that changed due to interoceptive training. Subsequently, the change in interoceptive precision weighting over the course of training (Δ*IP*_2_) was derived using these parameter estimates and the model equations that govern learning.

Associations found between computational variables (*IP*_1_ and Δ*IP*_2_) and improvements in heartbeat discrimination task performance support the model’s validity in explaining the dynamics that underpin learning during the training sessions. Furthermore, the association between *IP*_1_ and mean heartrate across heartbeat discrimination trials supports the assumption made in the model that *IP*_1_ represents a latent cardiovascular trait associated with afferent signal noise. Computational parameters (*IP*_1_, *IP*_2_, and *pS*) showed differential patterns of association with confusion matrix indices of task responses, thus appearing to interact to produce the overall pattern of behaviour.

### Explaining treatment response

Computational variables explained individual variation in anxiety reduction in participants who received interoceptive training, while conventional interoceptive task measures did not. Specifically, state anxiety reduction was explained by the parameter *pS*, such that participants with more balanced prior beliefs (i.e., values of *pS* closer to .5) derived a greater anxiolytic benefit from the interoceptive training. In contrast, participants with the strongest prior biases (towards believing that the stimuli are synchronous) tended to have worsened state anxiety following training. Therefore, biased prior beliefs for interoceptive-exteroceptive integration represent a potential contraindication for interoceptive training.

Trait anxiety reduction was explained by Δ*IP*_2_, such that participants with the most enhanced interoceptive precision weighting also showed the greatest reduction in trait anxiety. **Figure 9 (bottom)** illustrates that participants whose interoceptive precision weighting increased (Δ*IP*_2_ > 0 or log-transformed Δ*IP*_2_ > −1.05) tended to also show alleviated trait anxiety, while those whose interoceptive precision weighting diminished (Δ*IP*_2_ < 0 or log-transformed Δ*IP*_2_ < −1.05) tended to show worsened trait anxiety. On the other hand, participants with the least reliable cardiac afferent signals (lowest values of *IP*_1_; slowest learning) tended to show the greatest reduction in trait anxiety (**Figure 9, middle**). The direction of this effect may initially appear surprising, given the strong positive association between *IP*_1_ and Δ*IP*_2_. However, since these are partial regression effects (from linear mixed models), they represent the effect of *IP*_1_ while holding Δ*IP*_2_ (and all other predictors) constant, and vice versa. It should also be noted that slower learning could lead to smoother convergence onto more stably increased precision weightings across the training.

Overall, these results present evidence for novel computational mechanisms that could explain anxiolytic responses to interoceptive training. However, this study could not identify computational phenotypes or interoceptive measures derived solely from the baseline assessment session that prospectively predicted subsequent anxiety reduction. This may be due to the lack of feedback during assessment sessions, which could have hindered estimation of parameter values that interact to characterise learning as well as perception.

Future work could therefore focus on designing a screening procedure of economical length using the heartbeat discrimination task (ideally with feedback) to prospectively predict treatment response. Success here could allow for personalised treatment allocation of interoceptive training as an intervention for anxiety.

### Mechanisms of interoceptive learning and anxiety

Our computational modelling approach provided novel insights about the mechanisms that may underlie interoceptive learning, with potential implications for understanding the role of interoceptive disruptions in psychopathology, clinical applications, and future neurocognitive research. Firstly, the present findings lend additional empirical support to Bayesian accounts of interoceptive psychopathology, which propose that anxious symptoms arise from maladaptively low interoceptive precision weighting (i.e., such that priors dominate perception), and that ‘normalising’ precision weighting should be anxiolytic (Owens et al., 2018; Paulus & Stein, 2006), at least in the case of trait anxiety.

The present findings also provide an empirical illustration of ideas proposed within the clinical psychology literature, such as that learning to intentionally evaluate one’s interoceptive sensations and their potential causes (‘mentalizing interoception’, or using interoceptive sensations to infer one’s own emotions) can be harnessed in psychotherapy (Duquette & Ainley, 2019; Smith et al., 2018; Smith & Lane, 2015). This idea posits that emotional states are inferred as the best explanation for a person’s interoceptive and exteroceptive cues (for supportive simulation work, see Smith, Lane, et al., 2019; Smith, Parr, et al., 2019). Accordingly, increasing the precision assigned to interoceptive signals should lead to therapeutic benefit, as supported by the present findings. Further, breaking down maladaptive prior beliefs about associations between interoceptive signals and anxious emotional states is proposed to lead to therapeutic benefit (Duquette & Ainley, 2019; Khalsa & Feinstein, 2018). The present intervention may have leveraged a similar mechanism by allowing participants to attend to autonomic arousal in a non-threatening context (self-paced physical activity). This account is supported by associations found in exploratory analyses between anxiety reduction and reduced habitual worrying about bodily sensations, and an increased capacity to attend to them, indexed by the MAIA subscales for Not Worrying and Attention Regulation. Computational parameter estimates were also found to be associated with changes in self-reported interoception. Notably, greater estimates of *pS* (i.e., more biased prior beliefs towards synchronous observations) were correlated with *reduced* scores on the MAIA Trusting and Noticing subscales, again suggesting that biased prior beliefs are a potential contraindication for interoceptive training.

The second mechanistic insight offered by our computational modelling approach is that interoceptive learning was best explained by a hierarchical Bayesian model, which allowed cardiac observations to have varying degrees of noise that affected learning, which was controlled by the parameter *IP*_1_. Importantly, greater *IP*_1_ estimates (i.e., less noisy, more precise afferent cardiac signals) were strongly associated with greater heartbeat discrimination task accuracy improvement due to training. This result suggests that learning to detect a signal (and learn from it) depends on the quality or precision of the signal itself, which may differ between individuals.

Future research should investigate the (peripheral or central) neural or physiological correlates of *IP*_1_. The present findings suggest that *IP*_1_ could reflect individual differences in cardiovascular function that result in varying noise in the afferent cardiac signal. That said, *IP*_1_ effectively controls the rate of evidence accumulation or learning, and so may plausibly relate to changes in plasticity as well, such as nucleus tractus solitaris function, consistent with Allen and colleagues’ (2022) proposal for neural circuits supporting interoceptive inference. Under this model, afferent signals from baroreceptors arrive via the vagus nerve to the nucleus tractus solitaris, which encodes the observed cardiac outcomes before passing inferred interoceptive states to the posterior insula. Inferred states are then thought to be passed on as observations to the anterior insular cortex, which is a candidate for implementing the higher level in our computational model (as also supported by computational architectures posited within neurovisceral integration theory; see Smith et al., 2017). To elucidate the neural and/or physiological basis of the computational model from this study, future research should measure potential cardiovascular correlates of baroreceptor activation during the heartbeat discrimination task (e.g., blood pressure, pulse wave amplitude, stroke volume), as well as task-related brain activity onto which the computational parameters can be regressed. Hypertensive patients offer a particularly promising clinical group for this research, as they present with chronic alterations to cardiovascular function, along with both lower heartbeat tracking accuracy and attenuated heartbeat-evoked potentials (i.e., where electrophysiological brain responses are time-locked to individual heartbeats; Yoris et al., 2018).

While the winning computational model used in this study posited that *IP*_1_ is a static individual difference, we also applied this model to data from the assessment sessions for both groups, producing estimates of *IP*_1_ that varied within-subjects between the baseline, midpoint, and final assessments. This illustrates two approaches for quantifying static vs. dynamic facets of afferent signal precision to identify its physiological correlates. One might speculate that, while there is plausibly a static physiological component of *IP*_1_, individuals could feasibly learn to enhance *IP*_1_ in a state-like manner, perhaps by altering their breathing to amplify heartbeat sensations. Supporting this, breath-holding has been demonstrated to increase cardiac interoceptive precision in related computational modelling studies (Smith, Kuplicki, Feinstein, Forthman, Stewart, Paulus, Tulsa 1000 investigators, et al., 2020; Smith, Kuplicki, Teed, et al., 2020) This is consistent with ’active’ interoceptive inference theory (Allen et al., 2022), which posits that agents act to improve the precision of incoming observations, in the aim of minimising prediction errors. That said, given moderate levels of recoverability and fewer available trials in these separate sessions, it is important to acknowledge that some differences could also reflect estimation error.

Our results also further link interoception to anxiety. Prior cross-sectional research has highlighted a complex relationship between these constructs, where results appear to be influenced by method of interoceptive testing (Domschke et al., 2010; Garfinkel et al., 2016), the nature of the anxiety disorder, and the presence of other comorbidities (Dunn et al., 2010). Indeed, a recent meta-analysis of 55 studies found no cross-sectional association between interoceptive accuracy measures and either trait or state anxiety (Adams et al., 2022). The current work instead provides evidence that intervening on cardiac interoceptive accuracy can have an anxiolytic effect and, to our knowledge, is the first interventional study to do so in a subclinical sample with a control group, complementing findings from a prior randomised controlled trial in a sample of autistic adults (Quadt et al., 2021), and a smaller study without a control group (Sugawara et al., 2020). The present findings also suggest that gains in interoceptive processing were achieved within four training sessions, as indicated by both computational and conventional accuracy measures. As such, future iterations of interoceptive training would likely benefit from reducing the number of training sessions.

The computational modelling approach in this study has some limitations to consider. For example, while the winning model was supported as having the most evidence within model comparison, and its robustness confirmed by recoverability and identifiability analyses, protected exceedance probabilities were not definitive and other modelling approaches could have been considered. The model also did not explicitly account for possible differences in perceptual processing of the tone stimulus in the task (i.e., it modelled the heartbeat and tone as a single combined observation), This could be relevant, as the heartbeat discrimination task assesses interoceptive-exteroceptive integration. An alternative model setup could explicitly include both cardiac and auditory signals that are jointly used to infer higher-level states. The approach taken in this study effectively assumed that auditory signals in the trial were perfectly precise (as supported by related modelling work; see Smith, Kuplicki, Feinstein, Forthman, Stewart, Paulus, Tulsa 1000 investigators, et al., 2020), such that relevant precisions in the model concerned only cardiac sensations and their integration with auditory signals. Nevertheless, building on recent single-level models (Smith, Kuplicki, Feinstein, Forthman, Stewart, Paulus, Tulsa 1000 investigators, et al., 2020; Smith, Mayeli, et al., 2021), this study presents the first use, to our knowledge, of hierarchical Bayesian modelling to characterise interoceptive learning and identify a computational phenotype that captures response to an interoceptive training intervention.

Overall, with these considerations in mind, the present findings support a growing body of evidence that interoceptive processes contribute to mental health symptoms (Khalsa et al., 2018). They also highlight the potential for novel interoceptive therapies for mental health conditions (Nord & Garfinkel, 2022). This is also the first study to elaborate on mechanisms of an interoception-based intervention using computational modelling, explaining treatment responses and underlying mechanisms. To validate the utility of this approach in the clinical management of anxiety, this behavioural intervention should be extended to patient populations, with the view to develop computational phenotypes that can guide personalised treatment allocation.

## Supporting information

Supplementary Results

## Acknowledgements

This work was supported by MQ (PsyImpact Grant awarded to HDC), the University of Sussex MSc programmes, and via a donation from the Dr. Mortimer and Theresa Sackler Foundation. CS is supported by a Wellcome Four-year PhD Studentship in Science (UCL Wellcome 4-year PhD in Mental Health Science).

