## Supplementary Results for "A hierarchical Bayesian model reveals increased precision weighting for afferent cardiac signals, and reduced anxiety, as a function of interoceptive training"

### 1 Computational modelling

#### 1.1 Parameter recovery results

|  | **Parameter** | **Pearson *r*(26)** | ***p* value** |
| --- | --- | --- | --- |
| **Model 1** | ${IP}_{2}$ | .98 | < .001 |
| **Model 2** | ${IP}_{2}$ | .99 | < .001 |
|  | $pS$ | .83 | < .001 |
| **Model 3** | ${IP}_{2}$ | .84 | < .001 |
|  | $\eta$ | .98 | < .001 |
| **Model 4** | ${IP}_{2}$ | .81 | < .001 |
|  | $pS$ | .93 | < .001 |
|  | $\eta$ | .98 | < .001 |
| **Model 5** | ${IP}_{1}$ | .97 | < .001 |
|  | ${IP}_{2}$ | .98 | < .001 |
| **Model 6** | ${IP}_{1}$ | .98 | < .001 |
|  | ${IP}_{2}$ | .98 | < .001 |
|  | $pS$ | .93 | < .001 |
| **Model 7** | ${IP}_{1}$ | .97 | < .001 |
|  | ${IP}_{2}$ | .78 | < .001 |
| **Model 8** | ${IP}_{1}$ | .97 | < .001 |
|  | ${IP}_{2}$ | .41 | .031 |
|  | $pS$ | .93 | < .001 |
| **Model 9** | ${IP}_{1}$ | .96 | < .001 |
|  | ${IP}_{2}$ | .83 | < .001 |
|  | $pS$ | .93 | < .001 |
|  | $\omega$ | .28 | .145 |
| **Model 10** | ${IP}_{1}$ | .97 | < .001 |
|  | ${IP}_{2}$ | .81 | < .001 |
|  | $pS$ | .92 | < .001 |
|  | $\omega_{Block}$ | -.23 | .237 |
| **Model 11** | ${IP}_{1}$ | .96 | < .001 |
|  | ${IP}_{2}$ | .81 | < .001 |
|  | $pS$ | .93 | < .001 |
|  | ${IP}_{1 Diff}$ | .74 | < .001 |
| **Model 12** | ${IP}_{1}$ | .98 | < .001 |
|  | ${IP}_{2}$ | .77 | < .001 |
|  | $pS$ | .96 | < .001 |
|  | $\eta_{\mathbf{D}}$ | .93 | < .001 |
| **Model 13** | ${IP}_{1}$ | .97 | < .001 |
|  | ${IP}_{2}$ | .67 | < .001 |
|  | $pS$ | .81 | < .001 |
|  | $\zeta$ | .90 | < .001 |

**Legend.** ${IP}_{1}$: lower-level signal precision; ${IP}_{2}$: interoceptive precision weighting; $\eta$: learning rate for interoceptive precision weighting; $pS$: prior bias; $\omega$: inverse forgetting rate across trials; $\omega_{Block}$: inverse forgetting rate across training sessions; $\eta_{\mathbf{D}}$: learning rate for prior bias; $\zeta$: precision of beliefs about state transitions (‘faulty memory’ mechanism).

#### 1.2 Model identifiability results

|  | **Identifiability winning model** | **pxp** | **Included in final model comparison** |
| --- | --- | --- | --- |
| **Model 1** | 5 | 1.00 | N |
| **Model 2** | 6 | .94 | N |
| **Model 3** | 7 | 1.00 | N |
| **Model 4** | 8 | .79 | N |
| **Model 5** | 5 | 1.00 | Y |
| **Model 6** | 6 | .90 | Y |
| **Model 7** | 7 | .90 | Y |
| **Model 8** | 8 | .90 | Y |
| **Model 9** | - | - | N |
| **Model 10** | - | - | N |
| **Model 11** | 11 | .96 | Y |
| **Model 12** | 12 | 1.00 | Y |
| **Model 13** | 7 | .56 | N |

**Legend.** Identifiability analysis results for models that survived parameter recovery analysis. Each model was used to generate a synthetic dataset, and all models that survived parameter recovery analysis were fit on each synthetic dataset. Y: Included in final model comparison; N: excluded from final model comparison. Models 9 and 10 were excluded from this step as they included unrecoverable parameters. A model was deemed identifiable and included in final model comparison if Bayesian model comparison that the model produced the best fit on the synthetic dataset generated by itself. pxp: protected exceedance probability.

### 2 Relationship between $\boldsymbol{IP}_{\boldsymbol{1}}$ and heartrate across heartbeat discrimination trials

#### 2.1 In both groups pooled

Multiple regression was used for this analysis as no random effects structure for a linear mixed effects model could be found that produced a non-singular fit.

*Multiple Regression to Predict* ${IP}_{1}$ *Parameter Estimates*

| *Predictors* | *Estimate* | *Std. Error* | *Statistic* | *p* | *df* |
| --- | --- | --- | --- | --- | --- |
| (Intercept) | 1.194 | 0.146 | 8.199 | **<0.001** | 47.000 |
| Mean Heart Rate | -0.003 | 0.001 | -2.869 | **0.006** | 47.000 |
| Mean SD in Heart Rate | -0.005 | 0.004 | -1.229 | 0.225 | 47.000 |
| ${IP}_{2}$ | -0.104 | 0.103 | -1.011 | 0.317 | 47.000 |
| $pS$ | -0.069 | 0.089 | -0.772 | 0.444 | 47.000 |
| Age | -0.002 | 0.001 | -1.487 | 0.144 | 47.000 |
| Sex [Male] | -0.018 | 0.026 | -0.687 | 0.495 | 47.000 |
| Observations | 54 | | | | |
| R^2^ / R^2^ adjusted | 0.187 / 0.084 | | | | |

*Quantile-Quantile Plot of Model Residuals*

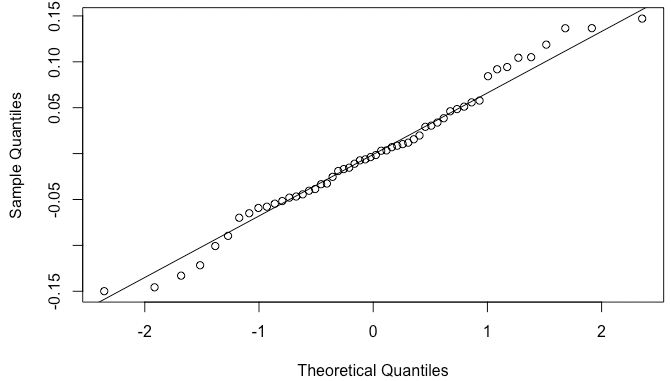

A Shapiro-Wilk test indicated that model residuals were normally distributed (*W* = .98; *p* = .60). Variance inflation factors for model predictors did not indicate multicollinearity (Mean Heart Rate: 1.04; Mean SD in Heart Rate: 1.20; ${IP}_{2}$: 1.08; $pS$: 1.14; Age: 1.22; Sex: 1.08).

#### 2.2 In training group only

Multiple regression was used for this analysis as no random effects structure for a linear mixed effects model could be found that produced a non-singular fit.

*Multiple Regression to Predict* ${IP}_{1}$ *Parameter Estimates*

| *Predictors* | *Estimate* | *Std. Error* | *Statistic* | *p* | *df* |
| --- | --- | --- | --- | --- | --- |
| (Intercept) | 1.611 | 0.229 | 7.040 | **<0.001** | 21.000 |
| Mean Heart Rate | -0.003 | 0.002 | -2.226 | **0.037** | 21.000 |
| Mean SD in Heart Rate | -0.006 | 0.006 | -0.956 | 0.350 | 21.000 |
| ${IP}_{2}$ | -0.136 | 0.112 | -1.219 | 0.237 | 21.000 |
| $pS$ | -0.718 | 0.263 | -2.726 | **0.013** | 21.000 |
| Age | -0.001 | 0.002 | -0.643 | 0.527 | 21.000 |
| Sex [Male] | -0.043 | 0.036 | -1.184 | 0.250 | 21.000 |
| Observations | 28 | | | | |
| R^2^ / R^2^ adjusted | 0.441 / 0.281 | | | | |

*Quantile-Quantile Plot of Model Residuals*

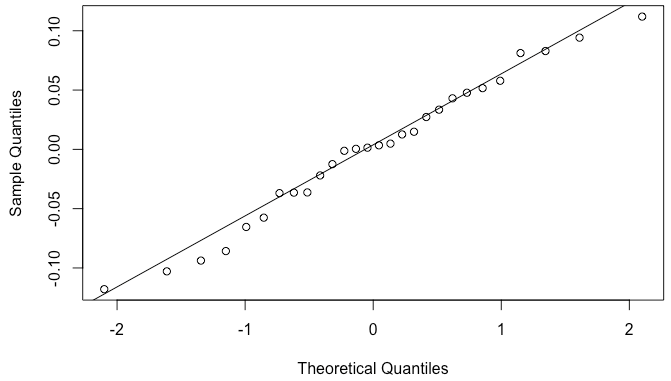

A Shapiro-Wilk test indicated that model residuals were normally distributed (*W* = .98; *p* = .84). Variance inflation factors for model predictors did not indicate multicollinearity (Mean Heart Rate: 1.35; Mean SD in Heart Rate: 1.68; ${IP}_{2}$: 1.22; $pS$: 1.55; Age: 1.73; Sex: 1.54).

### 3 Change in conventional interoceptive task measures across assessments

#### 3.1 Change in heartbeat tracking accuracy (HBT)

*Linear Mixed Effects Model to Predict Heartbeat Tracking Accuracy*

| *Predictors* | *Estimates* | *std. Error* | *Statistic* | *p* | *df* |
| --- | --- | --- | --- | --- | --- |
| (Intercept) | 0.59 | 0.03 | 16.85 | **<0.001** | 74.01 |
| Group | -0.03 | 0.03 | -1.02 | 0.308 | 83.60 |
| Time [Final] | 0.18 | 0.03 | 6.93 | **<0.001** | 102.01 |
| Time [Midpoint] | 0.13 | 0.03 | 4.73 | **<0.001** | 102.48 |
| Age | 0.00 | 0.00 | 0.58 | 0.561 | 49.77 |
| Sex | -0.05 | 0.03 | -1.54 | 0.131 | 49.73 |
| Group × Time [Final] | 0.12 | 0.03 | 4.58 | **<0.001** | 102.01 |
| Group × Time [Midpoint] | 0.13 | 0.03 | 4.83 | **<0.001** | 102.48 |
| **Random Effects** | | | | | |
| σ^2^ | 0.02 | | | | |
| τ_00_ _Participant_ | 0.03 | | | | |
| ICC | 0.61 | | | | |
| N _Participant_ | 54 | | | | |
| Observations | 160 | | | | |
| Marginal R^2^ / Conditional R^2^ | 0.239 / 0.706 | | | | |

**Legend.** Kenward-Roger approximation was used for degrees of freedom and *p*-values. The marginal R-squared accounts for the variance of the fixed effects only, while the conditional R-squared accounts for both the fixed and random effects. σ^2^: Residual variance; τ_00_: variance for this random intercepts factor; ICC: intra-class correlation for this random intercepts factor; N: number of levels for this random intercepts factor.

*Quantile-Quantile Plot of Model Residuals*

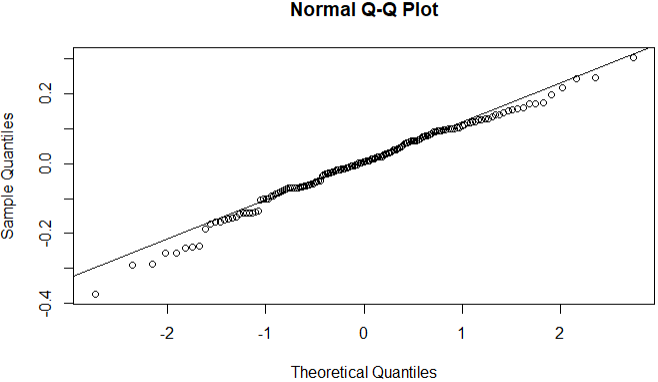

A Shapiro-Wilk test indicated that model residuals were normally distributed (*W* = .99; *p* = .13).

#### 3.2 Change in heartbeat discrimination sensitivity (*d’*)

*Linear Mixed Effects Model to Predict Heartbeat Discrimination Sensitivity*

| *Predictors* | *Estimates* | *std. Error* | *Statistic* | *p* | *df* |
| --- | --- | --- | --- | --- | --- |
| (Intercept) | 0.35 | 0.15 | 2.33 | **0.022** | 82.78 |
| Group | -0.23 | 0.13 | -1.77 | 0.080 | 95.37 |
| Time [final] | 0.74 | 0.13 | 5.87 | **<0.001** | 102.31 |
| Time [midpoint] | 0.53 | 0.13 | 4.18 | **<0.001** | 102.63 |
| Age | -0.02 | 0.01 | -1.16 | 0.250 | 49.70 |
| Sex | 0.01 | 0.13 | 0.07 | 0.942 | 49.66 |
| Group × Time [final] | 0.61 | 0.13 | 4.83 | **<0.001** | 102.31 |
| Group × Time [midpoint] | 0.58 | 0.13 | 4.54 | **<0.001** | 102.63 |
| **Random Effects** | | | | | |
| σ^2^ | 0.42 | | | | |
| τ_00_ _Participant_ | 0.47 | | | | |
| ICC | 0.53 | | | | |
| N _Participant_ | 54 | | | | |
| Observations | 160 | | | | |
| Marginal R^2^ / Conditional R^2^ | 0.198 / 0.621 | | | | |

**Legend.** Kenward-Roger approximation was used for degrees of freedom and *p*-values. The marginal R-squared accounts for the variance of the fixed effects only, while the conditional R-squared accounts for both the fixed and random effects. σ^2^: Residual variance; τ_00_: variance for this random intercepts factor; ICC: intra-class correlation for this random intercepts factor; N: number of levels for this random intercepts factor.

*Quantile-Quantile Plot of Model Residuals*

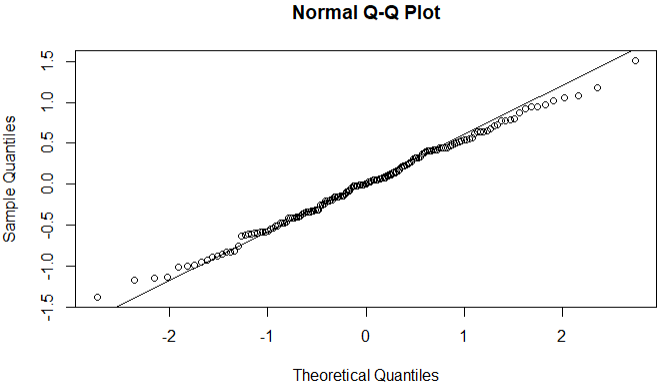

A Shapiro-Wilk test indicated that model residuals were normally distributed (*W* = 1.00; *p* = .94).

#### 3.3 Change in heartbeat discrimination criterion (*C*)

*Linear Mixed Effects Model to Predict Heartbeat Discrimination Criterion*

| *Predictors* | *Estimates* | *std. Error* | *Statistic* | *p* | *df* |
| --- | --- | --- | --- | --- | --- |
| (Intercept) | 0.01 | 0.06 | 0.21 | 0.835 | 59.85 |
| Group | -0.02 | 0.06 | -0.33 | 0.741 | 53.67 |
| Time [final] | -0.07 | 0.05 | -1.35 | 0.180 | 102.15 |
| Time [midpoint] | -0.14 | 0.05 | -2.67 | **0.009** | 102.31 |
| Age | 0.00 | 0.00 | 0.26 | 0.794 | 27.84 |
| Sex | -0.01 | 0.03 | -0.23 | 0.820 | 28.27 |
| Group × Time [final] | 0.01 | 0.05 | 0.13 | 0.900 | 102.15 |
| Group × Time [midpoint] | -0.08 | 0.05 | -1.42 | 0.157 | 102.31 |
| **Random Effects** | | | | | |
| σ^2^ | 0.07 | | | | |
| τ_00_ _Participant_ | 0.05 | | | | |
| τ_11_ _Participant.Group_ | 0.06 | | | | |
| ρ_01_ _Participant_ | -0.99 | | | | |
| ICC | 0.59 | | | | |
| N _Participant_ | 54 | | | | |
| Observations | 160 | | | | |
| Marginal R^2^ / Conditional R^2^ | 0.036 / 0.60 | | | | |

**Legend.** Kenward-Roger approximation was used for degrees of freedom and *p*-values. The marginal R-squared accounts for the variance of the fixed effects only, while the conditional R-squared accounts for both the fixed and random effects. σ^2^: Residual variance; τ_00_: variance for this random intercepts factor; τ_11_: variance for this random slopes factor; ρ_01_: random slope-intercept correlation; ICC: intra-class correlation for this random intercepts factor; N: number of levels for this random intercepts factor.

*Quantile-Quantile Plot of Model Residuals*

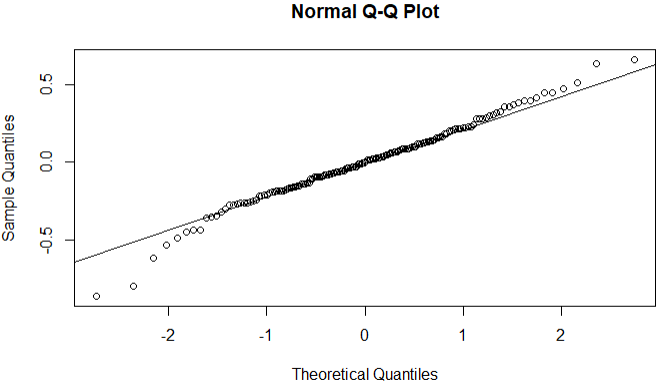

A Shapiro-Wilk test indicated that model residuals were normally distributed (*W* = .98; *p* = .06).

### 4 Change in computational parameter estimates across assessments

#### 4.1 Change in ${IP}_{1}$

*Linear Mixed Effects Model to Predict* ${IP}_{1}$ *Parameter Estimates*

| *Predictors* | *Estimates* | *Std. Error* | *Statistic* | *p* | *df* |
| --- | --- | --- | --- | --- | --- |
| (Intercept) | 0.73 | 0.01 | 70.01 | **<0.001** | 94.10 |
| Group | -0.01 | 0.01 | -1.12 | 0.267 | 109.46 |
| Time [Final] | 0.07 | 0.01 | 6.69 | **<0.001** | 103.02 |
| Time [Midpoint] | 0.05 | 0.01 | 4.64 | **<0.001** | 103.39 |
| Sex | 0.00 | 0.01 | 0.22 | 0.828 | 49.77 |
| Age | -0.00 | 0.00 | -1.31 | 0.195 | 49.79 |
| Group × Time [Final] | 0.04 | 0.01 | 4.08 | **<0.001** | 103.02 |
| Group × Time [Midpoint] | 0.04 | 0.01 | 3.76 | **<0.001** | 103.39 |
| **Random Effects** | | | | | |
| σ^2^ | 0.00 | | | | |
| τ_00_ _Participant_ | 0.00 | | | | |
| ICC | 0.41 | | | | |
| N _Participant_ | 54 | | | | |
| Observations | 161 | | | | |
| Marginal R^2^/ Conditional R^2^ | 0.242 / 0.557 | | | | |

**Legend.** Kenward-Roger approximation was used for degrees of freedom and *p*-values. The marginal R-squared accounts for the variance of the fixed effects only, while the conditional R-squared accounts for both the fixed and random effects. σ^2^: Residual variance; τ_00_: variance for this random intercepts factor; ICC: intra-class correlation for this random intercepts factor; N: number of levels for this random intercepts factor.

*Quantile-Quantile Plot of Model Residuals*

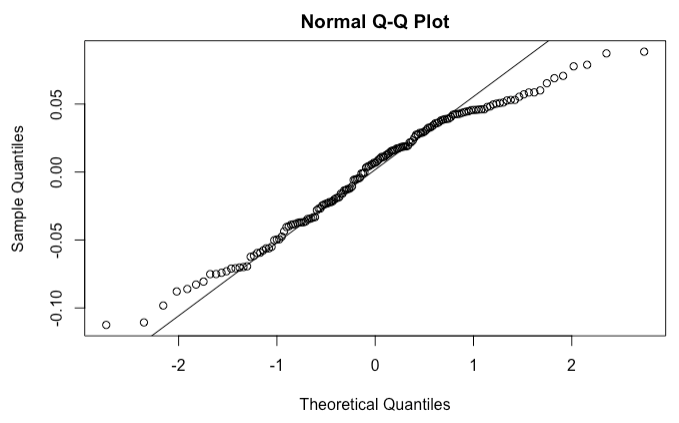

A Shapiro-Wilk test indicated that model residuals were not normally distributed (*W* = .97; *p* = .003).

#### 4.2 Change in ${IP}_{2}$

*Linear Mixed Effects Model to Predict* ${IP}_{2}$ *Parameter Estimates*

| *Predictors* | *Estimates* | *Std. Error* | *Statistic* | *p* | *df* |
| --- | --- | --- | --- | --- | --- |
| (Intercept) | 0.74 | 0.01 | 139.33 | **<0.001** | 101.97 |
| Group | -0.01 | 0.00 | -2.35 | **0.021** | 118.58 |
| Time [Final] | 0.02 | 0.01 | 4.62 | **<0.001** | 103.03 |
| Time [Midpoint] | 0.03 | 0.01 | 5.10 | **<0.001** | 103.45 |
| Sex | 0.00 | 0.00 | 0.23 | 0.821 | 49.74 |
| Age | -0.00 | 0.00 | -0.49 | 0.627 | 49.76 |
| Group × Time [Final] | 0.02 | 0.01 | 4.14 | **<0.001** | 103.03 |
| Group × Time [Midpoint] | 0.02 | 0.01 | 3.62 | **<0.001** | 103.45 |
| **Random Effects** | | | | | |
| σ^2^ | 0.00 | | | | |
| τ_00_ _Participant_ | 0.00 | | | | |
| ICC | 0.35 | | | | |
| N _Participant_ | 54 | | | | |
| Observations | 161 | | | | |
| Marginal R^2^/ Conditional R^2^ | 0.186 / 0.470 | | | | |

**Legend.** Kenward-Roger approximation was used for degrees of freedom and *p*-values. The marginal R-squared accounts for the variance of the fixed effects only, while the conditional R-squared accounts for both the fixed and random effects. σ^2^: Residual variance; τ_00_: variance for this random intercepts factor; ICC: intra-class correlation for this random intercepts factor; N: number of levels for this random intercepts factor.

*Quantile-Quantile Plot of Model Residuals*

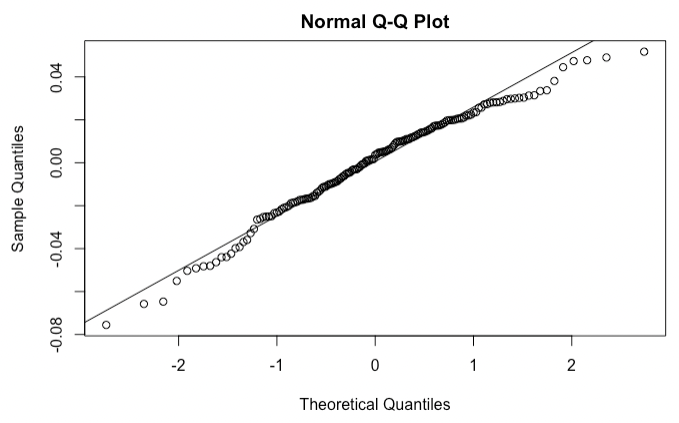

A Shapiro-Wilk test indicated that model residuals were not normally distributed (*W* = .98; *p* = .012).

#### 4.3 Change in $pS$

*Linear Mixed Effects Model to Predict* $pS$ *Parameter Estimates*

| *Predictors* | *Estimates* | *std. Error* | *Statistic* | *p* | *df* |
| --- | --- | --- | --- | --- | --- |
| (Intercept) | 0.49 | 0.02 | 28.98 | **<0.001** | 85.07 |
| Group | 0.00 | 0.01 | 0.10 | 0.918 | 98.21 |
| Time [Final] | 0.02 | 0.01 | 1.28 | 0.205 | 103.01 |
| Time [Midpoint] | 0.04 | 0.01 | 2.46 | **0.016** | 103.33 |
| Sex | 0.02 | 0.01 | 1.15 | 0.255 | 49.81 |
| Age | -0.00 | 0.00 | -0.94 | 0.353 | 49.83 |
| Group × Time [Final] | 0.00 | 0.01 | 0.20 | 0.838 | 103.01 |
| Group × Time [Midpoint] | 0.03 | 0.01 | 1.81 | 0.073 | 103.33 |
| **Random Effects** | | | | | |
| σ^2^ | 0.01 | | | | |
| τ_00_ _Participant_ | 0.01 | | | | |
| ICC | 0.50 | | | | |
| N _Participant_ | 54 | | | | |
| Observations | 161 | | | | |
| Marginal R^2^/ Conditional R^2^ | 0.062 / 0.529 | | | | |

**Legend.** Kenward-Roger approximation was used for degrees of freedom and *p*-values. The marginal R-squared accounts for the variance of the fixed effects only, while the conditional R-squared accounts for both the fixed and random effects. σ^2^: Residual variance; τ_00_: variance for this random intercepts factor; ICC: intra-class correlation for this random intercepts factor; N: number of levels for this random intercepts factor.

*Quantile-Quantile Plot of Model Residuals*

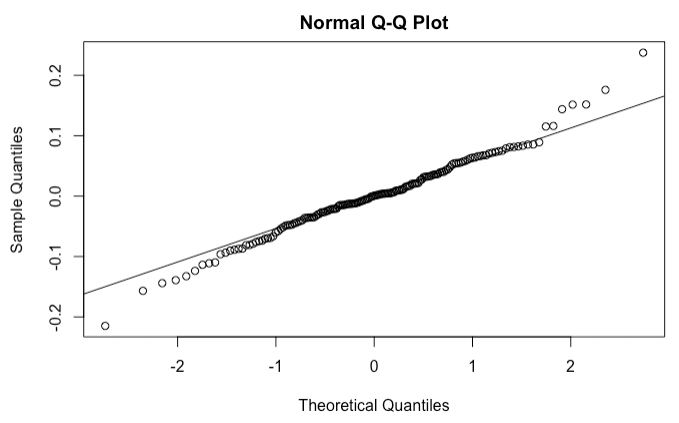

A Shapiro-Wilk test indicated that model residuals were normally distributed (*W* = .99; *p* = .08).

### 5 Change in anxiety across assessments

#### 5.1 Change in state anxiety

*Linear Mixed Effects Model to Predict State Anxiety*

| *Predictors* | *Estimates* | *Std. Error* | *Statistic* | *p* | *df* |
| --- | --- | --- | --- | --- | --- |
| (Intercept) | 37.71 | 1.70 | 22.16 | **<0.001** | 75.19 |
| Group | 1.91 | 1.50 | 1.28 | 0.205 | 83.24 |
| Time [final] | 0.01 | 1.56 | 0.01 | 0.993 | 52.00 |
| Age | 0.36 | 0.16 | 2.22 | **0.031** | 50.00 |
| Sex | 1.50 | 1.52 | 0.98 | 0.330 | 50.00 |
| Group × Time [final] | -3.37 | 1.56 | -2.15 | **0.036** | 52.00 |
| **Random Effects** | | | | | |
| σ^2^ | 65.99 | | | | |
| τ_00_ _Participant_ | 51.02 | | | | |
| ICC | 0.44 | | | | |
| N _Participant_ | 54 | | | | |
| Observations | 108 | | | | |
| Marginal R^2^ / Conditional R^2^ | 0.094 / 0.489 | | | | |

**Legend.** Kenward-Roger approximation was used for degrees of freedom and *p*-values. The marginal R-squared accounts for the variance of the fixed effects only, while the conditional R-squared accounts for both the fixed and random effects. σ^2^: Residual variance; τ_00_: variance for this random intercepts factor; ICC: intra-class correlation for this random intercepts factor; N: number of levels for this random intercepts factor.

*Quantile-Quantile Plot of Model Residuals*

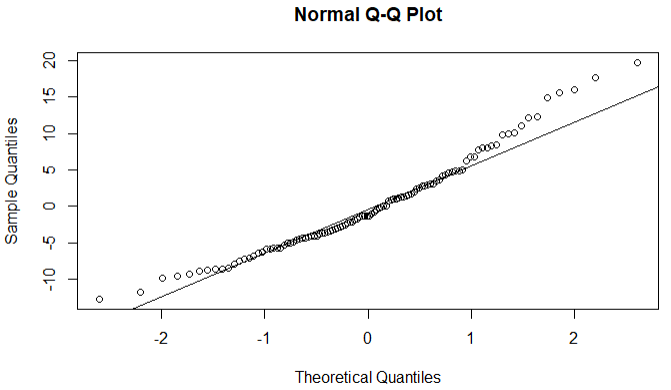

A Shapiro-Wilk test indicated that model residuals were not normally distributed (*W* = .96; *p* = .004).

#### 5.2 Change in trait anxiety

*Linear Mixed Effects Model to Predict Trait Anxiety*

| *Predictors* | *Estimates* | *Std. Error* | *Statistic* | *p* | *df* |
| --- | --- | --- | --- | --- | --- |
| (Intercept) | 43.48 | 1.73 | 25.15 | **<0.001** | 58.00 |
| Group | 1.92 | 1.48 | 1.29 | 0.201 | 61.13 |
| Time [final] | -3.04 | 0.94 | -3.23 | **0.002** | 52.00 |
| Age | 0.19 | 0.18 | 1.08 | 0.284 | 50.00 |
| Sex | 1.31 | 1.67 | 0.79 | 0.436 | 50.00 |
| Group × Time [final] | -2.35 | 0.94 | -2.49 | **0.016** | 52.00 |
| **Random Effects** | | | | | |
| σ^2^ | 23.98 | | | | |
| τ_00_ _Participant_ | 89.80 | | | | |
| ICC | 0.79 | | | | |
| N _Participant_ | 54 | | | | |
| Observations | 108 | | | | |
| Marginal R^2^ / Conditional R^2^ | 0.064 / 0.803 | | | | |

**Legend.** Kenward-Roger approximation was used for degrees of freedom and *p*-values. The marginal R-squared accounts for the variance of the fixed effects only, while the conditional R-squared accounts for both the fixed and random effects. σ^2^: Residual variance; τ_00_: variance for this random intercepts factor; ICC: intra-class correlation for this random intercepts factor; N: number of levels for this random intercepts factor.

*Quantile-Quantile Plot of Model Residuals*

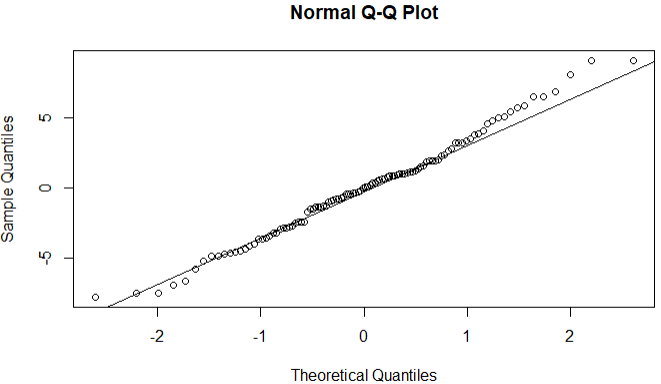

A Shapiro-Wilk test indicated that model residuals were normally distributed (*W* = .99; *p* = .56).

### 6 Relationships between change in anxiety and computational parameters

#### 6.1 In training group only

##### 6.1.1 State anxiety change

*Linear Mixed Effects Model to Predict Change in State Anxiety*

| *Predictors* | *Estimates* | *Std. Error* | *Statistic* | *p* | *df* |
| --- | --- | --- | --- | --- | --- |
| (Intercept) | -151.57 | 62.59 | -2.42 | **0.025** | 20.01 |
| ${IP}_{1}$ | 133.21 | 69.92 | 1.91 | 0.071 | 20.00 |
| ${IP}_{2}$ | -3.42 | 3.52 | -0.97 | 0.344 | 20.00 |
| ${\Delta IP}_{2}$ | -23.59 | 15.05 | -1.57 | 0.133 | 20.00 |
| $pS$ | 109.05 | 44.59 | 2.45 | **0.024** | 20.00 |
| Baseline State Anxiety | -0.70 | 0.15 | -4.61 | **<0.001** | 20.00 |
| Age | 0.10 | 0.24 | 0.42 | 0.678 | 20.00 |
| Sex | -2.84 | 2.60 | -1.09 | 0.285 | 26.67 |
| **Random Effects** | | | | | |
| σ^2^ | 98.79 | | | | |
| τ_00_ _Sex_ | 1.82 | | | | |
| ICC | 0.02 | | | | |
| N _Sex_ | 2 | | | | |
| Observations | 28 | | | | |
| Marginal R^2^ / Conditional R^2^ | 0.513 / 0.522 | | | | |

**Legend.** Kenward-Roger approximation was used for degrees of freedom and *p*-values. The marginal R-squared accounts for the variance of the fixed effects only, while the conditional R-squared accounts for both the fixed and random effects. σ^2^: Residual variance; τ_00_: variance for this random intercepts factor; ICC: intra-class correlation for this random intercepts factor; N: number of levels for this random intercepts factor.

*Quantile-Quantile Plot of Model Residuals*

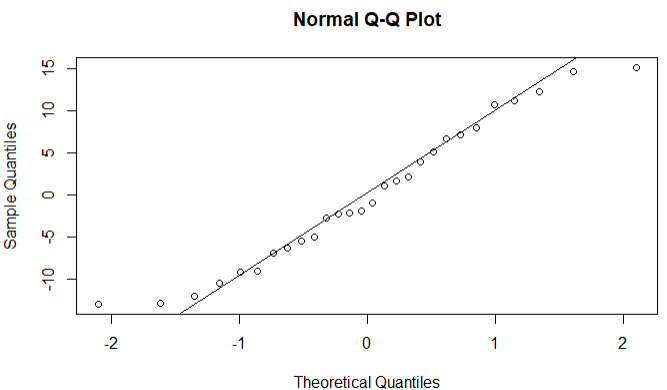

A Shapiro-Wilk test indicated that model residuals were normally distributed (*W* = .96; *p* = .32). Variance inflation factors for model predictors strongly indicated multicollinearity (${IP}_{1}$: 8.85; ${IP}_{2}$: 4.71; ${\Delta IP}_{2}$: 15.57; $pS$: 2.05; Baseline State Anxiety: 1.26; Age: 1.26; Sex: 1.30).

##### 6.1.2 Trait anxiety change

*Linear Mixed Effects Model to Predict Change in Trait Anxiety*

| *Predictors* | *Estimates* | *Std. Error* | *Statistic* | *p* | *df* |
| --- | --- | --- | --- | --- | --- |
| (Intercept) | -124.88 | 41.28 | -3.03 | **0.007** | 20.00 |
| ${IP}_{1}$ | 122.52 | 46.33 | 2.64 | **0.016** | 20.00 |
| ${IP}_{2}$ | -3.56 | 2.32 | -1.53 | 0.142 | 20.00 |
| ${\Delta IP}_{2}$ | -24.57 | 9.97 | -2.46 | **0.023** | 20.00 |
| $pS$ | 43.74 | 29.11 | 1.50 | 0.149 | 20.00 |
| Baseline Trait Anxiety | -0.27 | 0.11 | -2.45 | **0.024** | 20.00 |
| Age | 0.03 | 0.16 | 0.21 | 0.836 | 20.00 |
| Sex | -0.71 | 1.62 | -0.44 | 0.668 | 20.94 |
| **Random Effects** | | | | | |
| σ^2^ | 42.81 | | | | |
| τ_00_ _Sex_ | 0.12 | | | | |
| ICC | 0.00 | | | | |
| N _Sex_ | 2 | | | | |
| Observations | 28 | | | | |
| Marginal R^2^ / Conditional R^2^ | 0.353 / 0.355 | | | | |

**Legend.** Kenward-Roger approximation was used for degrees of freedom and *p*-values. The marginal R-squared accounts for the variance of the fixed effects only, while the conditional R-squared accounts for both the fixed and random effects. σ^2^: Residual variance; τ_00_: variance for this random intercepts factor; ICC: intra-class correlation for this random intercepts factor; N: number of levels for this random intercepts factor.

*Quantile-Quantile Plot of Model Residuals*

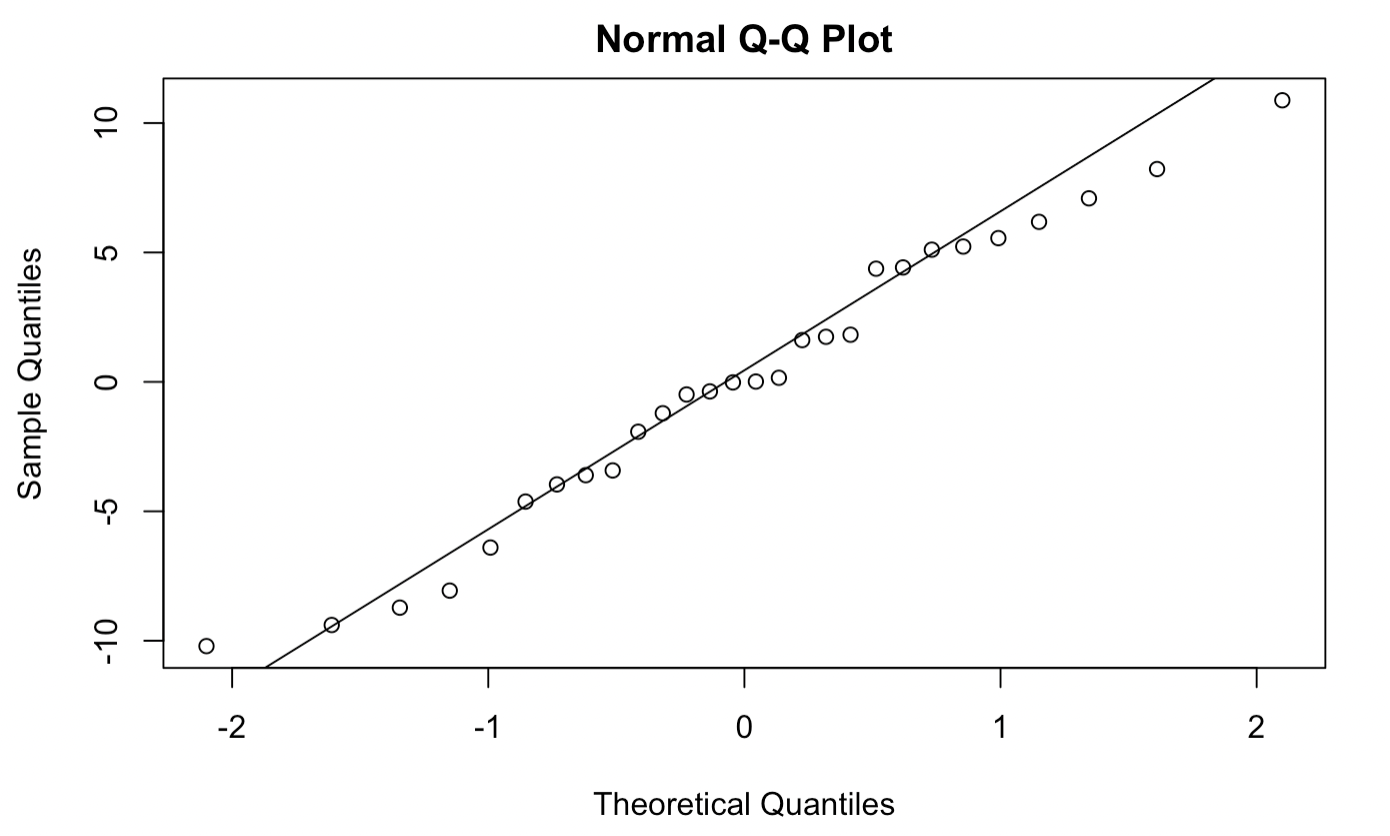

A Shapiro-Wilk test indicated that model residuals were normally distributed (*W* = .97; *p* = .72). Variance inflation factors for model predictors strongly indicated multicollinearity (${IP}_{1}$: 8.98; ${IP}_{2}$: 4.74; ${\Delta IP}_{2}$: 15.78; $pS$: 2.09; Baseline Trait Anxiety: 1.10; Age: 1.19; Sex: 1.36).

##### 6.1.3 State anxiety change (${IP}_{2}$excluded)

*Linear Mixed Effects Model to Predict Change in State Anxiety*

| *Predictors* | *Estimates* | *Std. Error* | *Statistic* | *p* | *df* |
| --- | --- | --- | --- | --- | --- |
| (Intercept) | -122.33 | 54.78 | -2.23 | **0.037** | 21.01 |
| ${IP}_{1}$ | 86.38 | 50.52 | 1.71 | 0.102 | 21.00 |
| ${\Delta IP}_{2}$ | -11.19 | 7.94 | -1.41 | 0.174 | 21.00 |
| $pS$ | 123.18 | 42.09 | 2.93 | **0.008** | 21.00 |
| Baseline State Anxiety | -0.70 | 0.15 | -4.59 | **<0.001** | 21.00 |
| Age | 0.14 | 0.24 | 0.60 | 0.553 | 21.00 |
| Sex | -3.05 | 2.59 | -1.17 | 0.250 | 28.08 |
| **Random Effects** | | | | | |
| σ^2^ | 98.51 | | | | |
| τ_00_ _Sex_ | 1.82 | | | | |
| ICC | 0.02 | | | | |
| N _Sex_ | 2 | | | | |
| Observations | 28 | | | | |
| Marginal R^2^ / Conditional R^2^ | 0.506 / 0.515 | | | | |

**Legend.** Kenward-Roger approximation was used for degrees of freedom and *p*-values. The marginal R-squared accounts for the variance of the fixed effects only, while the conditional R-squared accounts for both the fixed and random effects. σ^2^: Residual variance; τ_00_: variance for this random intercepts factor; ICC: intra-class correlation for this random intercepts factor; N: number of levels for this random intercepts factor.

*Quantile-Quantile Plot of Model Residuals*

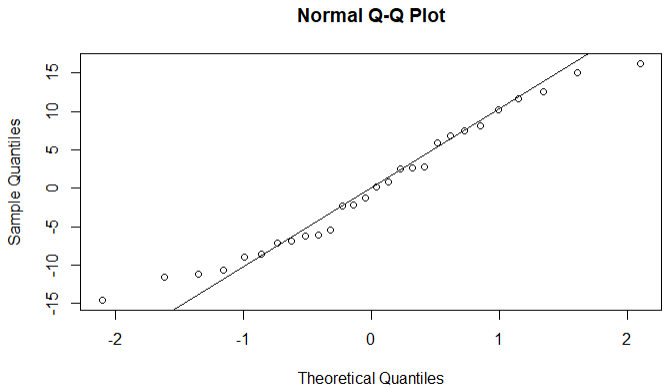

A Shapiro-Wilk test indicated that model residuals were normally distributed (*W* = .96; *p* = .40). Variance inflation factors for model predictors did not indicate multicollinearity (${IP}_{1}$: 4.63; ${\Delta IP}_{2}$: 4.35; $pS$: 1.83; Baseline State Anxiety: 1.26; Age: 1.22; Sex: 1.29).

##### 6.1.4 Trait anxiety change (${IP}_{2}$ excluded)

*Linear Mixed Effects Model to Predict Change in Trait Anxiety*

| *Predictors* | *Estimates* | *std. Error* | *Statistic* | *p* | *df* |
| --- | --- | --- | --- | --- | --- |
| (Intercept) | -94.26 | 37.24 | -2.53 | **0.019** | 21.00 |
| ${IP}_{1}$ | 73.41 | 34.47 | 2.13 | **0.045** | 21.00 |
| ${\Delta IP}_{2}$ | -11.59 | 5.41 | -2.14 | **0.044** | 21.00 |
| $pS$ | 57.98 | 28.45 | 2.04 | 0.054 | 21.00 |
| Baseline Trait Anxiety | -0.26 | 0.11 | -2.26 | **0.035** | 21.00 |
| Age | 0.08 | 0.16 | 0.49 | 0.630 | 21.00 |
| Sex | -0.90 | 1.67 | -0.54 | 0.594 | 21.99 |
| **Random Effects** | | | | | |
| σ^2^ | 45.55 | | | | |
| τ_00_ _Sex_ | 0.13 | | | | |
| ICC | 0.00 | | | | |
| N _Sex_ | 2 | | | | |
| Observations | 28 | | | | |
| Marginal R^2^ / Conditional R^2^ | 0.301 / 0.303 | | | | |

**Legend.** Kenward-Roger approximation was used for degrees of freedom and *p*-values. The marginal R-squared accounts for the variance of the fixed effects only, while the conditional R-squared accounts for both the fixed and random effects. σ^2^: Residual variance; τ_00_: variance for this random intercepts factor; ICC: intra-class correlation for this random intercepts factor; N: number of levels for this random intercepts factor.

*Quantile-Quantile Plot of Model Residuals*

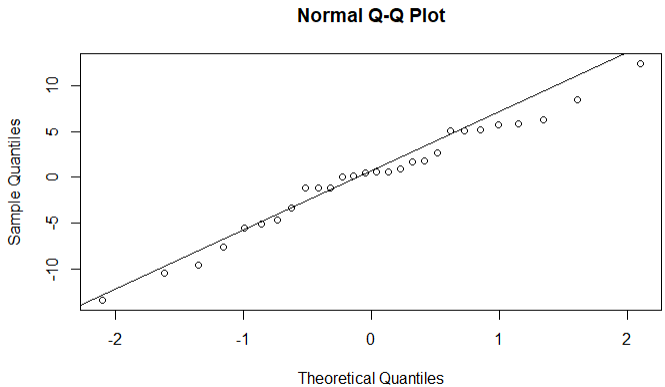

A Shapiro-Wilk test indicated that model residuals were normally distributed (*W* = .98; *p* = .78). Variance inflation factors for model predictors did not indicate multicollinearity (${IP}_{1}$: 4.67; ${\Delta IP}_{2}$: 4.37; $pS$: 1.88; Baseline Trait Anxiety: 1.09; Age: 1.15; Sex: 1.35).

##### 6.1.5 Ridge regression – state anxiety change (${IP}_{2}$ excluded)

*Ridge Regression Model to Predict Change in State Anxiety*

| *Predictors* | *Estimates* | *Scaled estimate* | *std. Error (scaled)* | *t-value (scaled)* | *p* |
| --- | --- | --- | --- | --- | --- |
| (Intercept) | -15.78 | NA | NA | NA | NA |
| ${IP}_{1}$ | 3.45 | 1.46 | 5.00 | 0.28 | .770 |
| ${\Delta IP}_{2}$ | -2.37 | -6.17 | 5.10 | 1.21 | .226 |
| $pS$ | 35.43 | 11.55 | 5.79 | **2.00** | **.046** |
| Baseline state anxiety | -0.32 | -23.49 | 6.13 | **3.83** | **.000** |
| Age | 0.02 | 0.71 | 6.16 | 0.12 | .909 |
| Sex | -0.88 | -4.20 | 6.09 | 0.69 | .491 |
| Ridge parameter | 0.80, chosen automatically, computed using 2 PCs | | | | |
| Degrees of freedom | Model: 2.85; variance: 1.61; residual: 4.09 | | | | |

##### 6.1.6 Ridge regression – trait anxiety change (${IP}_{2}$ excluded)

*Ridge Regression Model to Predict Change in Trait Anxiety*

| *Predictors* | *Estimates* | *Scaled estimate* | *std. Error (scaled)* | *t-value (scaled)* | *p* |
| --- | --- | --- | --- | --- | --- |
| (Intercept) | -9.48 | NA | NA | NA | NA |
| ${IP}_{1}$ | 0.407 | 0.17 | 1.75 | 0.10 | .922 |
| ${\Delta IP}_{2}$ | -0.87 | -2.26 | 1.78 | 1.27 | .206 |
| $pS$ | 9.19 | 3.00 | 1.92 | 1.56 | .118 |
| Baseline trait anxiety | -0.05 | -3.36 | 2.12 | 1.59 | .113 |
| Age | 0.02 | 0.86 | 2.11 | 0.41 | .684 |
| Sex | 0.03 | 0.17 | 2.05 | 0.08 | .935 |
| Ridge parameter | 2.39, chosen automatically, computed using 1 PCs | | | | |
| Degrees of freedom | Model: 1.59; variance: 0.55; residual: 2.64 | | | | |

#### 6.2 In both groups

##### 6.2.1 State anxiety change

*Linear Mixed Effects Model to Predict Change in State Anxiety*

| *Predictors* | *Estimates* | *Std. Error* | *Statistic* | *p* | *df* |
| --- | --- | --- | --- | --- | --- |
| (Intercept) | -18.47 | 23.33 | -0.79 | 0.433 | 46.02 |
| ${IP}_{1}$ | 29.27 | 26.95 | 1.09 | 0.283 | 46.00 |
| ${IP}_{2}$ | 3.48 | 2.77 | 1.26 | 0.215 | 46.00 |
| ${\Delta IP}_{2}$ | -1.89 | 2.66 | -0.71 | 0.482 | 46.00 |
| $pS$ | 6.74 | 4.01 | 1.68 | 0.100 | 46.00 |
| Baseline State Anxiety | -0.61 | 0.13 | -4.76 | **<0.001** | 46.00 |
| Age | 0.09 | 0.18 | 0.49 | 0.623 | 46.00 |
| Sex | 0.17 | 1.67 | 0.10 | 0.919 | 49.10 |
| **Random Effects** | | | | | |
| σ^2^ | 91.41 | | | | |
| τ_00_ _Sex_ | 0.18 | | | | |
| ICC | 0.00 | | | | |
| N _Sex_ | 2 | | | | |
| Observations | 54 | | | | |
| Marginal R^2^ / Conditional R^2^ | 0.403 / 0.404 | | | | |

**Legend.** Kenward-Roger approximation was used for degrees of freedom and *p*-values. The marginal R-squared accounts for the variance of the fixed effects only, while the conditional R-squared accounts for both the fixed and random effects. σ^2^: Residual variance; τ_00_: variance for this random intercepts factor; ICC: intra-class correlation for this random intercepts factor; N: number of levels for this random intercepts factor.

*Quantile-Quantile Plot of Model Residuals*

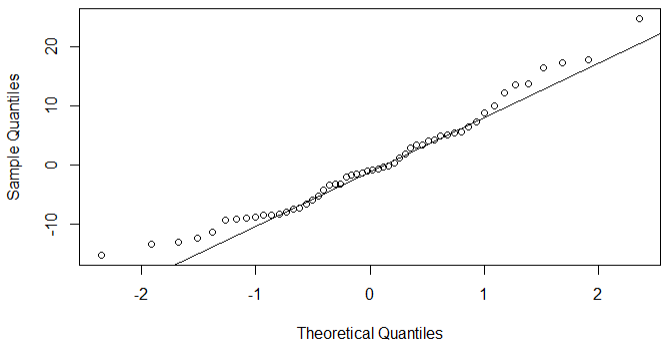

A Shapiro-Wilk test indicated that model residuals were normally distributed (*W* = .97; *p* = .13). Variance inflation factors for model predictors did not indicate multicollinearity (${IP}_{1}$: 2.62; ${IP}_{2}$: 1.78; ${\Delta IP}_{2}$: 3.01; $pS$: 1.19; Baseline State Anxiety: 1.25; Age: 1.19; Sex: 1.09).

##### 6.2.2 Trait anxiety change

*Linear Mixed Effects Model to Predict Change in Trait Anxiety*

| *Predictors* | *Estimates* | *Std. Error* | *Statistic* | *p* | *df* |
| --- | --- | --- | --- | --- | --- |
| (Intercept) | -23.02 | 16.23 | -1.42 | 0.163 | 46.01 |
| ${IP}_{1}$ | 28.29 | 18.29 | 1.55 | 0.129 | 46.00 |
| ${IP}_{2}$ | 0.45 | 1.88 | 0.24 | 0.811 | 46.00 |
| ${\Delta IP}_{2}$ | -3.72 | 1.82 | -2.04 | **0.047** | 46.00 |
| $pS$ | 1.68 | 2.69 | 0.63 | 0.535 | 46.00 |
| Baseline Trait Anxiety | -0.27 | 0.08 | -3.19 | **0.003** | 46.00 |
| Age | 0.15 | 0.12 | 1.29 | 0.205 | 46.00 |
| Sex | 0.44 | 1.12 | 0.39 | 0.698 | 48.23 |
| **Random Effects** | | | | | |
| σ^2^ | 41.68 | | | | |
| τ_00_ _Sex_ | 0.06 | | | | |
| ICC | 0.00 | | | | |
| N _Sex_ | 2 | | | | |
| Observations | 54 | | | | |
| Marginal R^2^ / Conditional R^2^ | 0.283 / 0.284 | | | | |

**Legend.** Kenward-Roger approximation was used for degrees of freedom and *p*-values. The marginal R-squared accounts for the variance of the fixed effects only, while the conditional R-squared accounts for both the fixed and random effects. σ^2^: Residual variance; τ_00_: variance for this random intercepts factor; ICC: intra-class correlation for this random intercepts factor; N: number of levels for this random intercepts factor.

*Quantile-Quantile Plot of Model Residuals*

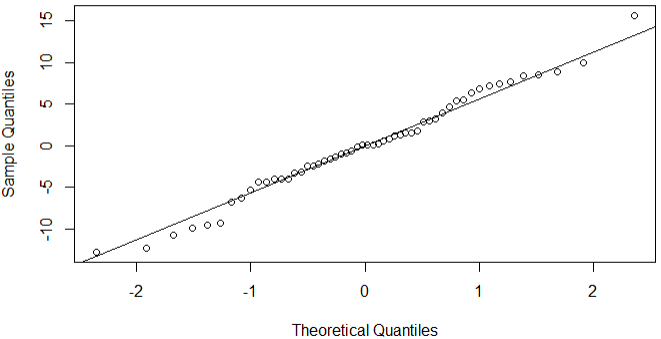

A Shapiro-Wilk test indicated that model residuals were normally distributed (*W* = .99; *p* = .80). Variance inflation factors for model predictors did not indicate multicollinearity (${IP}_{1}$: 2.65; ${IP}_{2}$: 1.80; ${\Delta IP}_{2}$: 3.10; $pS$: 1.17; Baseline Trait Anxiety: 1.08; Age: 1.09; Sex: 1.09).

#### 6.3 In control group only

##### 6.3.1 State anxiety change

Multiple regression was used for this analysis as no random effects structure for a linear mixed effects model could be found that produced a non-singular fit.

*Multiple Regression Model to Predict Change in State Anxiety*

| *Predictors* | *Estimates* | *std. Error* | *Statistic* | *p* | *df* |
| --- | --- | --- | --- | --- | --- |
| (Intercept) | -53.60 | 50.47 | -1.06 | 0.302 | 18.00 |
| ${IP}_{1}$ | 109.07 | 72.34 | 1.51 | 0.149 | 18.00 |
| ${IP}_{2}$ | -9.34 | 14.08 | -0.66 | 0.515 | 18.00 |
| ${\Delta IP}_{2}$ | -4.44 | 4.64 | -0.96 | 0.352 | 18.00 |
| $pS$ | -1.37 | 5.21 | -0.26 | 0.795 | 18.00 |
| Baseline State Anxiety | -0.64 | 0.23 | -2.73 | **0.014** | 18.00 |
| Age | -0.28 | 0.27 | -1.03 | 0.318 | 18.00 |
| Sex | 0.26 | 2.30 | 0.11 | 0.910 | 18.00 |
| Observations | 26 | | | | |
| R^2^ / R^2^ adjusted | 0.487 / 0.288 | | | | |

*Quantile-Quantile Plot of Model Residuals*

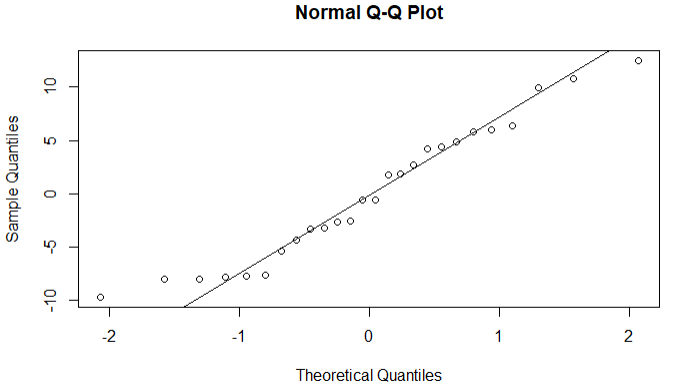

A Shapiro-Wilk test indicated that model residuals were normally distributed (*W* = .95; *p* = .24). Variance inflation factors for model predictors strongly indicated multicollinearity (${IP}_{1}$: 12.94; ${IP}_{2}$: 3.03; ${\Delta IP}_{2}$: 9.52; $pS$: 1.27; Baseline State Anxiety: 1.33; Age: 1.35; Sex: 1.24).

##### 6.3.2 Trait anxiety change

Multiple regression was used for this analysis as no random effects structure for a linear mixed effects model could be found that produced a non-singular fit.

*Multiple Regression Model to Predict Change in Trait Anxiety*

| *Predictors* | *Estimates* | *std. Error* | *Statistic* | *p* | *df* |
| --- | --- | --- | --- | --- | --- |
| (Intercept) | -33.06 | 35.94 | -0.92 | 0.370 | 18.00 |
| ${IP}_{1}$ | 83.85 | 50.58 | 1.66 | 0.115 | 18.00 |
| ${IP}_{2}$ | -15.88 | 9.77 | -1.63 | 0.121 | 18.00 |
| ${\Delta IP}_{2}$ | -4.56 | 3.26 | -1.40 | 0.179 | 18.00 |
| $pS$ | -3.09 | 3.62 | -0.86 | 0.404 | 18.00 |
| Baseline Trait Anxiety | -0.41 | 0.12 | -3.50 | **0.003** | 18.00 |
| Age | 0.26 | 0.18 | 1.47 | 0.159 | 18.00 |
| Sex | 1.32 | 1.67 | 0.79 | 0.439 | 18.00 |
| Observations | 26 | | | | |
| R^2^ / R^2^ adjusted | 0.475 / 0.271 | | | | |

*Quantile-Quantile Plot of Model Residuals*

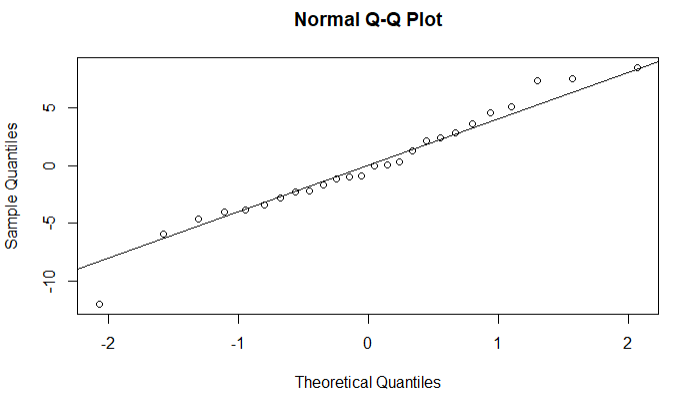

A Shapiro-Wilk test indicated that model residuals were normally distributed (*W* = .97; *p* = .68). Variance inflation factors for model predictors strongly indicated multicollinearity (${IP}_{1}$: 12.44; ${IP}_{2}$: 2.87; ${\Delta IP}_{2}$: 9.22; $pS$: 1.21; Baseline Trait Anxiety: 1.12; Age: 1.17; Sex: 1.29).

### 7 Relationships between change in anxiety and conventional interoceptive task measures

#### 7.1 In training group only

##### 7.1.1 State anxiety change

Multiple regression was used for this analysis, as no mixed effects model produced a non-singular fit.

*Multiple Regression Model to Predict Change in State Anxiety*

| *Predictors* | *Estimates* | *std. Error* | *Statistic* | *p* | *df* |
| --- | --- | --- | --- | --- | --- |
| (Intercept) | 22.82 | 8.98 | 2.54 | **0.019** | 21.00 |
| *d’* change | 0.70 | 2.64 | 0.27 | 0.793 | 21.00 |
| *C* change | -0.10 | 9.10 | -0.01 | 0.991 | 21.00 |
| HBT change | -12.81 | 10.21 | -1.25 | 0.223 | 21.00 |
| Baseline State Anxiety | -0.57 | 0.20 | -2.80 | **0.011** | 21.00 |
| Age | 0.19 | 0.29 | 0.64 | 0.526 | 21.00 |
| Sex | -0.02 | 2.57 | -0.01 | 0.995 | 21.00 |
| Observations | 28 | | | | |
| R^2^ / R^2^adjusted | 0.394 / 0.221 | | | | |

**Legend.** *d’*: signal detection sensitivity from heartbeat discrimination task; *C*: signal detection criterion from heartbeat discrimination task; HBT: performance accuracy from heartbeat counting task (calculation detailed in main text).

*Quantile-Quantile Plot of Model Residuals*

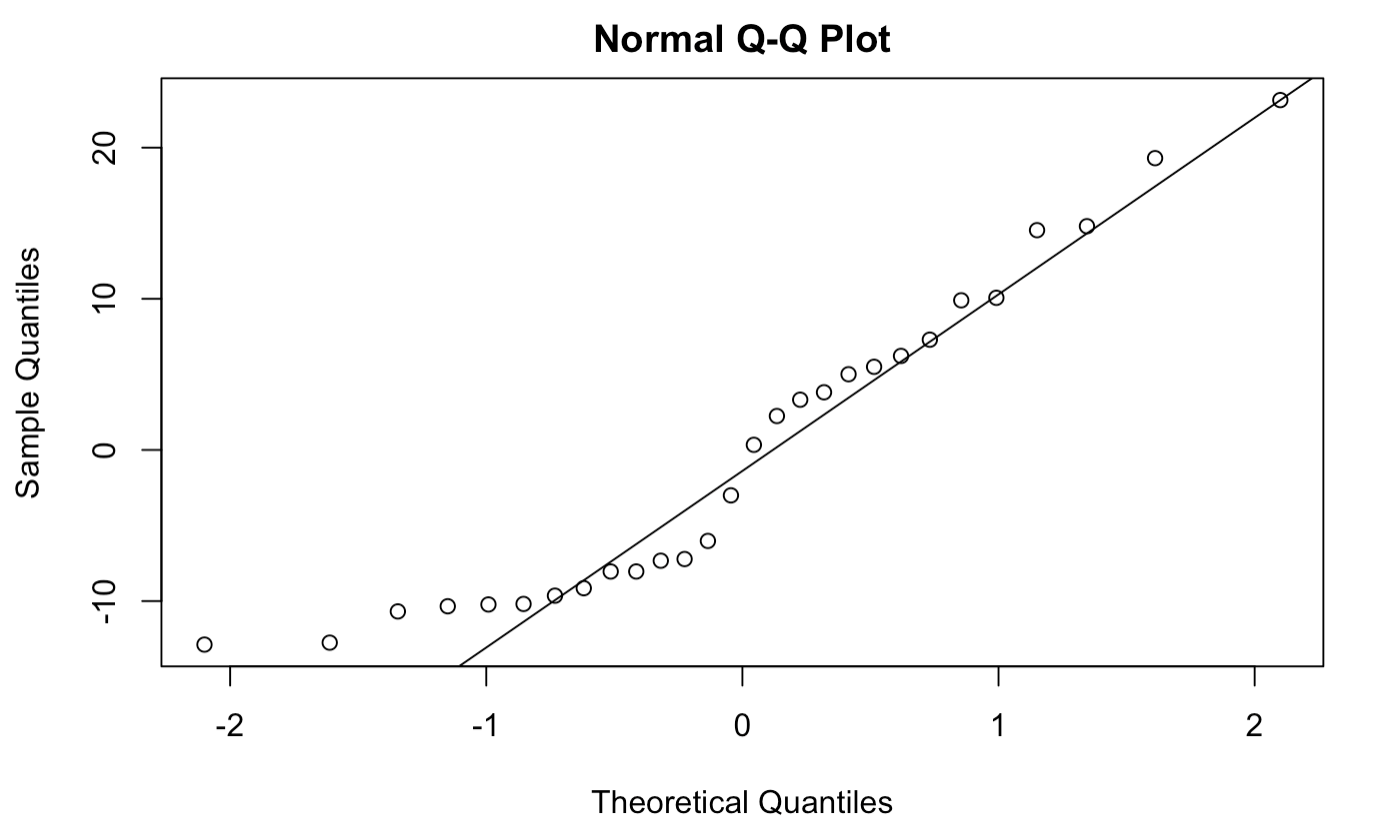

A Shapiro-Wilk test indicated that model residuals were not normally distributed (*W* = .91; *p* = .03). Variance inflation factors for model predictors did not indicate multicollinearity (*d’* change: 1.52; *C* change: 1.18; HBT change: 1.16; Baseline State Anxiety: 1.60; Age: 1.30; Sex: 1.08).

##### 7.1.2 Trait anxiety change

*Linear Mixed Effects Model to Predict Change in Trait Anxiety*

| *Predictors* | *Estimates* | *std. Error* | *Statistic* | *p* | | *df* | |
| --- | --- | --- | --- | --- | --- | --- | --- |
| (Intercept) | 2.91 | 6.89 | 0.42 | | 0.677 | | 21.07 |
| *d’* change | 1.40 | 1.56 | 0.90 | | 0.381 | | 21.00 |
| *C* change | 1.55 | 5.69 | 0.27 | | 0.789 | | 21.00 |
| HBT change | -7.12 | 6.70 | -1.06 | | 0.300 | | 21.00 |
| Baseline Trait Anxiety | -0.19 | 0.13 | -1.47 | | 0.155 | | 21.00 |
| Age | 0.10 | 0.17 | 0.58 | | 0.569 | | 21.00 |
| Sex | 0.65 | 1.69 | 0.38 | | 0.705 | | 22.28 |
| **Random Effects** | | | | | | | |
| σ^2^ | 60.09 | | | | | | |
| τ_00_ _Sex_ | 0.17 | | | | | | |
| ICC | 0.00 | | | | | | |
| N _Sex_ | 2 | | | | | | |
| Observations | 28 | | | | | | |
| Marginal R^2^/ Conditional R^2^ | 0.122 / 0.125 | | | | | | |

**Legend.** Kenward-Roger approximation was used for degrees of freedom and *p*-values. The marginal R-squared accounts for the variance of the fixed effects only, while the conditional R-squared accounts for both the fixed and random effects.

*d’*: signal detection sensitivity from heartbeat discrimination task; *C*: signal detection criterion from heartbeat discrimination task; HBT: performance accuracy from heartbeat counting task (calculation detailed in main text); σ^2^: Residual variance; τ_00_: variance for this random intercepts factor; ICC: intra-class correlation for this random intercepts factor; N: number of levels for this random intercepts factor.

*Quantile-Quantile Plot of Model Residuals*

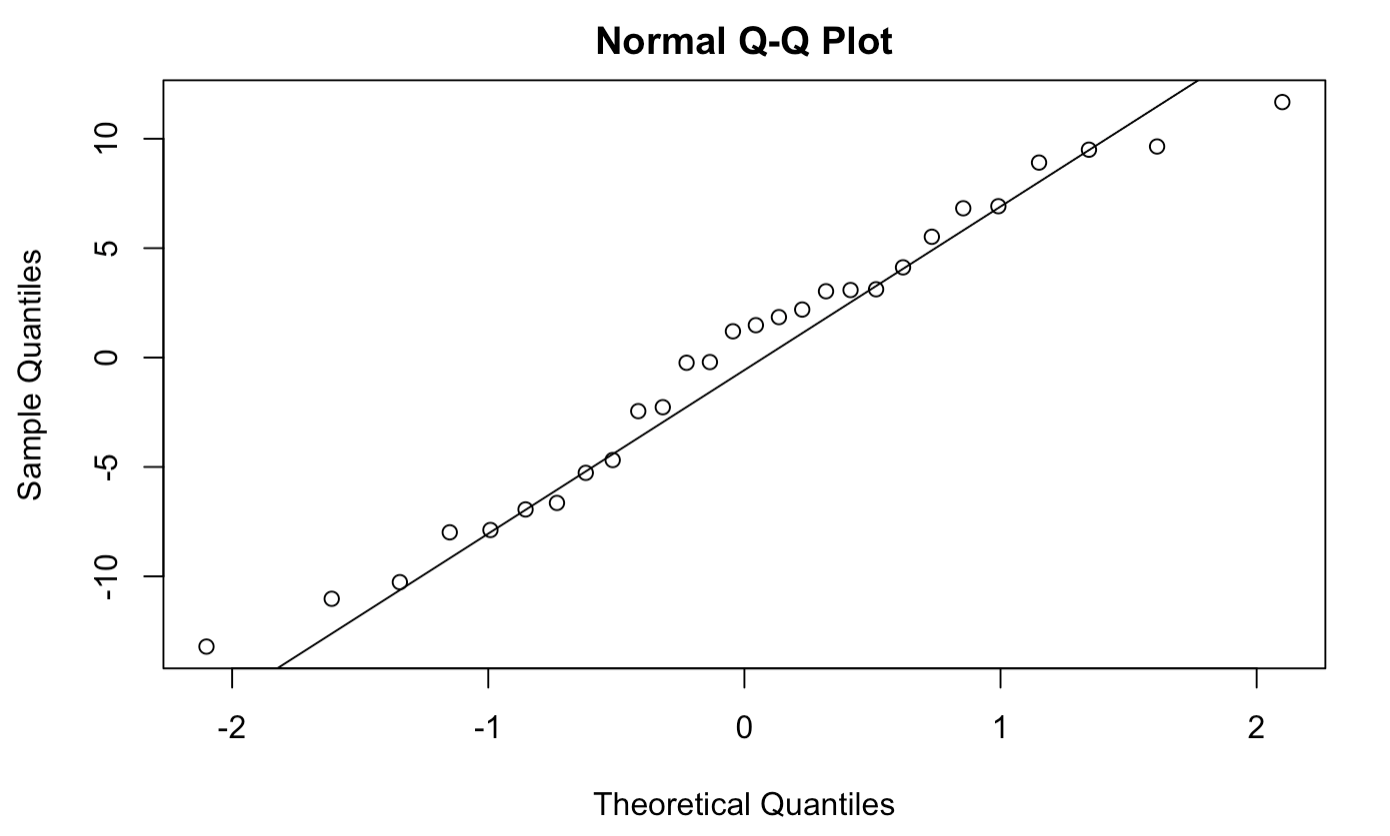

A Shapiro-Wilk test indicated that model residuals were normally distributed (*W* = .97; *p* = .55). Variance inflation factors for model predictors did not indicate multicollinearity (*d’* change: 1.24; *C* change: 1.07; HBT change: 1.15; Baseline Trait Anxiety: 1.05; Age: 1.06; Sex: 1.06).

#### 7.2 In control group only

##### 7.2.1 State anxiety change

Multiple regression was used for this analysis, as no mixed effects model produced a non-singular fit.

*Multiple Regression Model to Predict Change in State Anxiety*

| *Predictors* | *Estimates* | *std. Error* | *Statistic* | *p* | *df* |
| --- | --- | --- | --- | --- | --- |
| (Intercept) | 18.15 | 8.63 | 2.10 | **0.050** | 18.00 |
| *d’* change | -0.94 | 1.86 | -0.50 | 0.620 | 18.00 |
| *C* change | 8.29 | 6.54 | 1.27 | 0.221 | 18.00 |
| HBT change | 7.04 | 8.95 | 0.79 | 0.442 | 18.00 |
| Baseline State Anxiety | -0.51 | 0.24 | -2.18 | **0.043** | 18.00 |
| Age | -0.37 | 0.27 | -1.38 | 0.184 | 18.00 |
| Sex | 0.53 | 2.36 | 0.22 | 0.826 | 18.00 |
| Observations | 25 | | | | |
| R^2^ / R^2^ adjusted | 0.406 / 0.208 | | | | |

**Legend.** *d’*: signal detection sensitivity from heartbeat discrimination task; *C*: signal detection criterion from heartbeat discrimination task; HBT: performance accuracy from heartbeat counting task (calculation detailed in main text).

*Quantile-Quantile Plot of Model Residuals*

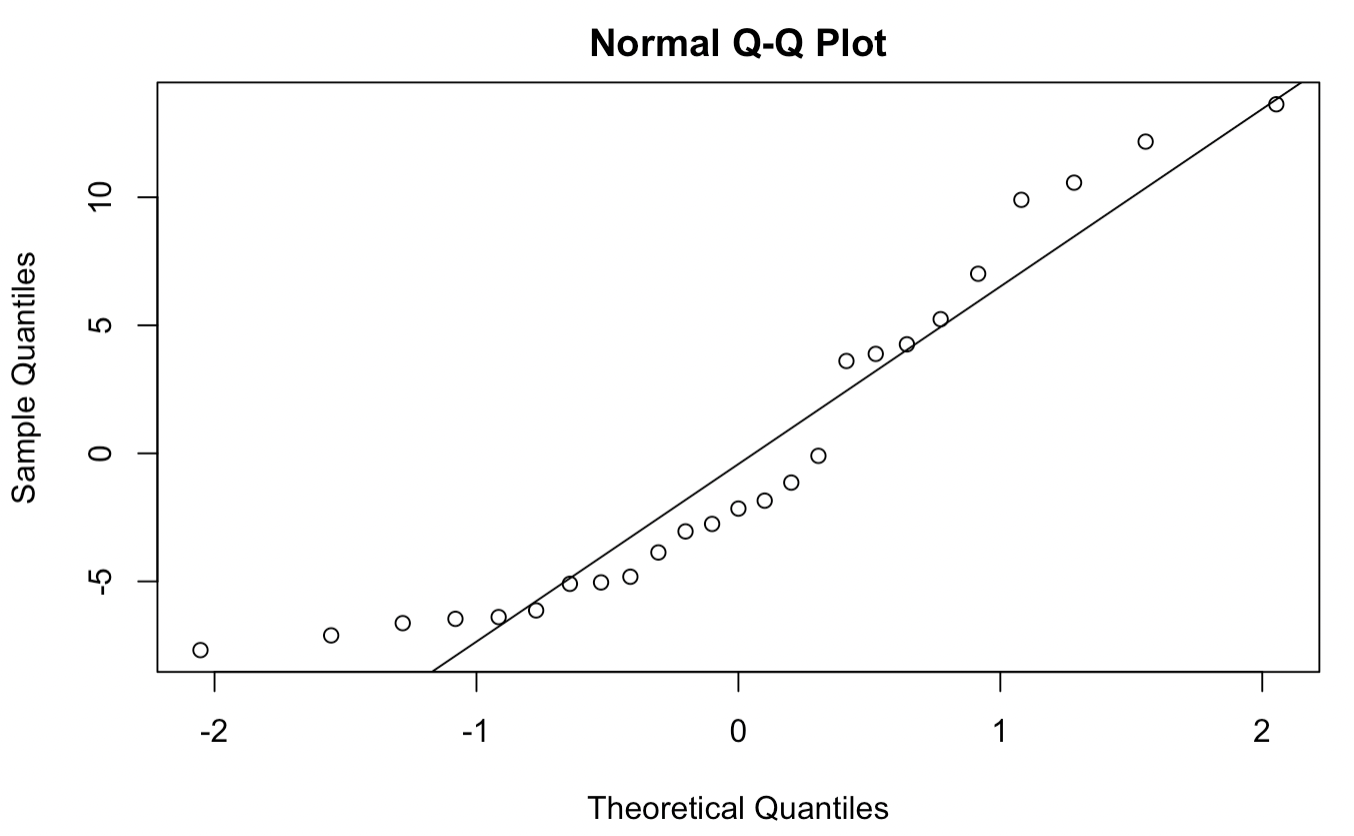

A Shapiro-Wilk test indicated that model residuals were not normally distributed (*W* = .89; *p* = .01). Variance inflation factors for model predictors did not indicate multicollinearity (*d’* change: 1.25; *C* change: 1.13; HBT change: 1.36; Baseline State Anxiety: 1.27; Age: 1.27; Sex: 1.27).

##### 7.2.2 Trait anxiety change

*Linear Mixed Effects Model to Predict Change in Trait Anxiety*

| *Predictors* | *Estimates* | *std. Error* | *Statistic* | *p* | | *df* | |
| --- | --- | --- | --- | --- | --- | --- | --- |
| (Intercept) | 14.54 | 5.58 | 2.60 | | **0.018** | | 18.19 |
| *d’* change | 0.71 | 1.42 | 0.50 | | 0.624 | | 18.00 |
| *C* change | 1.08 | 4.86 | 0.22 | | 0.826 | | 18.00 |
| HBT change | -3.75 | 6.54 | -0.57 | | 0.573 | | 18.00 |
| Baseline Trait Anxiety | -0.38 | 0.12 | -3.07 | | **0.007** | | 18.00 |
| Age | 0.27 | 0.19 | 1.47 | | 0.160 | | 18.00 |
| Sex | 0.80 | 1.88 | 0.43 | | 0.673 | | 19.79 |
| **Random Effects** | | | | | | | |
| σ^2^ | 33.34 | | | | | | |
| τ_00_ _Sex_ | 0.33 | | | | | | |
| ICC | 0.01 | | | | | | |
| N _Sex_ | 2 | | | | | | |
| Observations | 25 | | | | | | |
| Marginal R^2^/ Conditional R^2^ | 0.334 / 0.341 | | | | | | |

**Legend.** Kenward-Roger approximation was used for degrees of freedom and *p*-values. The marginal R-squared accounts for the variance of the fixed effects only, while the conditional R-squared accounts for both the fixed and random effects.

*d’*: signal detection sensitivity from heartbeat discrimination task; *C*: signal detection criterion from heartbeat discrimination task; HBT: performance accuracy from heartbeat counting task (calculation detailed in main text); σ^2^: Residual variance; τ_00_: variance for this random intercepts factor; ICC: intra-class correlation for this random intercepts factor; N: number of levels for this random intercepts factor.

*Quantile-Quantile Plot of Model Residuals*

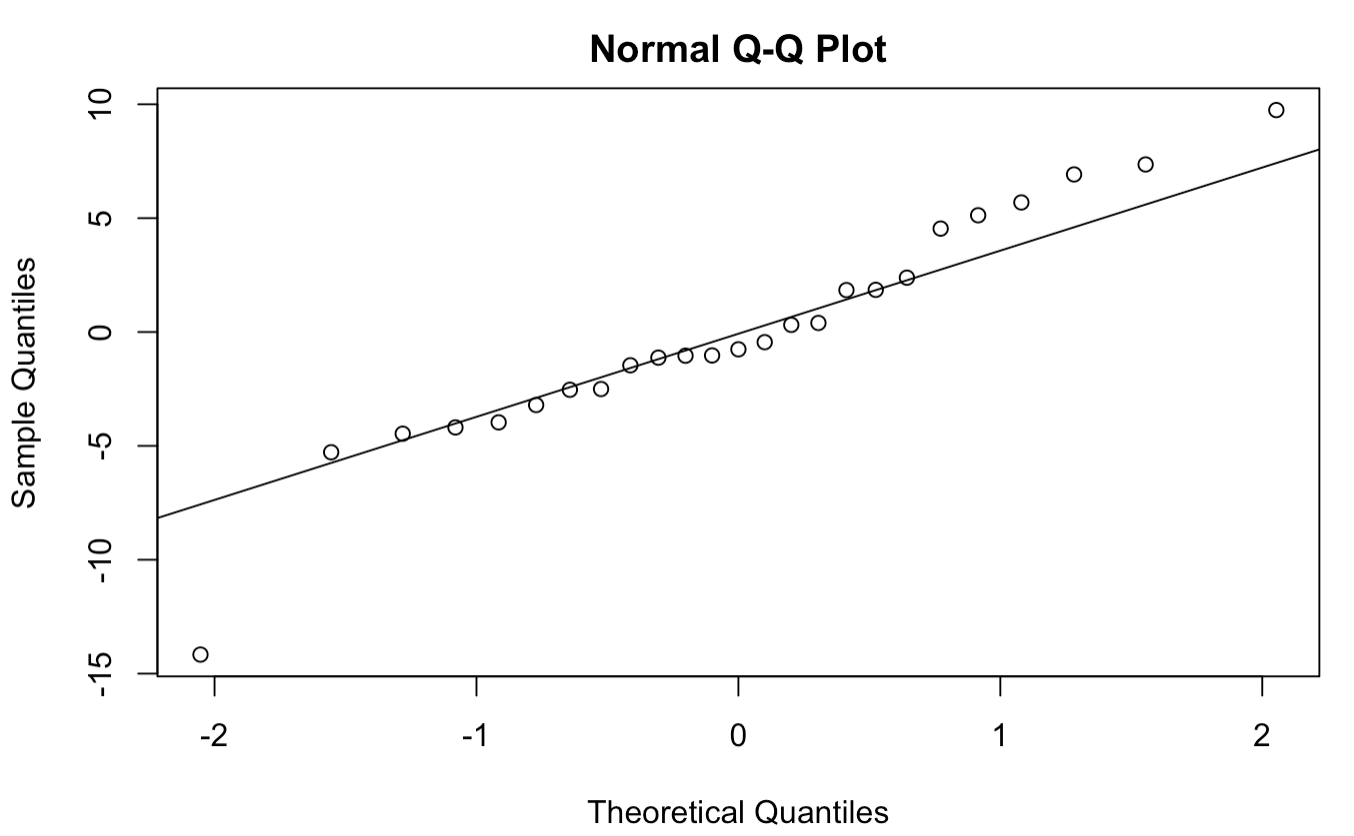

A Shapiro-Wilk test indicated that model residuals were normally distributed (*W* = .95; *p* = .26). Variance inflation factors for model predictors did not indicate multicollinearity (*d’* change: 1.27; *C* change: 1.10; HBT change: 1.27; Baseline Trait Anxiety: 1.09; Age: 1.10; Sex: 1.33).

#### 7.3 In both groups

##### 7.3.1 State anxiety change

Multiple regression was used for this analysis, as no mixed effects model produced a non-singular fit.

*Multiple Regression Model to Predict Change in State Anxiety*

| *Predictors* | *Estimates* | *std. Error* | *Statistic* | *p* | *df* |
| --- | --- | --- | --- | --- | --- |
| (Intercept) | 19.80 | 5.93 | 3.34 | **0.002** | 46.00 |
| *d’* Change | -0.64 | 1.45 | -0.44 | 0.663 | 46.00 |
| *C* Change | 3.64 | 5.46 | 0.67 | 0.509 | 46.00 |
| HBT Change | -7.99 | 6.37 | -1.25 | 0.216 | 46.00 |
| Baseline State Anxiety | -0.51 | 0.14 | -3.72 | **0.001** | 46.00 |
| Age | -0.05 | 0.19 | -0.26 | 0.796 | 46.00 |
| Sex | 1.11 | 1.64 | 0.67 | 0.504 | 46.00 |
| Observations | 53 | | | | |
| R^2^ / R^2^adjusted | 0.386 / 0.306 | | | | |

**Legend.** *d’*: signal detection sensitivity from heartbeat discrimination task; *C*: signal detection criterion from heartbeat discrimination task; HBT: performance accuracy from heartbeat counting task (calculation detailed in main text).

*Quantile-Quantile Plot of Model Residuals*

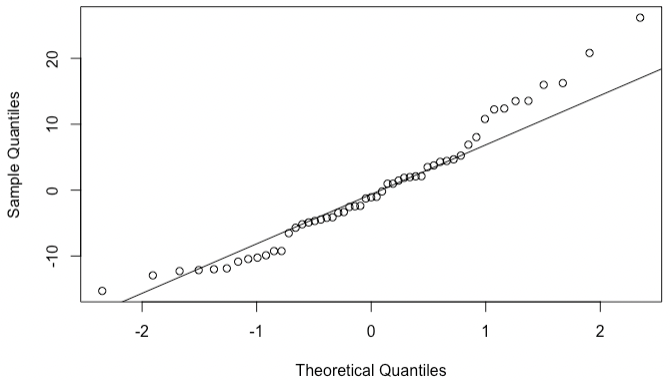

A Shapiro-Wilk test indicated that model residuals were normally distributed (*W* = .96; *p* = .07). Variance inflation factors for model predictors did not indicate multicollinearity (*d’* change: 1.53; *C* change: 1.06; HBT change: 1.39; Baseline State Anxiety: 1.36; Age: 1.20; Sex: 1.03).

##### 7.3.2 Trait anxiety change

*Linear Mixed Effects Model to Predict Change in Trait Anxiety*

| *Predictors* | *Estimates* | *std. Error* | *Statistic* | *p* | *df* |
| --- | --- | --- | --- | --- | --- |
| (Intercept) | 8.66 | 4.62 | 1.87 | 0.078 | 17.28 |
| *d’* change | 1.07 | 1.06 | 1.01 | 0.317 | 44.93 |
| *C* change | 0.50 | 3.59 | 0.14 | 0.889 | 45.00 |
| HBT change | -6.76 | 4.63 | -1.46 | 0.151 | 45.81 |
| Baseline Trait Anxiety | -0.27 | 0.09 | -3.12 | **0.003** | 45.32 |
| Age | 0.15 | 0.12 | 1.24 | 0.221 | 45.72 |
| Sex | 0.51 | 1.12 | 0.45 | 0.654 | 45.70 |
| **Random Effects** | | | | | |
| σ^2^ | 42.99 | | | | |
| τ_00_ _Group_ | 5.54 | | | | |
| ICC | 0.11 | | | | |
| N _Group_ | 2 | | | | |
| Observations | 53 | | | | |
| Marginal R^2^/ Conditional R^2^ | 0.191 / 0.284 | | | | |

**Legend.** Kenward-Roger approximation was used for degrees of freedom and *p*-values. The marginal R-squared accounts for the variance of the fixed effects only, while the conditional R-squared accounts for both the fixed and random effects. *d’*: signal detection sensitivity from heartbeat discrimination task; *C*: signal detection criterion from heartbeat discrimination task; HBT: performance accuracy from heartbeat counting task (calculation detailed in main text); σ^2^: Residual variance; τ_00_: variance for this random intercepts factor; ICC: intra-class correlation for this random intercepts factor; N: number of levels for this random intercepts factor.

*Quantile-Quantile Plot of Model Residuals*

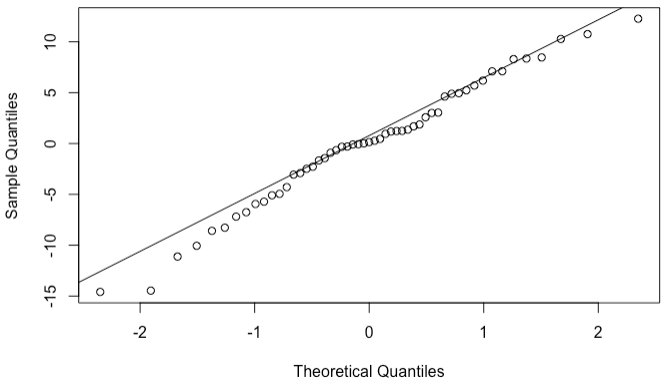

A Shapiro-Wilk test indicated that model residuals were normally distributed (*W* = .98; *p* = .61). Variance inflation factors for model predictors did not indicate multicollinearity (*d’* change: 1.23; *C* change: 1.04; HBT change: 1.21; Baseline Trait Anxiety: 1.03; Age: 1.03; Sex: 1.03).

### 8 Change in self-reported interoception across assessments (Multidimensional Assessment of Interoceptive Awareness sub-scales)

#### 8.1 Total score

*Linear Mixed Effects Model to Predict MAIA Total Score*

| *Predictors* | *Estimates* | *std. Error* | *Statistic* | *p* | *df* |
| --- | --- | --- | --- | --- | --- |
| (Intercept) | 21.76 | 0.78 | 27.96 | **<0.001** | 58.02 |
| Group | -0.83 | 0.67 | -1.25 | 0.217 | 61.15 |
| Time [Final] | 1.13 | 0.43 | 2.65 | **0.011** | 52.00 |
| Age | 0.16 | 0.08 | 2.01 | **0.050** | 50.00 |
| Sex | 0.90 | 0.75 | 1.19 | 0.238 | 50.00 |
| Group × Time [Final] | 1.28 | 0.43 | 3.01 | **0****.004** | 52.00 |
| **Random Effects** | | | | | |
| σ^2^ | 4.87 | | | | |
| τ_00_ _Participant_ | 18.19 | | | | |
| ICC | 0.79 | | | | |
| N _Participant_ | 54 | | | | |
| Observations | 108 | | | | |
| Marginal R^2^ / Conditional R^2^ | 0.109 / 0.812 | | | | |

**Legend.** Kenward-Roger approximation was used for degrees of freedom and *p*-values. The marginal R-squared accounts for the variance of the fixed effects only, while the conditional R-squared accounts for both the fixed and random effects. σ^2^: Residual variance; τ_00_: variance for this random intercepts factor; ICC: intra-class correlation for this random intercepts factor; N: number of levels for this random intercepts factor.

*Quantile-Quantile Plot of Model Residuals*

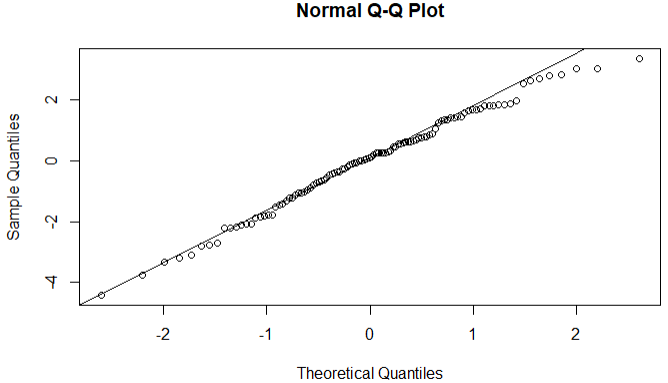

A Shapiro-Wilk test indicated that model residuals were normally distributed (*W* = .99; *p* = .48).

#### 8.2 Subscale scores

##### 8.2.1 Noticing

*Linear Mixed Effects Model to Predict MAIA Noticing Score*

| *Predictors* | *Estimates* | *std. Error* | *Statistic* | *p* | *df* |
| --- | --- | --- | --- | --- | --- |
| (Intercept) | 2.94 | 0.15 | 19.21 | **<0.001** | 65.71 |
| Group | 0.06 | 0.13 | 0.48 | 0.631 | 71.49 |
| Time [Final] | 0.21 | 0.11 | 1.83 | 0.073 | 52.00 |
| Age | 0.03 | 0.02 | 1.77 | 0.083 | 50.00 |
| Sex | 0.30 | 0.14 | 2.12 | **0.039** | 50.00 |
| Group × Time [Final] | 0.26 | 0.11 | 2.25 | **0.028** | 52.00 |
| **Random Effects** | | | | | |
| σ^2^ | 0.35 | | | | |
| τ_00_ _Participant_ | 0.57 | | | | |
| ICC | 0.62 | | | | |
| N _Participant_ | 54 | | | | |
| Observations | 108 | | | | |
| Marginal R^2^/ Conditional R^2^ | 0.153 / 0.677 | | | | |

**Legend.** Kenward-Roger approximation was used for degrees of freedom and *p*-values. The marginal R-squared accounts for the variance of the fixed effects only, while the conditional R-squared accounts for both the fixed and random effects. σ^2^: Residual variance; τ_00_: variance for this random intercepts factor; ICC: intra-class correlation for this random intercepts factor; N: number of levels for this random intercepts factor.

*Quantile-Quantile Plot of Model Residuals*

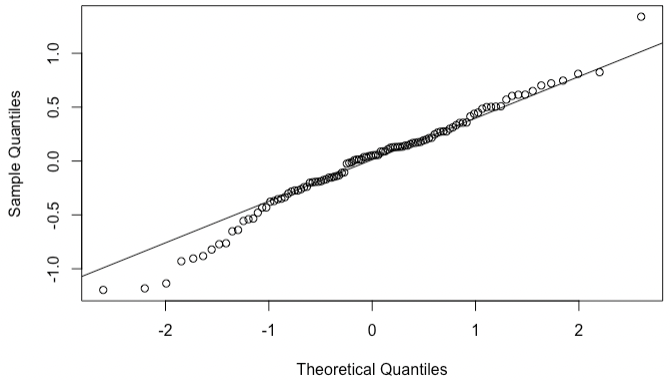

A Shapiro-Wilk test indicated that model residuals were normally distributed (*W* = .98; *p* = .13).

##### 8.2.2 Not-Distracting

*Linear Mixed Effects Model to Predict MAIA Not-Distracting Score*

| *Predictors* | *Estimates* | *std. Error* | *Statistic* | *p* | *df* |
| --- | --- | --- | --- | --- | --- |
| (Intercept) | 2.59 | 0.15 | 16.85 | **<0.001** | 71.24 |
| Group | -0.62 | 0.13 | -4.60 | **<0.001** | 78.52 |
| Time [final] | 0.10 | 0.13 | 0.77 | 0.442 | 52.00 |
| Age | 0.00 | 0.01 | 0.21 | 0.831 | 50.00 |
| Sex | -0.10 | 0.14 | -0.72 | 0.473 | 50.00 |
| Group × Time [final] | 0.13 | 0.13 | 0.97 | 0.334 | 52.00 |
| **Random Effects** | | | | | |
| σ^2^ | 0.46 | | | | |
| τ_00_ _Participant_ | 0.48 | | | | |
| ICC | 0.51 | | | | |
| N _Participant_ | 54 | | | | |
| Observations | 108 | | | | |
| Marginal R^2^/ Conditional R^2^ | 0.245 / 0.630 | | | | |

**Legend.** Kenward-Roger approximation was used for degrees of freedom and *p*-values. The marginal R-squared accounts for the variance of the fixed effects only, while the conditional R-squared accounts for both the fixed and random effects. σ^2^: Residual variance; τ_00_: variance for this random intercepts factor; ICC: intra-class correlation for this random intercepts factor; N: number of levels for this random intercepts factor.

*Quantile-Quantile Plot of Model Residuals*

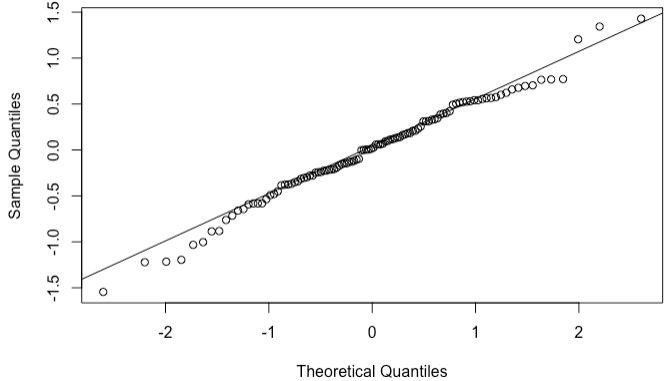

A Shapiro-Wilk test indicated that model residuals were normally distributed (*W* = .99; *p* = .40).

##### 8.2.3 Not-Worrying

*Linear Mixed Effects Model to Predict MAIA Not-Worrying Score*

| *Predictors* | *Estimates* | *std. Error* | *Statistic* | *p* | *df* |
| --- | --- | --- | --- | --- | --- |
| (Intercept) | 2.88 | 0.19 | 15.10 | **<0.001** | 65.73 |
| Group | 0.03 | 0.17 | 0.16 | 0.876 | 71.51 |
| Time [Final] | -0.00 | 0.14 | -0.02 | 0.988 | 52.00 |
| Age | -0.00 | 0.02 | -0.02 | 0.986 | 50.00 |
| Sex | -0.02 | 0.18 | -0.14 | 0.892 | 50.00 |
| Group × Time [Final] | 0.05 | 0.14 | 0.35 | 0.727 | 52.00 |
| **Random Effects** | | | | | |
| σ^2^ | 0.54 | | | | |
| τ_00_ _Participant_ | 0.88 | | | | |
| ICC | 0.62 | | | | |
| N _Participant_ | 54 | | | | |
| Observations | 108 | | | | |
| Marginal R^2^/ Conditional R^2^ | 0.003 / 0.620 | | | | |

**Legend.** Kenward-Roger approximation was used for degrees of freedom and *p*-values. The marginal R-squared accounts for the variance of the fixed effects only, while the conditional R-squared accounts for both the fixed and random effects. σ^2^: Residual variance; τ_00_: variance for this random intercepts factor; ICC: intra-class correlation for this random intercepts factor; N: number of levels for this random intercepts factor.

*Quantile-Quantile Plot of Model Residuals*

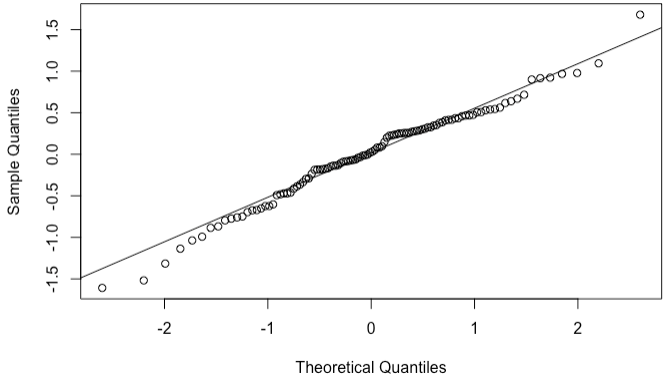

A Shapiro-Wilk test indicated that model residuals were normally distributed (*W* = .98; *p* = .18).

##### 8.2.4 Attention Regulation

*Linear Mixed Effects Model to Predict MAIA Attention Regulation Score*

| *Predictors* | *Estimates* | *std. Error* | *Statistic* | *p* | *df* |
| --- | --- | --- | --- | --- | --- |
| (Intercept) | 2.61 | 0.16 | 15.86 | **<0.001** | 61.23 |
| Group | 0.08 | 0.14 | 0.59 | 0.559 | 65.53 |
| Time [Final] | 0.20 | 0.11 | 1.89 | 0.064 | 52.00 |
| Age | 0.02 | 0.02 | 1.09 | 0.281 | 50.00 |
| Sex | 0.13 | 0.16 | 0.84 | 0.404 | 50.00 |
| Group × Time [Final] | 0.19 | 0.11 | 1.80 | 0.078 | 52.00 |
| **Random Effects** | | | | | |
| σ^2^ | 0.30 | | | | |
| τ_00_ _Participant_ | 0.75 | | | | |
| ICC | 0.71 | | | | |
| N _Participant_ | 54 | | | | |
| Observations | 108 | | | | |
| Marginal R^2^/ Conditional R^2^ | 0.074 / 0.736 | | | | |

**Legend.** Kenward-Roger approximation was used for degrees of freedom and *p*-values. The marginal R-squared accounts for the variance of the fixed effects only, while the conditional R-squared accounts for both the fixed and random effects. σ^2^: Residual variance; τ_00_: variance for this random intercepts factor; ICC: intra-class correlation for this random intercepts factor; N: number of levels for this random intercepts factor.

*Quantile-Quantile Plot of Model Residuals*

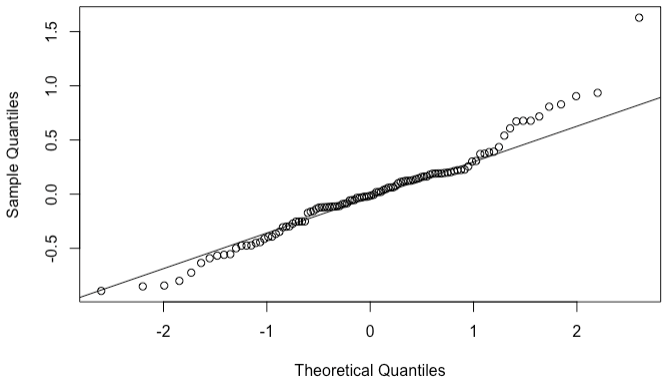

A Shapiro-Wilk test indicated that model residuals were not normally distributed (*W* = .96; *p* = .006).

##### 8.2.5 Emotional Awareness

*Linear Mixed Effects Model to Predict MAIA Emotional Awareness Score*

| *Predictors* | *Estimates* | *std. Error* | *Statistic* | *p* | | *df* | |
| --- | --- | --- | --- | --- | --- | --- | --- |
| (Intercept) | 3.04 | 0.16 | 18.70 | | **<0.001** | | 59.94 |
| Group | -0.04 | 0.14 | -0.26 | | 0.795 | | 63.78 |
| Time [Final] | 0.11 | 0.10 | 1.07 | | 0.290 | | 52.00 |
| Age | 0.04 | 0.02 | 2.71 | | **0.009** | | 50.00 |
| Sex | 0.26 | 0.16 | 1.67 | | 0.101 | | 50.00 |
| Group × Time [Final] | 0.24 | 0.10 | 2.44 | | **0.018** | | 52.00 |
| **Random Effects** | | | | | | | |
| σ^2^ | 0.26 | | | | | | |
| τ_00_ _Participant_ | 0.76 | | | | | | |
| ICC | 0.74 | | | | | | |
| N _Participant_ | 54 | | | | | | |
| Observations | 108 | | | | | | |
| Marginal R^2^/ Conditional R^2^ | 0.159 / 0.785 | | | | | | |

**Legend.** Kenward-Roger approximation was used for degrees of freedom and *p*-values. The marginal R-squared accounts for the variance of the fixed effects only, while the conditional R-squared accounts for both the fixed and random effects. σ^2^: Residual variance; τ_00_: variance for this random intercepts factor; ICC: intra-class correlation for this random intercepts factor; N: number of levels for this random intercepts factor.

*Quantile-Quantile Plot of Model Residuals*

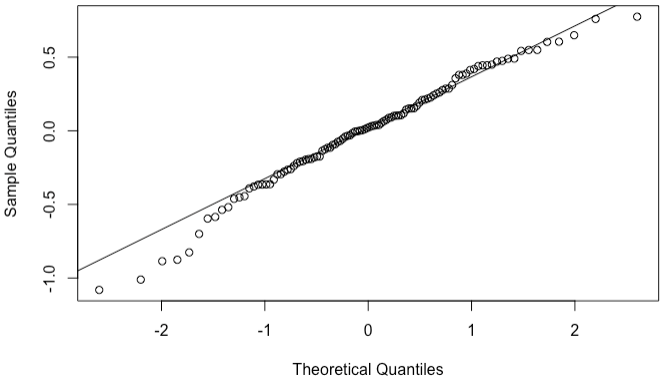

A Shapiro-Wilk test indicated that model residuals were normally distributed (*W* = .98; *p* = .15).

##### 8.2.6 Self-Regulation

*Linear Mixed Effects Model to Predict MAIA Self-Regulation Score*

| *Predictors* | *Estimates* | *std. Error* | *Statistic* | *p* | *df* |
| --- | --- | --- | --- | --- | --- |
| (Intercept) | 2.68 | 0.18 | 15.16 | **<0.001** | 62.60 |
| Group | -0.12 | 0.15 | -0.80 | 0.426 | 67.37 |
| Time [Final] | 0.16 | 0.12 | 1.37 | 0.177 | 52.00 |
| Age | 0.02 | 0.02 | 1.32 | 0.194 | 50.00 |
| Sex | 0.07 | 0.17 | 0.42 | 0.676 | 50.00 |
| Group × Time [Final] | 0.29 | 0.12 | 2.47 | **0.017** | 52.00 |
| **Random Effects** | | | | | |
| σ^2^ | 0.38 | | | | |
| τ_00_ _Participant_ | 0.83 | | | | |
| ICC | 0.68 | | | | |
| N _Participant_ | 54 | | | | |
| Observations | 108 | | | | |
| Marginal R^2^/ Conditional R^2^ | 0.053 / 0.701 | | | | |

**Legend.** Kenward-Roger approximation was used for degrees of freedom and *p*-values. The marginal R-squared accounts for the variance of the fixed effects only, while the conditional R-squared accounts for both the fixed and random effects. σ^2^: Residual variance; τ_00_: variance for this random intercepts factor; ICC: intra-class correlation for this random intercepts factor; N: number of levels for this random intercepts factor.

*Quantile-Quantile Plot of Model Residuals*

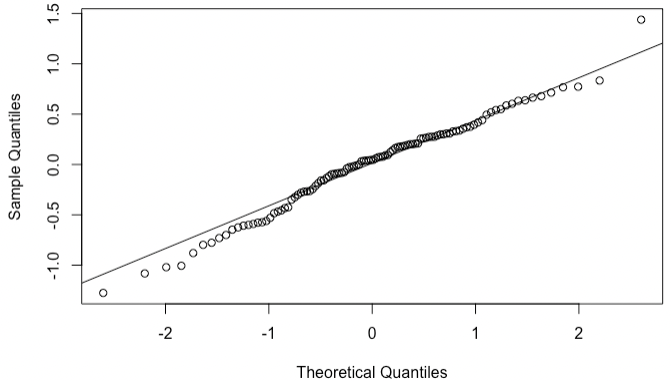

A Shapiro-Wilk test indicated that model residuals were normally distributed (*W* = .98; *p* = .22).

##### 8.2.7 Body Listening

*Linear Mixed Effects Model to Predict MAIA Body Listening Score*

| *Predictors* | *Estimates* | *std. Error* | *Statistic* | *p* | *df* |
| --- | --- | --- | --- | --- | --- |
| (Intercept) | 1.78 | 0.20 | 8.72 | **<0.001** | 60.83 |
| Group | -0.14 | 0.18 | -0.78 | 0.437 | 64.99 |
| Time [Final] | 0.25 | 0.13 | 1.96 | 0.055 | 52.00 |
| Age | 0.04 | 0.02 | 1.95 | 0.057 | 50.00 |
| Sex | 0.12 | 0.19 | 0.64 | 0.524 | 50.00 |
| Group × Time [Final] | 0.17 | 0.13 | 1.36 | 0.178 | 52.00 |
| **Random Effects** | | | | | |
| σ^2^ | 0.44 | | | | |
| τ_00_ _Participant_ | 1.15 | | | | |
| ICC | 0.72 | | | | |
| N _Participant_ | 54 | | | | |
| Observations | 108 | | | | |
| Marginal R^2^/ Conditional R^2^ | 0.076 / 0.745 | | | | |

**Legend.** Kenward-Roger approximation was used for degrees of freedom and *p*-values. The marginal R-squared accounts for the variance of the fixed effects only, while the conditional R-squared accounts for both the fixed and random effects. σ^2^: Residual variance; τ_00_: variance for this random intercepts factor; ICC: intra-class correlation for this random intercepts factor; N: number of levels for this random intercepts factor.

*Quantile-Quantile Plot of Model Residuals*

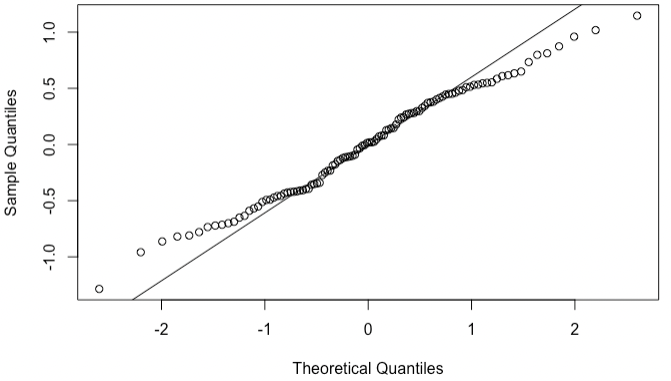

A Shapiro-Wilk test indicated that model residuals were normally distributed (*W* = .99; *p* = .36).

##### 8.2.8 Trusting

*Linear Mixed Effects Model to Predict MAIA Trusting Score*

| *Predictors* | *Estimates* | *std. Error* | *Statistic* | *p* | *df* |
| --- | --- | --- | --- | --- | --- |
| (Intercept) | 3.24 | 0.19 | 17.52 | **<0.001** | 66.63 |
| Group | -0.09 | 0.16 | -0.58 | 0.565 | 72.69 |
| Time [Final] | 0.10 | 0.14 | 0.72 | 0.476 | 52.00 |
| Age | 0.00 | 0.02 | 0.25 | 0.806 | 50.00 |
| Sex | 0.13 | 0.17 | 0.77 | 0.442 | 50.00 |
| Group × Time [Final] | -0.05 | 0.14 | -0.37 | 0.711 | 52.00 |
| **Random Effects** | | | | | |
| σ^2^ | 0.54 | | | | |
| τ_00_ _Participant_ | 0.81 | | | | |
| ICC | 0.60 | | | | |
| N _Participant_ | 54 | | | | |
| Observations | 108 | | | | |
| Marginal R^2^/ Conditional R^2^ | 0.024 / 0.610 | | | | |

**Legend.** Kenward-Roger approximation was used for degrees of freedom and *p*-values. The marginal R-squared accounts for the variance of the fixed effects only, while the conditional R-squared accounts for both the fixed and random effects. σ^2^: Residual variance; τ_00_: variance for this random intercepts factor; ICC: intra-class correlation for this random intercepts factor; N: number of levels for this random intercepts factor.

*Quantile-Quantile Plot of Model Residuals*

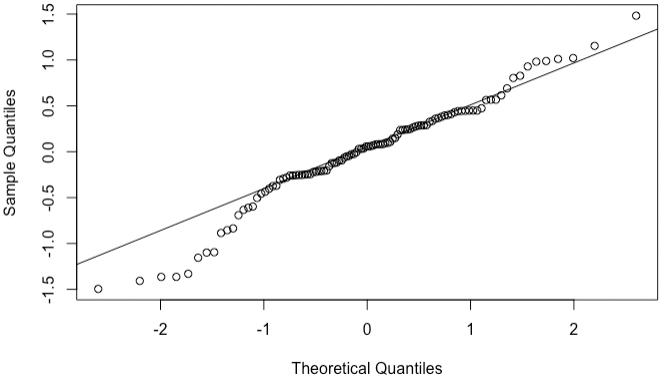

A Shapiro-Wilk test indicated that model residuals were not normally distributed (*W* = .97; *p* = .012).
